# Coherence of Terrestrial Vertebrate Species Richness with External Drivers Across Scales and Taxonomic Groups

**DOI:** 10.1101/2022.01.21.477239

**Authors:** Conor P. B. O’Malley, Gareth G. Roberts, Philip D. Mannion, Jan Hackel, Yanghua Wang

**Author notes:** **Corresponding Author:** Conor P. B. O’Malley, Royal School of Mines, Prince Consort Road, London SW7 2BP, UK. **Data Availability Statement:** A full working example analysis, with open-source (CC-BY-4.0) code is available at https://doi.org/10.5281/zenodo.7357231. Species richness, topographic and climate data used are publicly available and cited throughout. **Authorship Statement:** C.O. and G.R. designed the project, developed code, interpreted results and wrote the manuscript with contributions from P.M., Y.W., J.H.; C.O performed the analyses and generated figures.

## Abstract

**Aim:** Understanding connections between environment and biodiversity is crucial for conservation, identifying causes of ecosystem stress, and predicting population responses to changing environments. Explaining biodiversity requires an understanding of how species richness and environment co-vary across scales. Here, we identify scales and locations at which biodiversity is generated and correlates with environment.

**Location:** Full latitudinal range per continent.

**Time period:** Present-day.

**Major taxa studied:** Terrestrial vertebrates: all mammals, carnivorans, bats, songbirds, hummingbirds, amphibians.

**Methods:** We describe the use of wavelet power spectra, cross-power and coherence for identifying scale-dependent trends across Earth’s surface. Spectra reveal scale- and location-dependent coherence between species richness and topography (*E*), mean annual precipitation (*Pn*), temperature (*Tm*) and annual temperature range (∆*T*).

**Results:** *>* 97% of species richness of taxa studied is generated at large scales, i.e. wavelengths 10^3^ km, with 30–69% generated at scales 10^4^ km. At these scales, richness tends to be highly coherent and anti-correlated with *E* and ∆*T*, and positively correlated with *Pn* and *Tm*. Coherence between carnivoran richness and ∆*T* is low across scales, implying insensitivity to seasonal temperature variations. Conversely, amphibian richness is strongly anti-correlated with ∆*T* at large scales. At scales 10^3^ km, examined taxa, except carnivorans, show highest richness within the tropics. Terrestrial plateaux exhibit high coherence between carnivorans and *E* at scales *∼* 10^3^ km, consistent with contribution of large-scale tectonic processes to biodiversity. Results are similar across different continents and for global latitudinal averages. Spectral admittance permits derivation of rules-of-thumb relating long-wavelength environmental and species richness trends.

**Main conclusions:** Sensitivities of mammal, bird and amphibian populations to environment are highly scale-dependent. At large scales, carnivoran richness is largely independent of temperature and precipitation, whereas amphibian richness correlates strongly with precipitation and temperature, and anti-correlates with temperature range. These results pave the way for spectral-based calibration of models that predict biodiversity response to climate change scenarios.

## 1 Introduction

Biological diversity is critical to many basic human needs, including health, food, water and shelter. It also plays an important role in moderating physical and chemical processes in natural environments (Balmford & Bond, 2005; Barrett *et al*., 2011; Corenblit *et al*., 2011; Fei *et al*., 2014; Mori *et al*., 2022). Quantifying links between environment and biodiversity is crucial for understanding the response of ecosystems to climatic and physiographic change, and for conservation efforts (Araújo & Rahbek, 2006; Hampe & Petit, 2005; Norris *et al*., 2013; Yasuhara *et al*., 2020a). Many extrinsic processes postulated to control biodiversity (e.g. climate) are rapidly changing; therefore quantifying the strength of relationships between them is a pressing concern (Nogués-Bravo *et al*., 2018).

Environmental variables and species richness exhibit variance in space across a range of scales (e.g. Belmaker & Jetz, 2011; Buckley *et al*., 2012; Keil & Chase, 2019). However, it is unclear whether coherence between richness and climate is uniform across all scales, or whether climate is actually a significant driver of biodiversity across local, regional and global scales (e.g. Storch *et al*., 2007). Identifying links between biodiversity and environment across scales has recently become significantly more tractable for three reasons. First, global patterns of species richness have been estimated with unprecedented detail, from horizontal scales as broad as continents, to those as fine as *∼* 10 km in wavelength (e.g. Jenkins & Joppa, 2009; Jenkins *et al*., 2013, 2020; Kass *et al*., 2022; Marsh *et al*., 2022). Second, values and variance of many environmental variables postulated to be responsible for determining distributions of species are now available globally at even higher resolution (e.g. Karger *et al*., 2017). Finally, wavelet spectral methods, which can identify the locations and scales at which signals (e.g. spatial series of taxa) are generated, as well as coherence and phase differences (offsets) between series such as species richness, topography and climate, are now well-established (see Materials and Methods; Grinsted *et al*., 2004; Torrence & Compo, 1998). Such approaches are key to understanding how the changing global climate will affect the distribution of biodiversity across Earth.

In this study we demonstrate how information about scale- and location-dependent forcing of biodiversity trends can be used to test hypothesized drivers of richness (e.g. Belmaker & Jetz, 2011; Chase & Knight, 2013; Scheiner *et al*., 2000; Suárez-Castro *et al*., 2022). We assess whether one-dimensional signal-processing techniques are appropriate for revealing interdependence across scales, as has been suggested by some previous studies (Carl & Kühn, 2008; Ding & Ma, 2021). We focus on using wavelet transforms, which have a number of benefits over more widely known Fourier-based techniques. Their principal advantage is that they can be used to reveal the scales and positions in space (i.e. latitude) where biodiversity signals are generated, and their correlation with environmental variables. We demonstrate how calculated coherence can be used to test hypothesized drivers of species richness at scales ranging from 10s of km to *>* 10, 000 km (e.g. watersheds to continents). Coherence between species richness and environmental variables is calculated for endothermic and ectothermic non-marine vertebrate taxa across scales. Statistically significant coherence, beyond what is expected by random auto-correlation, is calculated. We assess whether scale-dependence of species richness and environmental variables results in scale-dependent coherence. Furthermore, we evaluate whether scale-dependence ought to be explicitly considered when examining ecological correlations and produce a framework to test predictions of species richness generated from environmental data.

The first hypothesis we test is that species richness is highly coherent, i.e. varies in concert, with environmental variables evenly across all scales. That would imply a direct forcing of richness by external drivers regardless of scale. It would give a basis for using theory developed to predict species richness at one scale (e.g. field sites) to predict richness at all scales. If confirmed, it would, for example, support the suggestion that environmental heterogeneity (e.g. elevation range, annual temperature range, etc.) acts as a primary driver of species richness regardless of scale (e.g. Stein *et al*., 2014). The second hypothesis tested is that species richness is most coherent with external variability at small scales, i.e. local changes in environment determine where species prosper, and therefore we must have a good understanding of processes operating at short scales to make reliable macroecological predictions. For example, Belmaker & Jetz (2012) find that local “environmental representativeness” with respect to regional climate may exert a strong control on local biodiversity. That result implies that regional environmental conditions are less important in controlling likelihood of amphibian, bird and mammal species occurrences. Their study, and others concerning a range of plant and animal taxa, imply that species richness and environment can be strongly coupled at short wavelengths in certain settings (Chase *et al*., 2019; Deák *et al*., 2021; Gutiérrez Illán *et al*., 2010; Mancera *et al*., 2005; Pincebourde *et al*., 2016; van Rensburg *et al*., 2002). Thirdly, we test whether species richness is most coherent with changes in environment at large scales, i.e. global scale variability (e.g. large-scale climate change), and that richness patterns have greatest variability at such scales, as has been suggested for several plant and animal families (e.g. Belmaker & Jetz, 2011; Field *et al*., 2009; Wang *et al*., 2009; Keil *et al*., 2012).

These three hypotheses concern the fundamental scale-dependence of interactions between terrestrial vertebrate richness and environment. They are usually tested via meta-analysis of studies carried out at different scales, be it extent, focus or grain, and not in the self-contained way which we present here (Scheiner *et al*., 2000). Field evidence and modelling suggests that richness relationships can follow each of the trends described above, depending on taxonomic group and habitat, and debate continues on whether there even exists a prevailing relationship between environment and richness across scales (e.g. Chase *et al*., 2019). Therefore, the fourth hypothesis we test is whether coherence of species richness with external variables depends on taxon. In other words, whether taxa have unique responses to environmental variables. This difference has been observed for some major terrestrial vertebrate and plant groups (see above; Boone & Krohn, 2000). Meta-analysis by Field *et al*. (2009) suggests that at least some species richness-environment interactions are consistent across different plant and animal taxa. Finally, we test the hypothesis that species richness does not directly depend on environment at any scale, or for any taxonomic group (i.e. coherence between species richness and environmental variables is generally/universally low). Instead, species richness depends upon other factors, namely biotic interactions (prey-predator, competition), and/or historical contingencies (e.g. Hagen *et al*., 2021; Ockendon *et al*., 2014; Sandom *et al*., 2013).

Here, we test each hypothesis by quantifying coherence between species richness of continental vertebrate taxa and elevation, precipitation, temperature, and annual temperature range, which are postulated to drive biodiversity (e.g. Antonelli *et al*., 2018; Rahbek & Graves, 2001). As a starting point, we focus on mapping coherence between contemporary biotic and environmental signals as a function of scale and location, using wavelet spectral analyses. Several approaches, e.g. spatial regression analyses, have previously been used to test such hypotheses (e.g. Butler *et al*., 2017; Chase *et al*., 2019). However, such methods require care when disentangling scale and location from biotic and environmental data to identify correlations (e.g. Sandel, 2015). Instead, here, we use wavelet spectral analyses, which inherently disentangle scale-dependent effects, and identify strength of correlation between variables at individual scales. Such analyses have been used to identify scale-dependence of temporal biotic series, to filter spatial series and identify outliers, and to investigate biodiversity on local or regional (500 km) scales, but not to investigate global trends or latitudinal gradients (Carl *et al*., 2008; Carl & Kühn, 2008; Carl *et al*., 2016; Dormann *et al*., 2007; Keitt, 2007; Ma & Zhang, 2015; Roberts & Mannion, 2019). We acknowledge that other processes, including species-species interactions, are also important for determining species richness (e.g. Chaudhary *et al*., 2021; Yasuhara *et al*., 2020b; Yasuhara & Deutsch, 2022). As such, we also present a preliminary assessment of the coherence between species richness of different taxa in Supporting Information. We return to discuss the results of these tests in the context of the five hypotheses described above, throughout the manuscript and in its concluding sections.

## 2 Materials and Methods

Several advantages exist to using wavelet analysis to tackle the hypotheses laid out in our Introduction. First, wavelet transforms are ideal for identifying scale- and location-dependent information in spatial signals, and make no requirement for signal stationarity (e.g. signals do not need to start and end at the same value). Secondly, they represent a self-contained means for investigation of scale-dependent effects from a given measurement, and do not rely on meta-analysis. Thirdly, calculated cross power and coherence can be used to identify respective locations of common high power, and commonalities between signals regardless of power, in distance-scale space. Those metrics provide a consistent framework for inter-comparison of many different taxa and environmental variables. Therefore, here we apply wavelet analysis to species richness and environmental data to evaluate the hypotheses laid out in the Introduction.

### 2.1 Species Richness Data

Species richness is here defined as the number of species of a given taxon within a 10*×*10 km square. We examine species richness trends in this study, since it is a straightforward measure of diversity, and has been determined for a wide range of taxa from fine scales up to near global scales. Here, we focus on terrestrial vertebrate taxa since terrestrial surface environmental conditions are well-mapped, as is terrestrial vertebrate biodiversity. Similar analysis is possible for marine taxa, invertebrates, plants etc., and for metrics other than species richness, for example range sizes and trophic interactions. We use richness grids compiled by Jenkins *et al*. (2013), which were generated by combining maps of species distributions, and counting the number of overlapping polygons in a given cell. For birds, the species richness data were calculated from breeding ranges compiled by BirdLife International NatureServe (2011). For amphibians and mammals, the data were based on expert range maps generated by the International Union for Conservation of Nature (2021).

Figure 1a–f shows species richness per 10*×*10 km cell for all mammals (Mammalia), carnivorans (Carnivora), bats (Chiroptera), songbirds (Passeriformes), hummingbirds (Trochilidae), and amphibians (Amphibia). In Supporting Information Figures 1 and 35, and Table 1, we show species richness and associated analyses for other evaluated taxonomic subgroups (including Cetartiodactyla, Eulipotyphla, primates, marsupials, rodents, parrots, and frogs). In Supporting Information Figure 2 we show results for species-species interactions for some of those taxa. The taxa which we focus on in the main text of this study have the greatest latitudinal coverage and are well-mapped, and they also cover a range of modes of life. The data shown in Figure 1 reinforce well-known large-scale observations, e.g. the latitudinal diversity gradient (Hillebrand, 2004; Willig *et al*., 2003), but also contain evidence of significant complexity across scales of interest, here wavelengths between 10–10^4^ km.

**Figure 1:**
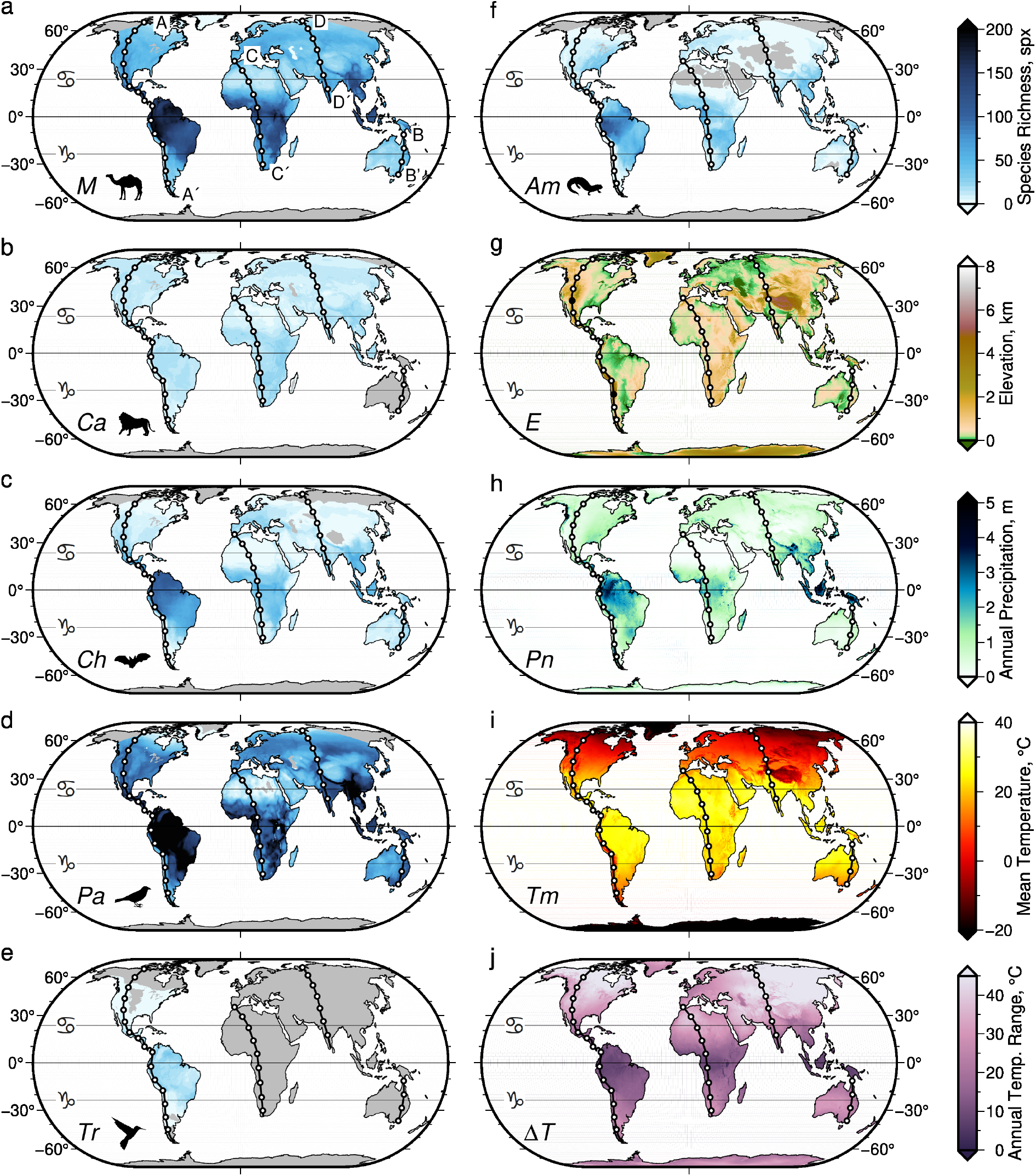
Global patterns of species richness and environment. (a) All Mammalia (*M*, mammals), (b) Carnivora (*Ca*, carnivorans), (c) Chiroptera (*Ch*, bats), (d) Passeriformes (*Pa*, songbirds), (e) Trochilidae (*Tr*, hummingbirds), (f) Amphibia (*Am*, amphibians); spx = species per 10 10 km pixel (Jenkins *et al*., 2013); horizontal lines = Tropics of Cancer (northern), Capricorn (southern), and Equator; A—A^*l*^ = transect through Americas investigated here; B—B^*l*^, C—C^*l*^, D–D^*l*^ = transects investigated in Supporting Information. Global latitudinal mean transects are also studied therein. (g) Elevation (*E*) from ETOPO1 global model with horizontal resolution of 1 arc-minute (Amante & Eakins, 2009); filled circles on A—A^*l*^ = Colorado Plateau/Mexican Highlands and Andean Altiplano. (h)–(j) Mean annual precipitation rate (*Pn*), temperature (*Tm*), and temperature range (∆*T*) from 1981–2010 (Karger *et al*., 2017).

**Figure 2:**
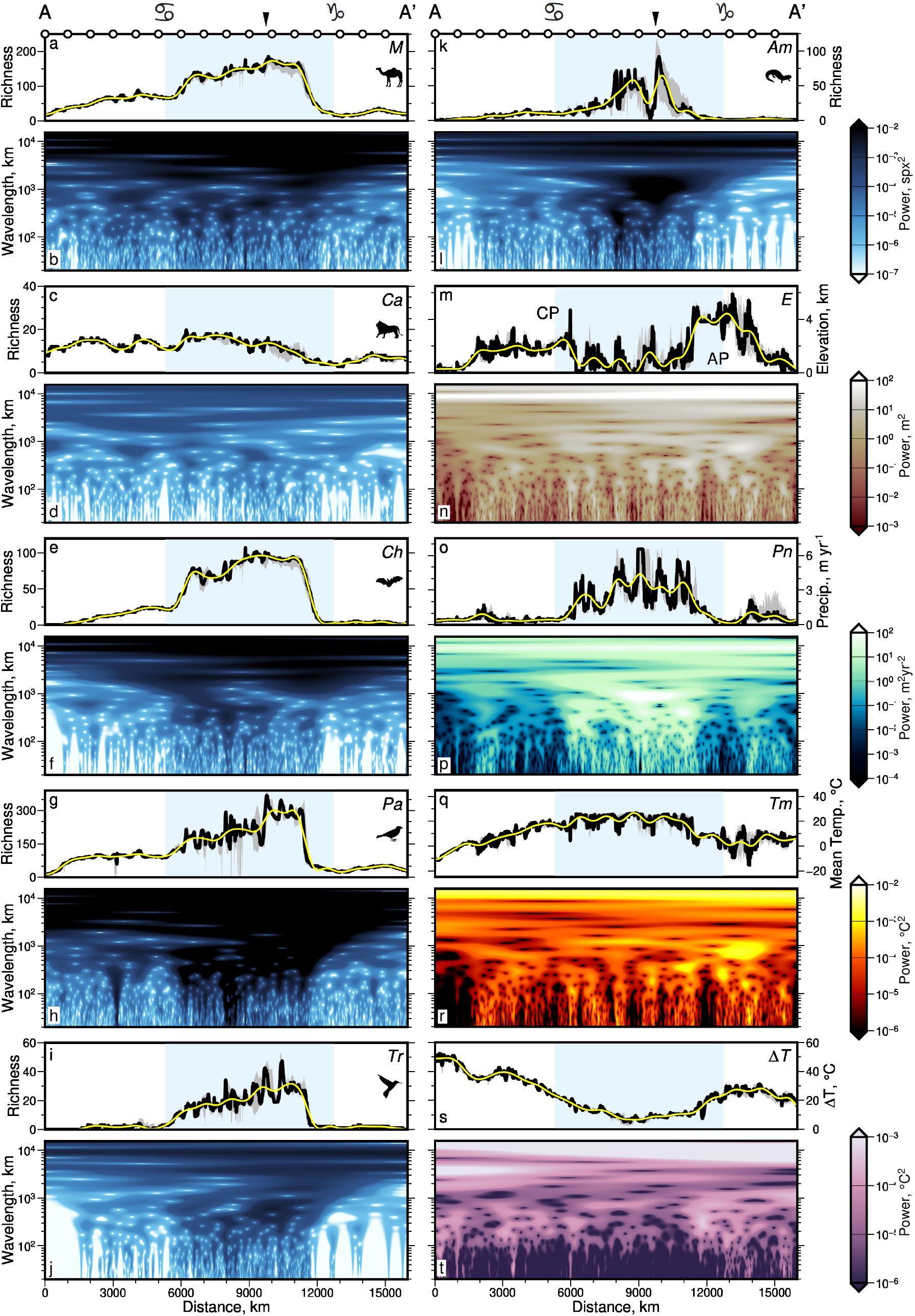
Wavelet transforms of species richness and environment. (a) Black line = species richness of Mammalia (*M*) along transect A—A^*l*^; gray bands = 100 km wide swaths centred on A—A^*l*^; blue bands = tropical latitudes; white circles are shown every 1000 km, see transect A—A^*l*^ in Figure 1; black arrow and symbols above top axis = Equator and tropics as in Figure 1. Yellow line = inverse wavelet transform of signal, filtering to pass only wavelengths *>* 1000 km; mean difference to input signal = 3.0 *±* 3.3 (1*σ*) spx. (b) Continuous wavelet transform of mammal richness spatial series (black line in panel a). Colors = rectified spectral power as a function of location and scale (wavelength); spx = species per pixel. (c)–(t) As (a)–(b) but for Carnivora (*Ca*), Chiroptera (*Ch*), Passeriformes (*Pa*), Trochilidae (*Tr*), Amphibia (*Am*), elevation (*E*), mean annual precipitation rate (*Pn*), temperature (*Tm*) and temperature range (∆*T*) along transect A—A^*l*^ (Amante & Eakins, 2009; Jenkins *et al*., 2013; Karger *et al*., 2017). Mean differences between signals and inverse transforms filtered to remove wavelengths *<* 1000 km = 0.7 *±* 0.6 spx (*Ca*), 1.5 *±* 2.1 spx (*Ch*), 11.6 *±* 16.7 spx (*Pa*), 1.5 *±* 2.5 spx (*Tr*), 2.9 *±* 5.3 spx (*Am*), 0.36 *±* 0.3 km (*E*), 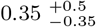, 2.2 *±* 2.2 C (*Tm*), and 1.2 *±* 1.1 C (∆*T*). See Supporting Information for results for transects B—B, C—C, D—D^*l*^ and average global latitudinal transect. Note high spectral power concentrated at wavelengths *>* 10^3^ km for all series. High species richness power (darker patches) at shorter wavelengths tends to be concentrated within the tropics.

Species richness varies as a function of the spatial range characteristics of a study, particularly “grain”, i.e. piece-wise horizontal resolution within a study (Gaston, 2000; Palmer & White, 1994; Willig *et al*., 2003). By using a constant grain (i.e. “focus” or grid spacing) of 10 km, challenges associated with comparing results generated using different grains are avoided (Willig *et al*., 2003). Here, scale-dependent trends are calculated as a function of “extent”, i.e. total width of study region, rather than “grain”, i.e. width of each plot/grid cell within the study region *sensu* Palmer & White (1994). Hurlbert & Jetz (2007) indicated that range map data might only be valid at wavelengths *>* 100 km. In this study, we evaluate how short-wavelength uncertainties in species richness contribute to uncertainties in calculated wavelet spectra by adding theoretical noise to transects before they are transformed into the spectral domain (Supporting Information Figure 3; panels a–c show results for several tests where large-amplitude white noise which contains wavelengths between 10–10,000 km is added to signals before analysis). Latitudinal transects through terrestrial vertebrate richness and environmental data are shown in Figure 2. We show data from the Americas, where transects can be generated that encompass almost all of Earth ‘s latitudinal range (Figures 1 & 2: A—A′). Transects through data for Australia (B—B′), Africa (C—C′), Eurasia (D—D′) and global averages are shown in Supporting Information Figures 4–7, and results of their wavelet analysis are summarized in Figure 4 of this document. In our choices of transect, we attempted to keep longitudinal variation to a minimum in order to better identify latitudinal trends. We note that A—A′, which we focus on here, intersects both topographic lowlands and highlands in North, Central and South America. Results for amphibian richness and environmental variables along alternative transects across the Americas, which do not follow lines of longitude as closely as A—A′, are shown in Supporting Information Figures 36–41.

**Figure 3:**
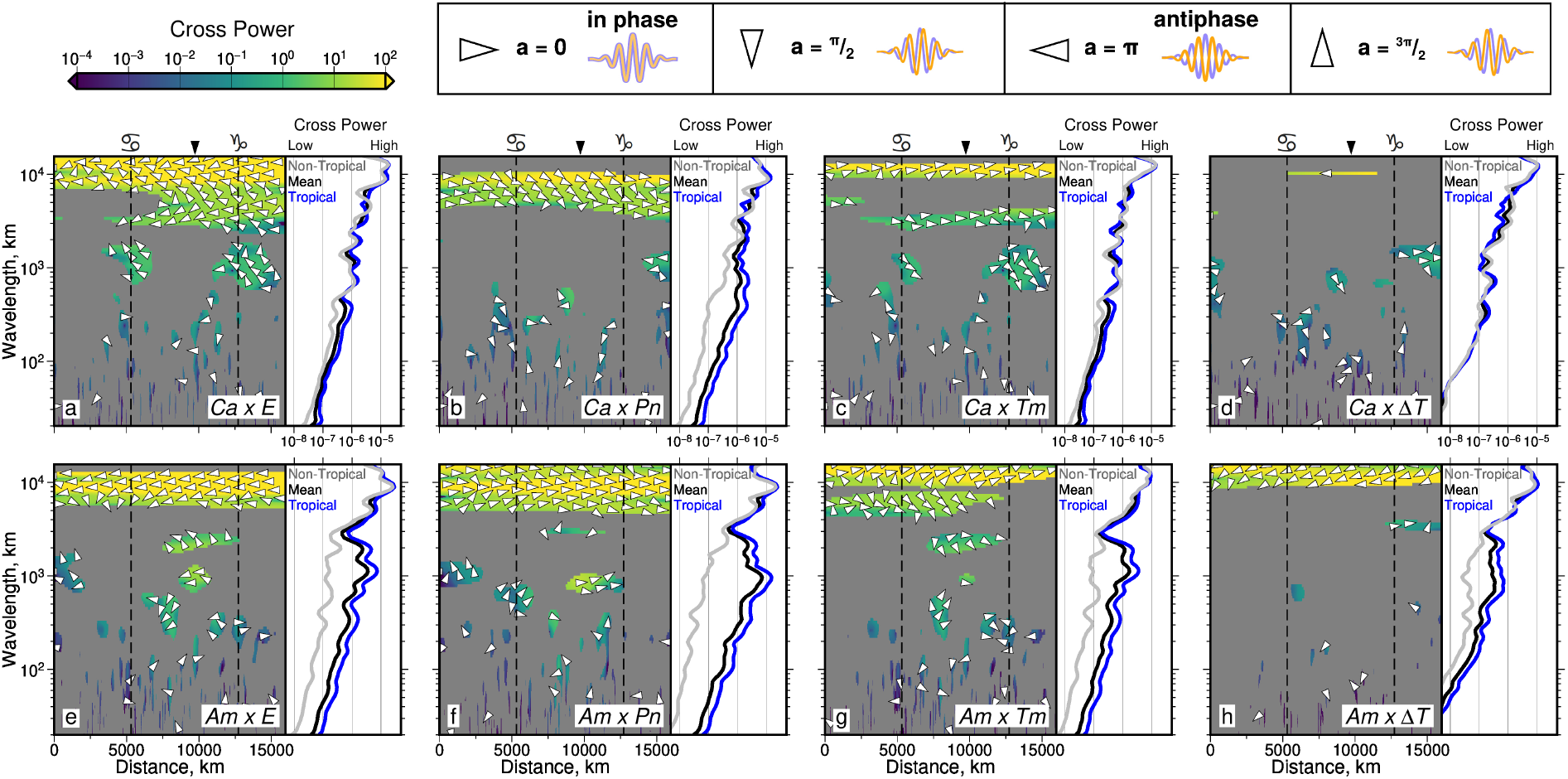
Coherence, cross power, and phase difference between species richness and environmental variables. (a) Comparison of Carnivora (*Ca*) and elevation (*E*) as a function of location and scale along transect A—A^*l*^ (Figures 1–2). Colors = cross wavelet power; yellow = co-located large (positive or negative) amplitude signals. Gray masks regions with coherence below 90% significance level (see body text, Materials and Methods). Arrows = phase difference between spatial series: right/left pointing = in-phase/antiphase (see guide above panels b–d). Black arrow and symbols above plot = Equator and tropics, as in Figure 1. Side panel: black/blue/gray lines = distance-averaged cross wavelet power of all/tropical/non-tropical latitudes (see Figure 2). High cross power = large co-located amplitudes in the two spatial series. (b)–(d) Comparison of Carnivora and mean annual precipitation rate (*Pn*), temperature (*Tm*) and annual temperature range (∆*T*). (e)–(h) Comparison of amphibian species richness and same environmental variables as panels a–d. Statistically significant coherence is concentrated at wavelengths *>* 10^3^ km, where species tend to be in- or anti-phase with environmental variables. The least statistically significant coherence is for Carnivora and temperature range (note gray mask across most of panel d).

**Figure 4:**
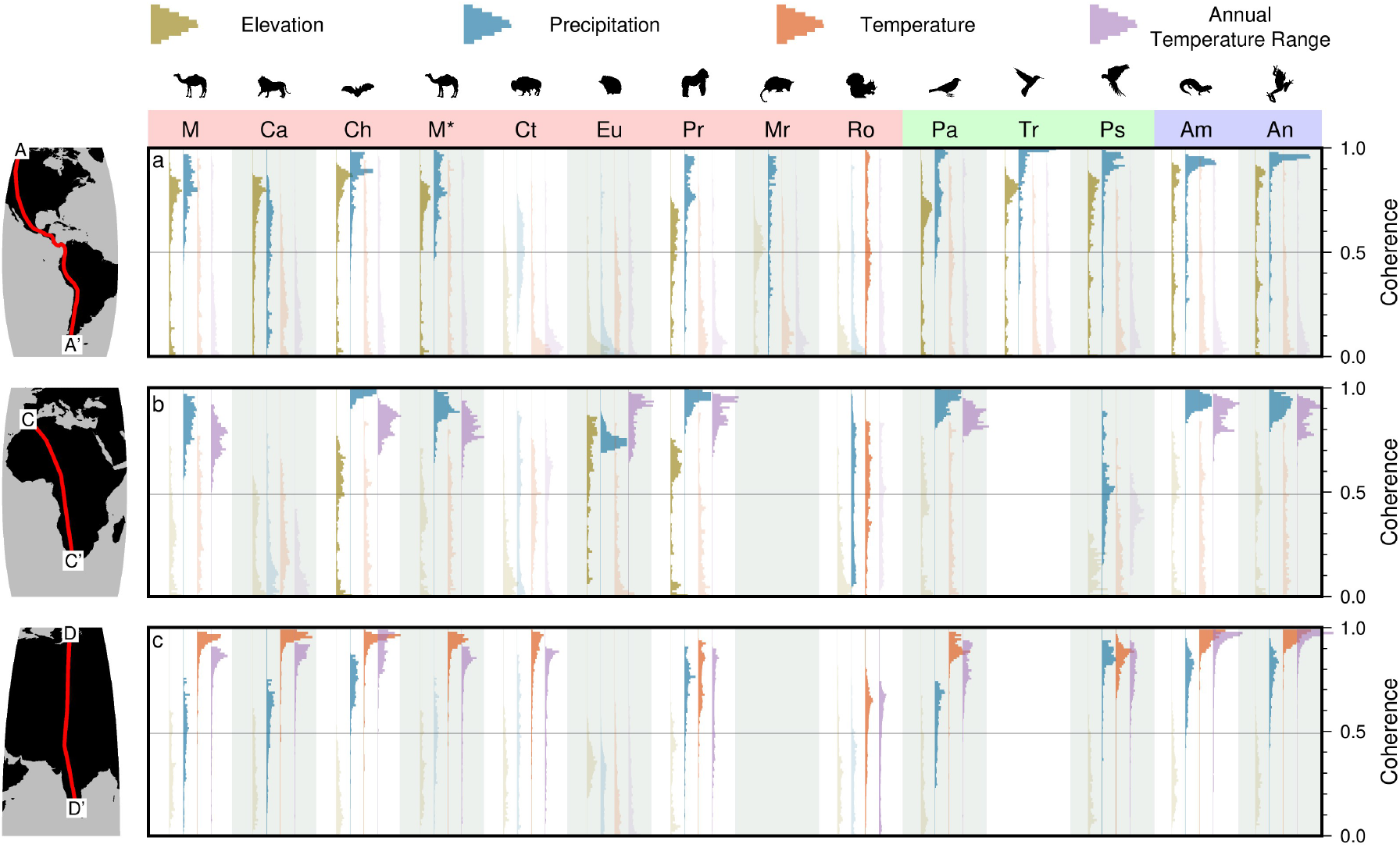
Summary of coherence between species richness and environmental variables at large scales. (a) Histograms = distribution of coherence (see right-hand *y* axis) between species richness and climate, for scales ≥3756 km (largest 25% of scales) and different taxa (see top *x* axis), across the Americas (transect A—A^*l*^). Brown/blue/red/purple histograms = coherence between species richness and elevation/mean annual precipitation/mean annual temperature/annual temperature range, respectively (see guide above plot; Jenkins *et al*., 2013; Karger *et al*., 2017). *M* = Mammalia, *Ca* = Carnivora, *Ch* = Chiroptera, *M*^*\**^ = Mammlia excluding Carnivora, *Ct* = Cetartiodactyla, *Eu* = Eulipotyphla, *Pr* = Primates, *Mr* = Marsupialia, *Ro* = Rodentia, *Pa* = Passeriformes, *Tr* = Trochilidae, *Ps* = Psittaciformes, *Am* = Amphibia, *An* = Anura. Pink/green/blue bands behind labels cover mammal/bird/amphibian groups, respectively. (b)–(c) As for panel (a) but for transects across Africa (C—C^*l*^) and Eurasia (D—D^*l*^), respectively—see adjacent maps and Figure 1 for transect locations.

### >2.2Environmental Variable Data

Figures 1g–j and 2m, o, q and s show examples of maps and cross sections through elevation and climatic data which we use, from the ETOPO1 and CHELSA datasets, respectively (Amante & Eakins, 2009; Karger *et al*., 2017). The global elevation grid ETOPO1 has a horizontal resolution of 1 arc-minute (Figure 1g; Amante & Eakins, 2009). It is primarily generated from *∼* 30 m resolution Shuttle Radar Topography Mission (SRTM30) data and includes interpolated coastlines and satellite altimetry (Jarvis *et al*., 2008). Amante & Eakins (2009) suggested a mean vertical error of *∼* 10 metres for ETOPO1. We down-sampled the data to a horizontal resolution of 10 km using Generic Mapping Tools to match resolution of species richness grids (Wessel *et al*., 2019). Annual mean values for climatic data, from 1981–2010, were extracted from the Climatologies at High Resolution for the Earth ‘s Land Surface Areas (CHELSA) dataset (Karger *et al*., 2017). CHELSA was generated by applying corrections to the ERA-Interim climatic reanalysis and has a horizontal resolution of up to 30 arc-seconds (Dee *et al*., 2011). Temperature data were corrected for elevation above sea level and precipitation rates were corrected using wind direction, valley exposition and boundary layers. Precipitation rate is weakly dependent on elevation. These values were successfully benchmarked against alternative climatology data and models: WorldClim, TRMM, GPCC and GHCN (Hijmans *et al*., 2005; Goddard Earth Sciences Data and Information Services Center, 2017; Lawrimore *et al*., 2011; Schneider *et al*., 2014). The data were down-sampled to 10 km prior to spectral analyses, a limit imposed by the 10*×*10 km resolution of species richness maps (Jenkins *et al*., 2013).

### 2.3 Continuous Wavelet Transform

Spatial series, *x*_*n*_, of species richness or environmental variables were transformed into distance-wavenumber space using continuous wavelet transforms (for practical guide, see Torrence & Compo, 1998). The transform convolves uniformly sampled spatial series with a mother wavelet, *ψ*. Since wavelet transformation reveals power as a function of both wavenumber (here spatial scale— but with worse frequency resolution than an equivalent Fourier transform), and position (here distance along each transect), the resulting continuous wavelet transform is a map colored by the degree to which the input signal contains high amplitude variability at given positions and scales (see Figure 2). The Morlet wavelet with dimensionless frequency *ω*_*◦*_ = 6 is used in this study, although other mother wavelets are investigated in Supporting Information Figure 8. Use of different mother wavelets (Morlet, order *ω*_*◦*_ = 4, 8; Paul, order *m* = 2, 4, 6; derivative of Gaussian, order *m* = 2, 4, 6) does not significantly change patterns of mapped power, and distance-averaged power shows similar trends to the results presented here. The mother wavelet is scaled and translated along spatial series by *n*′ to reveal variations in amplitude as a function of scale, *s*, and position, *x*_*n*_. Sampling interval *δt* = 10 km, *n* = 0, 1 … *N −* 1, where *N* is number of measurements. The wavelet transformation is given by

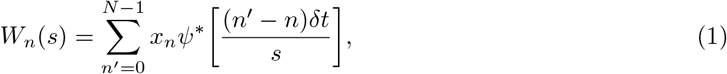

where *\** denotes the complex conjugate. We use the PyCWT Python module to transform the spatial series (Krieger *et al*., 2020), which is based on the methods summarized by Torrence & Compo (1998). Scales were calculated using the approach described in Torrence & Compo (1998), such that 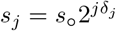, where *j* = 0, 1, … *J*. The smallest scale, *s*_*◦*_ = 2*δt*. A minimum grid spacing of 10 km therefore yields a minimum scale for wavelet spectral analysis of *∼* 20 km (Torrence & Compo, 1998). In the example A—A′ shown in Figure 2, *N* = 1598, *δ*_*j*_ = 0.1 and *J* = 96, which yields a total of 97 scales that range from *∼* 20 to *∼* 15, 521 km. Spatial series were mirrored across the *x* (distance) and *y* (dependent variable) axes to reduce edge effects (Roberts *et al*., 2019). If signals are not mirrored, edge effects will be included in power spectra and subsequent coherence analysis, which we wish to avoid. In Supporting Information Figures 34–35 we demonstrate how results differ if mirroring is not applied to signals. Inverse transforms were reconstructed for each signal to quantify fidelity of transformed series (see Torrence & Compo, 1998). Median difference between input signals and inverse transforms were always *≤* 0.9%. Signals can be filtered by calculating inverse transforms at specific wavelengths. For example, Figure 2a, c, e, g, i and k show inverse transforms of species richness spectra using only components of the signal of scales *>* 10^3^ km, across the whole length of each transect. Depending on taxonomic group, these filtered signals fit the input species richness trends with mean differences of 4.4–25% (see Figure 2 caption for mean raw differences in terms of species per pixel, spx). The same filtering process, but including wavelengths *>* 10^2^ km, yields mean differences of only 0.7–4.7%.

In Figure 2 we plot rectified wavelet power, |*W*_*n*_(*s*)|^2^*s*^*−*1^ as a function of location and scale. Rectification of spectra in this way is necessary to ensure wavelet power spectra and associated slopes are comparable with equivalent Fourier power spectra (Liu *et al*., 2007). The distance-averaged power spectrum is given by

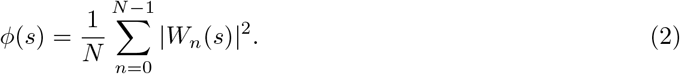

In subsequent analyses we use rectified power and rectified distance-averaged power *φ*_*r*_ = *φ*(*s*)*s*^*−*1^, which yields similar results to Fourier transformation (see e.g. Supporting Information Figures 9– 23). We calculate distance-averaged power within and outside of the tropics, but note that in those calculations, power was normalized by the proportion of the transect within/outside of the tropics respectively (Supporting Information Figures 19–23). Therefore there is no bias in the distance-averaged power shown if the transect has a greater distance within/outside of tropical latitudes. A guide to scale-dependence and self-similarity of spatial series is the color of spectral noise that they possess. For example, red (Brownian) noise occurs when *φ*_*r*_ *∝ k*^*−*2^, where *k* is wavenumber or spatial frequency, proportional to 1*/*wavelength, indicating self-similarity. Pink noise occurs when *φ*_*r*_ *∝ k*^*−*1^, and white noise indicates that power is equal across all scales, *φ*_*r*_ *∝* 1. Best-fitting spectral slopes for all variables and transects were identified using simple one- and two-slope models after Roberts *et al*. (2019); see Supporting Information Figures 9–18.

### 2.4 Cross Wavelet Power & Wavelet Coherence

Cross wavelet power is calculated to identify signals in separate spatial series (e.g. amphibian richness and precipitation) that have large amplitudes located at the same position in distance-wavenumber space. To facilitate comparison, signals are normalized to zero mean and unit variance prior to transformation. The normalized signals *X* and *Y*, are transformed to yield *W*^*X*^ and *W*^*Y*^. Cross wavelet power *W*^*XY*^ is calculated such that

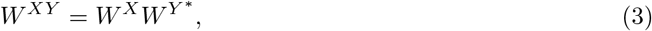

where *\** denotes complex conjugation. Wavelet coherence, 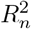, is calculated to identify parts of signals that are coherent, but not necessarily of common high amplitude, such that

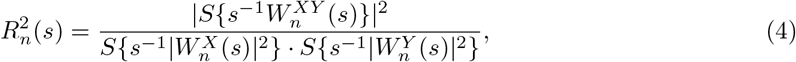

where *s, n* and *W*_*n*_(*s*) are as in Equation 1. *S* is an operator that smooths along distance and scale (Grinsted *et al*., 2004). Note that for the coherence calculation, signals are not normalized before cross power is calculated, since the denominator of 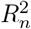 contains the power of *X* and *Y*.

Signals with certain spectral distributions (e.g. red noise) can, by chance, correlate without true interdependence. Therefore, it is important to calculate the coherence between each pair of signals, and not simply their cross wavelet power. Torrence & Compo (1998), Grinsted *et al*. (2004) and others have shown that coherence between signals above (assumed) background noise (i.e. spectral distributions) can be estimated by first calculating the coherence between large numbers of surrogate datasets with the same auto-regressive (AR) coefficients as the original data set. In this study, the minimum bound for statistically significant coherence (above assumed background noise) per scale, for each transect, was calculated from cross wavelet power spectral analysis of 300 random signals. Guided by the spectral content of actual biotic and environmental signals, we assumed that each theoretical random signal has a red noise distribution, which was generated using the same auto-correlation coefficient as the actual input signals, as suggested by Cazelles *et al*. (2014). The surrogates also have the same length and number of measurements, *N*, as the actual signals. The 90% significance limit for coherence, which was used to mask Figure 3, depends only on scale and not position, and was calculated using Monte Carlo methods with the PyCWT Python module (Grinsted *et al*., 2004; Krieger *et al*., 2020).

The local phase difference (angular offset, 0 *≤ a ≤* 2*π*) of two signals is given by the complex argument of their cross wavelet transform, arg *W*^*XY*^ (Grinsted *et al*., 2004). Figure 3 indicates phase difference as arrows measured from horizontal: in-phase, *a* = 0, t>; anti-phase, *a* = *π*, <J. A working example for species richness and elevation, including continuous wavelet transformation, cross wavelet power and wavelet coherence calculations, can be found at doi.org/10.5281/zenodo.7357231 (O ‘Malley & Roberts, 2022).

## 3 Results and Discussion

### 3.1 Wavelet Transformation of Richness and Environmental Variables

Spectral analyses of vertebrate species richness and environmental variables are shown in Figures 2 and 3. Figure 2 shows that highest spectral power, *φ*_*r*_ (*∝ z*^2^, where *z* is signal amplitude), is concentrated at largest scales for all taxa and environmental variables studied. Dependent on taxonomic group, from 96% to almost 100% of power resides at wavelengths *>* 10^3^ km. 30–69% of power resides at wavelengths 10^4^ km. These results reinforce the notion that species richness is dominated by long wavelength, latitudinal, variability, i.e. the third hypothesis outlined in the Introduction.

In fact, species richness tends to have a pink noise spectrum (see Supporting Information Figures 9–18, where slopes are fitted to average power spectra formally, for each variable). Thereby, shorter wavelength features in species richness signals tend to have the lowest amplitudes and comprise relatively little (few %) of species richness signal at a particular location. Mammal and bat species richness is better characterized by red noise at long wavelengths. This result implies self-similarity across scales, and that signal amplitudes decrease even more rapidly with decreasing wavelength than for other taxa. Therefore, in agreement with the fourth hypothesis stated in the Introduction, there seems to be some dependence on taxonomic group when considering the connections between richness and environment at the scales we study. At wavelengths 10^3^ km, species richness power for amphibians is best characterized as blue noise, i.e. *φ*_*r*_ *∝ k*^1^. This trend is not observed along the entire transect, but indicates that short wavelength features can be increasingly important contributors to amphibian richness (see Figure 2f). However, a single spectral slope akin to pink noise can adequately fit the amphibian richness spectrum (see e.g. Supporting Information Figure 9f). To address the hypotheses posed in the Introduction, it is important to consider colors of spectral noise (i.e. spectral slopes/scaling regimes) of both species richness and environmental variables for three reasons. First, it is of general scientific interest to highlight scale-dependence of individual signals. Secondly, colors of noise of individual wavelet transforms directly control the distribution of cross wavelet power between the two signals. We wish to identify both regions in scale-position space of common high cross power and coherence. If coherence between two variables is high at certain positions/scales, then it is more likely that there is a causal/interdependent link between two variables. However, it cannot be said that the environmental variable is contributing to a large proportion of species richness trend unless cross power between the two is *also* high. Lastly, colors of noise are directly related to auto-regression coefficients, which are used to generate expected correlation between random signals to calculate coherence thresholds, and therefore to assess whether two variables are indeed significantly correlated (see Materials and Methods).

We address concerns regarding the accuracy of range maps in two ways. First, we show the results of inverse wavelet transforms at scales *>* 1000 km alongside the results for transformation of the full frequency content of the available data (Figure 2). Secondly, we explore the impact of inserting distributions of theoretical, hitherto unknown, species in a suite of increasing severe tests for our results and conclusions. White noise was added to the amphibian transect in a systematic set of tests. These tests examined changes in calculated spectra when noise with maximum amplitudes of 10%, 50% and 100% of the standard deviation of the original signal ‘s amplitude (in this case = 24 species per pixel) was added to the transect prior to transformation. These tests included adding noise containing wavelengths between 10 and 100, 1000 and 10,000 km (Supporting Information Figure 3). As expected, these tests indicated that spectral power is least likely to be precisely known at the shortest wavelengths. Nonetheless, they demonstrate that even high amplitude uniformly distributed noise does not significantly change the overall spectral characteristics of terrestrial species richness. Finally, we note that that Hurlbert & Jetz (2007) suggest that species richness values estimated from range maps are likely to be overestimates at short length scales compared to richness estimated from atlas data (e.g. their Figure 5). That result indicates that range maps are more likely to generate higher power at short wavelengths than maps derived from atlas data. Therefore, spectra derived from atlas data are likely to be even redder than those obtained from range maps. In other words, our conclusion that species richness is dominated by long wavelength structure is expected to be insensitive to the choice of range or atlas data. In fact, atlas data is likely to more strongly emphasize the importance of long wavelength variability for determining species richness. We suggest that these observations and results indicate that range maps are useful for our purposes, and they reinforce our conclusion that, for the taxa studied here, the third hypothesis outlined in our Introduction is supported. That is to say, long-wavelength trends dominate species richness patterns.

Although almost no power is concentrated below wavelengths of *∼* 100 km for any of the taxa examined here, there are regions of some wavelet transforms which show increased power in the range *∼* 300–1000 km. This deviation, away from a broadly monotonic decrease in power towards shorter wavelengths, is driven principally by species richness within tropical latitudes, and is especially prominent for songbirds, hummingbirds and amphibians (Figure 2h, j, l). Supporting Information Figure 19a–f shows that at wavelengths 1000 km, there is no notable difference between power in species richness within or outside the tropics, across the Americas. However, at wavelengths 1000 km, there is significantly greater power for regions within the tropics. This trend arises because power spectral slopes remain close to *−*2 at shorter wavelengths outside of the tropics (i.e. red noise; Supporting Information Figure 19), before increasing to be closer to *−*1 (i.e. pink noise). We suggest that these results are consistent with the concept that variations in topography within tropical regions can generate higher species richness variability towards the equator via the increased effectiveness of altitudinal variation in habitat at isolating species either physically or physiologically (i.e. by being associated with variation in other environmental variables such as temperature; Ghalambor *et al*., 2006; Janzen, 1967). Figure 3 reveals that within the tropics, low elevations may be associated with increased species richness, which we discuss in Section 3.2. We find that this effect has the greatest impact on species richness power of hummingbirds and amphibians; the impact on bat and songbird richness appears to be more modest. Tropical increases in species richness of carnivorans, and mammals more generally, are much more subdued (Supporting Information Figure 19). Therefore we demonstrate different scale-dependence of species richness trends for different taxonomic groups, supporting the fourth hypothesis outlined in the Introduction. Although the spectral “flattening” from red to pink noise signals at scales *≈* 1000 km suggests there is increased short-wavelength variability in tropical hummingbirds and amphibians compared with other taxa, a pink noise spectral slope still suggests that highest amplitude richness trends reside at the larger scales even within this scale range. So, results for all taxa support the third hypothesis from Section 1, i.e. that large-scale trends dominate richness patterns, to varying degrees.

Elevation transects exhibit red and pink noise spectral characteristics at wavelengths 10^3^ km and 10^3^ km, respectively, which we note is similar to distance-averaged power from wavelet transforms of longitudinal river profiles and other topographic transects (Supporting Information Figures 9g, 9q, 10g, 10q; Roberts *et al*., 2019; Wapenhans *et al*., 2021). Precipitation rate, temperature and annual temperature range can also be characterized as red and pink noise (Supporting Information Figures 9h–j, r–t & 10h–j, r–t). Similar results are obtained for latitudinal transects through Africa, Eurasia and Australia, as well as across global, latitudinally-averaged sections (see Supporting Information Figures 11–23).

### 3.2 Coherence between Richness and Environment

Visual inspection of Figure 2 indicates that there is strong, location- and scale-dependent, similarity between the wavelet transforms of transects through species richness and environmental variables. To quantify the strength of these relationships we calculate cross wavelet power, which identifies colocated high amplitudes in the location-scale domain. We also calculate wavelet coherence, the degree to which signals are correlated above an expected background amount that auto-correlated signals would correlate, at given locations and scales (see Materials and Methods). Figure 3 shows results for carnivorans (which are similar to those for mammals generally), and amphibians (which are similar to those for bats, songbirds and hummingbirds). See Supporting Information Figure 1, 35 and Table 1 for analyses of those other taxa. We think it is important to compare and contrast the coherence of taxa with different modes of life, i.e mammals/carnivorans versus amphibians, with environment. It is useful and interesting to assess whether the long-theorized differences in environmental sensitivities of these taxa is borne out in our wavelet spectral analysis (e.g. Belmaker & Jetz, 2011). Figure 3a shows cross wavelet power between species richness of carnivorans along transect A—A′ and elevation. Almost no short-wavelength (*<* 10^3^ km) features are coherent above a 90% confidence limit. These short wavelength regions contain almost no cross wavelet power; 94% of all cross wavelet power is in the region of high coherence colored on Figure 3a, which accounts for 30% of the location-scale domain. 79% of the area of the cross wavelet spectrum that is significantly coherent resides at wavelengths 10^3^ km. Distance-averaged cross wavelet power for all parts of the power spectrum, not just those parts which are coherent above the 90% significance threshold, is shown to the right of each panel, on a logarithmic scale. Full, unmasked, plots of cross wavelet power are shown in Supporting Information Figure 24. Masked and unmasked cross power plots for other transects and global latitudinal averages are shown in Supporting Information Figures 25–32. Distance-averaged cross wavelet power between all taxa and environmental variables studied is shown in Supporting Information Figures 19–23, panels g–ad.

Cross wavelet power between amphibians and elevation is also highest at long wavelengths, although overall there is a smaller proportion of the two signals that is coherent: 78% of the plot region is masked in Figure 3e (gray regions). Only a small part of the cross wavelet transform for amphibians and elevation is coherent below wavelengths of *∼* 5000 km, and that part lies near the centre of the transect, i.e. within the tropics. Distance-averaged power outside the tropics, plotted to the right of Figure 3e as a gray curve, is an order of magnitude lower than within the tropics, especially at wavelengths 3000 km. This observation is in contrast to cross power between species richness of carnivorans and elevation, where there is almost no difference between the results within and outside of the tropics, across all scales. These results may indicate that carnivorans are less affected by “mountain passes” (*sensu* Janzen, 1967) in the tropics, compared with amphibians (cf. Antonelli *et al*., 2018; Eronen *et al*., 2015; Rahbek *et al*., 2019; Rolland *et al*., 2015). At wavelengths *∼* 10^3^ km, carnivoran species richness is most coherent with elevation and mean annual temperature atop terrestrial plateaux (e.g. Rocky-Mountains-Colorado Plateau and Altiplano, between 4000 *−* 7000 km and 13, 000 *−* 14, 000 km distance along transect A—A′, respectively; Figures 1–3). An obvious interpretation is the local importance of tectonics for determining biodiversity (Antonelli *et al*., 2018).

Statistically significant (above background red noise), coherent cross wavelet power between carnivoran species richness, mean annual precipitation rate, temperature and annual temperature range is shown in Figure 3b–d. Results for amphibians are shown in panels f–h. Cross power between amphibian species richness and precipitation rate, temperature, and temperature range tends to be higher within the tropics compared to outside the tropics at wavelengths 3000 km (cf. grey, blue, black lines in Figure 3). Those differences are absent or reduced for carnivorans. Furthermore, a smaller area of the power spectra of these three climatic variables is significantly coherent with carnivoran richness, compared to amphibian richness (cf. extent of gray masks in Figure 3). One likely interpretation of these results is that carnivorans are less sensitive to changes in those variables than amphibians (e.g. Rolland *et al*., 2018). Calculated phase indicates long-wavelength anti-correlation between elevation and species richness for both carnivorans and amphibians (left-pointing arrows in Figure 3a and e; phase angle, *a* = *π*; see Materials and Methods). Highly coherent long-wavelength anti-correlation between amphibian species richness and annual temperature range is also observed across the entire transect. Highly coherent, long-wavelength cross power between precipitation rate or temperature, and species richness of both carnivorans and amphibians, is in phase, i.e. there is positive correlation at these scales. This result is in agreement with the idea that faster diversification rates contribute to heightened species richness, since it suggests that both taxonomic groups benefit from increased energy and higher productivity associated with greater availability of heat and water (cf. Allen *et al*., 2006).

Since we observe that coherence and high cross wavelet power are both generally highest across taxa at long wavelengths (10^3^ km), in Figure 4 we plot histograms of coherence between richness and environment, above scales of 3756 km which represents the upper quartile of scales. We show results for each environmental variable and taxon previously mentioned, as well as for a range of groups whose species richness patterns were also compiled by Jenkins *et al*. (2013). Figure 4a shows results for the Americas i.e. transect A—A′. This figure summarizes our results in a way that can be directly related to the hypotheses posited in the Introduction. The background expected coherence value between random signals with the same auto-regression coefficient as each variable/species richness signal, at scales 3756 km is *≈* 0.5 and is shown in Figure 4. Results from Figure 3 are reflected in Figure 4a— note the cluster of high coherence values for amphibians and carnivorans and elevation, far above the 90% confidence level. Similarity of results for carnivorans with those for other mammals, including bats and primates, is evident, and the third hypothesis is likely true for those groups, i.e. large-scale trends dominate species richness and are strongly coherent with external variables, especially precipitation and elevation. However, groups such as Cetartiodactyla (even-toed ungulates & cetaceans), Eulipotyphla (hedgehogs, etc.) and rodents, show very low coherence at long wavelengths with any environmental variables. Therefore the third hypothesis is clearly not a generalized rule-of-thumb for all taxa, i.e. hypothesis four is likely to be true, namely that scale-dependent interactions between species richness and environment are taxon-dependent. Results for other transects are discussed below, and shown in Figure 4b–c.

### 3.3 Global and Local Species Richness and Environment

The results for the Americas can be compared to those for transects from Australia, Eurasia and Africa. For Australia, similar trends in power spectral slopes, distance-averaged power and cross wavelet power are observed (Figure 1: B—B′; Supporting Information Figures 4, 11–12, 20, 25–26). However, for Australia there is almost no difference in power or cross power between tropical regions and regions outside the tropics. We note, however, that the Australian transect does not include the entirety of the tropics; it only spans latitudes between 11.2^*◦*^S–37.6^*◦*^S. Signals are mostly coherent at wavelengths 10^3^ km, and the same pattern of correlation/anti-correlation is observed with climatic variables as that recovered for the Americas (Supporting Information Figures 25–26). Since no latitudinal transect of length *>* 3756 km is possible across Australia, filtered coherence results for B—B′ are not shown in Figure 4.

In Africa, songbirds and amphibians have greater species richness power within the tropics but the differences outside of the tropics are not as stark as for the Americas (Figure 1: C—C′; Supporting Information Figure 21a–f). This result may reflect differences in Cenozoic paleoclimatic history between Africa and the Americas (Hagen *et al*., 2021). The greatest difference between cross power within and outside the tropics is for precipitation rate, suggesting that water availability is a more important control species richness for all African taxa studied here. This result is reflected in Figure 4b, especially for mammals— histograms show that coherence between species richness of most taxa and annual precipitation is higher than for the Americas. However, wavelet coherence indicates that African carnivoran species richness does not correlate with environmental variables, whereas species richness of African amphibians is strongly positively correlated with precipitation rate at long wavelengths, in agreement with the findings of Buckley & Jetz (2007) and Buckley *et al*. (2012). Anti-correlation is observed between amphibian species richness and temperature across Africa.

Results for Eurasia are dominated by the presence of the Tibetan Plateau, and the low proportion of the transect within tropical latitudes, making it difficult to draw conclusions about latitudinal diversity gradients (Figure 1: D—D′; Supporting Information Figures 6, 15–16, 22, 29–30). Some similar trends to the Americas are observed, albeit with generally lower cross power and coherence; in particular there is still high coherence between large-scale species richness and annual precipitation for most taxa. However, a persistent result which is specific to the Eurasian transect is that there is high coherence and positive correlation between richness of most taxa and mean annual temperature, and high coherence and negative correlation between richness and annual temperature range, in agreement with Luo *et al*. (2015) (Figure 4c). The increased correlation between richness and temperature is likely because of the extremely low mean temperatures across the Tibetan Plateau causing temperature to become a controlling variable on species survival.

Mean terrestrial values of each variable across all latitudes globally were transformed into the location-scale domain. Distance-averaged wavelet power spectra of the resulting transects have spectral slopes between *−*2 and *−*1 (red to pink noise), reflecting the importance of long-wavelength trends (Supporting Information Figures 17–18). Species richness power for all taxa, except Mammalia and Carnivora, is at least an order of magnitude lower outside of tropical latitudes, at wavelengths 3000 km, consistent with results obtained from transforming the American transect (Figures 2 and 3, Supporting Information Figure 23). This result suggests that the increase in species richness power at short wavelengths may be a global phenomenon, reflecting sensitivity of tropical species to local climatic effects. However, care should be taken when interpreting global results since where there is high coherence, transects may not necessarily have high power at the same longitudes, just at common latitudes.

Supporting Information Figure 33a–f shows a summary of results for the transect across the Americas, A—A′. Panels a and b show inverse wavelet transforms generated using only the the longest 25% of scales (wavelengths *>* 3756 km; approximately one quarter of the length of the transect), for amphibian and carnivoran species richness respectively. Those taxa have significantly different modes of life and their wavelet power, cross power and coherence with environment exhibit the greatest differences of any taxa studied, hence we also present cross power and coherence for those taxa in Figure 3. The low-pass filtered series account for *∼* 41 and *∼* 11% of species richness respectively, in terms of mean difference to input signals. Supporting Information Figure 33c–f shows inverse wavelet transforms of environmental series filtered in the same way. Coherence, 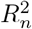, between the filtered series and amphibian and carnivoran species richness trends is annotated on each panel. These results for the Americas are consistent with global averages, although coherence is generally lower for global results, which is perhaps to be expected since regions of common power may not be sited at the same longitudes (see Figure 4).

### 3.4 Transfer Functions Between Environment and Species Richness

It is useful to derive ‘rules-of-thumb ‘ to estimate species richness from environmental variables. If several variables exhibit high coherence with richness of a given taxon, spectral admittance values may indicate which is likely to contribute to a greater proportion of the variability in richness. Once the hypotheses outlined in our Introduction have been supported or rejected, admittance values between variables can be investigated at the appropriate scales. Since this study found general support for hypotheses three and four, i.e. richness trends are likely generated by large-scale environmental patterns, conversion schemes (AKA transfer functions) are likely to be particularly useful at large scales, where most species richness appears to be determined and where coherence with environmental variables is highest. In the scale-distance domain, after wavelet transformation (see Methods and Materials), species richness *W* of any given taxon *X* can be expressed:

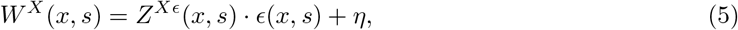

where *x* is distance, *s* is scale, and *η* is noise or contributions from variables that have not been considered. *Z*^*Xε*^ refers to the admittance (transfer function) between a set of environmental variables *E*, and richness in the scale-distance domain, *W*^*X*^. For any individual variable *Y* within *E*, the admittance between *W*^*X*^ and *W*^*Y*^ can be expressed:

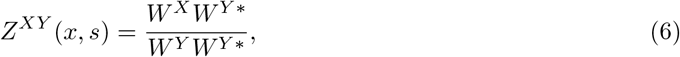

where ^*\**^ denotes complex conjugate. Thereby, a species richness signal *X* can be estimated from calibrated admittance *Z*^*XY*^, by convolving the inverse transform of *Z*^*XY*^ with *Y* (see Methods and Materials; Torrence & Compo, 1998).

Supporting Information Table 1 and Figure 33 show estimates of admittance between environmental variables and amphibian and carnivoran richness series for the largest 25% of scales (wavelengths *≥* 3756 km). The mean of the transfer function between mean annual precipitation (i.e. *W*^*Y*^) and amphibian richness (*W*^*X*^) in the Americas, for example, is 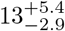 spx/m. These scales account for ≳ 60% of observed amphibian species richness. Stated uncertainties are distances to 1^st^ and 3^rd^ quartiles of admittance values, across all (*x, s*) space for *s ≥* 3756 km. Calculated admittance is most likely to be a reliable rule of thumb for converting environmental variable values into species richness when coherence is high. In Supporting Information Table 1, bold values indicate values where coherence is *>* 0.5 (see Figure 4). For example, consider the median large-scale coherence between amphibian richness and precipitation, which is 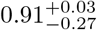 for the Americas. This result, coupled with the associated high positive admittance value, is suggestive of positive amphibian richness correlation with large-scale precipitation patterns. Compare that result to the relatively low median large-scale coherence between e.g. American carnivoran richness and annual temperature range, which is only 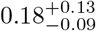, reflecting their likely independence. A comparison with results generated using global mean species richness and environmental variables indicates that these rules may be generally applicable (see Supporting Information Table 1 & Figure 33). However, there are key differences in correlations between richness in environment across different transects, suggesting that different environmental variables come to the forefront when governing species richness trends across longitudes (see Figure 4). We also caution against interpreting admittance values where there is lower coherence than could be generated by pairs of random signals with the same autocorrelation coefficients (i.e. 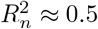 for the scales in question). In cases of high large-scale coherence, such simple rules-of-thumb appear to provide a means to predict species richness from external variables at large (≳ 1000 km) scales, but careful calibration and validation is necessary before such calculations are applied in practice or at global scales.

### 3.5 Drivers of Species Richness

In the Introduction of this paper we described five hypotheses to test. First, we hypothesized that species richness is highly coherent with environmental variables across all scales. Given the spectral analyses we present, which indicate that most species richness does not have statistically significant coherence with environmental variables at wavelengths ≲ 1000 km at most latitudes, this hypothesis can be rejected. Our second hypothesis—species richness is most coherent with external variability at small scales—is also thus rejected. Our third hypothesis—species richness is most coherent with changes in environment at large scales—was found to be reasonable. Generally, our results indicate that species richness is most coherent with environment at wavelengths ≳ 10^3^ km, where highest species richness power also resides. Our fourth hypothesis—that coherence of species richness with external variables depends on taxonomic group—was also found to be reasonable. For example, amphibian richness is found to be highly coherent with temperature range at scales *>* 5000 km, whereas carnivoran richness has very low coherence at these scales (cf. Figure 3d and h). Furthermore, amphibian richness seems to be more coherent with precipitation and temperature within the tropics, whereas carnivoran richness is not. Figure 4 summarizes the different distributions of coherence between large-scale species richness and environmental patterns for several terrestrial vertebrate taxa, across the Americas, Africa and Eurasia. Our fifth and final hypothesis—that species richness does not directly depend on environment, and that instead species richness depends upon biotic interactions—requires modification. Species richness was found to have both high and relatively low coherence with environmental variables, depending on location, scale and the environmental variable being considered. We have shown that wavelet transformation provides a means to identify coherence in the space-frequency domains. Our preliminary assessment of species-species interactions indicates that their coherence is also scale- and location-dependent (Supporting Information Figure 2). However, we note that calculated coherence and particularly cross power between species tends to be lower than that between species richness and environmental variables. We note that historical effects, i.e. speciation/extinction rates over geologic time, are not identified within this study, solely modern correlations between variables, although long-term speciation/extinction rates may themselves depend on environment (e.g. Skeels *et al*., 2022). Nonetheless, we tentatively suggest that these results indicate that environment is more important in determining species richness than species-species interactions.

### 3.6 Implications for Macroecological Biodiversity Patterns

A principal result of this study is that terrestrial species richness tends to be most coherent with topography, precipitation and temperature at long wavelengths (*>* 10^3^ km). These results indicate that large-scale variation in tectonic and climatic processes play a governing role in generating the latitudinal diversity gradient (e.g. Brodie & Mannion, 2023; Field *et al*., 2009). However, our results also indicate that the distribution of taxa, and their coherence and phase with environmental variables, is highly location- and scale-dependent. For example, whereas carnivorans and amphibians are in phase and coherent with mean annual precipitation and temperature at wavelengths *>* 10^4^ km, that is not true at smaller scales (i.e. shorter wavelengths). Significant deviations from the latitudinal diversity gradient indicate that external variables such as elevation, climatic patterns and tectonic history, play important roles in determining biodiversity at specific locations and scales (e.g. Archibald *et al*., 2010, 2013; Hagen *et al*., 2021; Jones *et al*., 2022; Mannion *et al*., 2014; Saupe, 2021; Song *et al*., 2020; Yasuhara *et al*., 2017).

Spectral analyses highlight the importance of the tropics for biodiversity, in particular for hummingbirds and amphibians, for which local changes in elevation and mean annual temperature (but not annual temperature range) are highly coherent with species richness. These results are consistent with the idea that increased resource availability in the tropics may generate higher primary productivity, supporting a greater number of individuals within a given area (i.e. higher carrying capacity), and therefore a greater number of different species (e.g. Fritz *et al*., 2016; Gillman *et al*., 2015; Hawkins *et al*., 2003; Kessler *et al*., 2014). Our results support the suggestion that elevated topography at the tropics is more likely to result in increased species diversity when compared to higher latitudes (Janzen, 1967; Ghalambor *et al*., 2006; Polato *et al*., 2018). However, this trend is not uniformly observed across taxa and for all continents. Species richness of carnivorans, for example, has no significant coherence with elevation or temperature range in the tropics, which suggests that this group is largely unaffected by the challenges posed by tropical mountain ranges. This might reflect the group ‘s relatively unusual biogeographical history and seemingly high dispersal ability, with carnivorans originating at high latitudes and dispersing into the tropics, with net diversification rates comparable in tropical and temperate regions (Rolland *et al*., 2015). Power spectral slopes for such taxa are steeper (more negative) at shorter wavelengths, whereas more environmentally-sensitive taxa, such as hummingbirds and amphibians, have shallower spectral slopes at longer wavelengths within tropical latitudes.

Cross wavelet power and coherence indicate that species richness is decoupled from short wavelength (≲ 10^3^ km) changes in elevation, temperature, annual temperature range and precipitation at nearly all locations, except for certain taxa within the tropics. Locally uplifted topography can be highly coherent with species richness. Trends across the Americas are reflected in global, latitudinally-averaged, transects and for other continents. In general, the species richness of taxa such as hummingbirds and amphibians is strongly and positively correlated with precipitation rate and temperature, except in Africa, where high temperatures may limit availability of water.

### 3.7 Conclusions

In summary, wavelet power spectral analysis provides insight into the coherence between species richness and environmental variables. Species richness is shown to vary as a function of location and scale. Comparisons with topography, temperature and precipitation show that species richness tends to be highly coherent with external forcing at large scales (wavelengths *>* 10^4^ km). Phase difference between signals reveals that species richness is in-phase with precipitation and temperature, and antiphase with elevation and annual temperature range, at these scales. That is to say, high temperatures and precipitation rates, which tend to occur within the tropics, generally relate to increased species richness, while high elevations and mean annual temperature ranges decrease richness. However, these relationships are dependent on scale and taxonomic group. At smaller scales, richness of bats, songbirds, hummingbirds and amphibians tends to be greatest in the tropics, where calculated coherence highlights the importance of topography and temperature range for determining species richness. Carnivorans, in contrast, show little coherence with environmental variables at these scales in the tropics. Instead, they are most coherent in the vicinity of terrestrial plateaux, for example the Colorado Plateau and Altiplano. These observations suggest that large scale (*>* 10^3^ km) variations in environmental variables determine almost all of the distribution of terrestrial vertebrates. Smaller scale (10^3^ km) variation can play an important role locally, particularly within the tropics. These results highlight the general importance of environmental change at the scale of tens degrees of latitude for determining biodiversity. They also indicate that changes at smaller scales are comparatively more important in the tropics for determining species richness. Crucially, these results could be used to predict the changes in biodiversity that could arise from different future Earth climate change scenarios.

## Acknowledgements

We thank P. Ball, C. Donaldson, S. Goes, F. Richards and A. Whittaker for helpful discussion. Two anonymous referees and M. Yasuhara provided helpful reviews. Figures were generated using Generic Mapping Tools v6.2.0 (Wessel *et al*., 2019).

