## Supporting Information for "Coherence of Terrestrial Vertebrate Species Richness with External Drivers Across Scales and Taxonomic Groups"

23rd February 2023

[1]Department of Earth Science & Engineering, Imperial College London, Royal School of Mines, Prince Consort Road, London SW7 2BP, UK

[2]Department of Earth Sciences, University College London, London WC1E 6BT, UK

[3]Department of Trait Diversity and Function, Royal Botanic Gardens, Kew, London TW9 3AE, UK

[4]now at Department of Biology, Phillipps-Universität Marburg, Karl-von-Frisch-Str. 8, 35043, Marburg, Germany

[a]**

[b]**

[c]**

[d]**

[e]**

### Investigation of Longitudinal Variability in Species Richness Patterns

The manuscript which this Supporting Information document accompanies shows results of wavelet power spectral analysis of species richness and environmental variables across the Americas (transect A—A′ in Figure 1 of main text), with coherence analysis for that transect and African and Eurasian transects (C—C′ and D—D′ in Figure 1 of the main text). The Materials and Methods section in the main text describes the methodologies used to generate the results presented in this document. The results are discussed in the main manuscript. Here, we carry out a number of tests to assess the reliability of results presented in the main text, with respect to different taxa, species-species interactions, noise distributions, transects, chosen mother wavelets, fitted spectral slopes, data processing conditions. The tests are listed as follows, in the order in which they are presented in this document:

1. Median and upper/lower quartile admittance and coherence values between species richness and environmental variables, for scales $\geq$ 3756 km, for all taxa studied (as in Figure 4 of the main text), across the Americas, Africa, Eurasia and global latitudinal mean transects are shown in Table 1.

2. Figure 1 shows transects, rectified wavelet power and filtered inverse transforms for species richness of taxa mapped by Jenkins *et al.* (2013) that are not included in the main manuscript. They either do not inhabit full latitudinal ranges or have similar results to those for taxa shown in the main text.

3. Cross wavelet power and phase difference, masked above a 90% coherence significance limit as in Figure 3 of the main text, are shown for species-species interactions between Carnivora and other mammals (Figure 2).

4. Rectified wavelet power spectra for amphibians across the Americas plus inserted white noise are shown in Figure 3.

5. Transects, rectified wavelet power and filtered inverse transforms for species richness and environmental variables, as in Figure 2 of the main text, for each of the transects B—B′ (Australia), C—C′ (Africa), D—D′ (Eurasia), and a global mean of each variable across latitudes are shown in Figures 4–7.

6. Rectified wavelet power of amphibian species richness across the Americas calculated using different mother wavelets is shown in Figure 8.

7. Figures 9–18 show best-fitting one- and two-spectral slope models, for both integer and non-integer slopes, for distance-averaged rectified power across the whole length of transects A—A′,

B—B′, C—C′, D—D′, and for global latitudinal averages.

8. We show distance-averaged rectified power for all taxa across all transects, and distance-averaged cross power between each taxon and each environmental variable, for (i) the entire transect, (ii) the tropical portion of the transect, and (iii) the portion of the transect outside the tropics (Figures 19–23).

9. Figures 24–32 show cross wavelet power and phase difference for carnivoran and amphibian species richness and environment for all transects in the main manuscript and for global latitudinal averages. Unmasked and masked (above a 90% coherence significance limit, as in Figure 3 of the main text) results are presented.

10. Figure 33 shows filtered inverse transforms for species richness transects at scales (Fourier periods) $\geq$ 3756 km. Mean coherence and admittance values are shown for both carnivorans and amphibians along transect A—A′ and global mean transects.

11. We present rectified wavelet power for mirrored and un-mirrored amphibian species richness across the American, Australian, African and Eurasian transects. Mirroring of the transects prior to transformation into the spectral domain is performed to reduce edge effects. Figures 34 and 35 show histograms of coherence between species richness and environmental variables for all taxa, across the Americas, Africa and Eurasia for the mirrored and un-mirrored transects.

12. Wavelet power spectra of amphibian species richness along three different transects spanning the Americas, in addition to the one analyzed in the main text and other figures in this document (A—A′), are shown in Figures 36–40. Calculated coherence between amphibian species richness and environmental variables along each transect, as a function of latitude and scale, is shown in Figure 41.

**Table 1: Rules of thumb to convert values of environmental variables into species richness**. $Z$ = admittance (transfer function) between environmental variables and species richness at large scales (> 3756 km); subscripts $E$, $Pn$, $Tm$ & $\Delta T$ indicate admittance/coherence between richness of given taxonomic group and elevation, annual precipitation, mean annual temperature, and annual temperature range respectively; units are spx/km ($E$), spx/m ($Pn$) and spx/°C ($Tm$ and $\Delta T$), where spx = species per pixel (see Figure 1 of main text and Figure 1 of this document). Bold text = admittance values for species-variable relationships with median coherence > 0.5 (90% coherence confidence threshold for scales ≥ 3756 km). $R_n^2$ = coherence between environmental variable and species; bold text = $R_n^2 > 0.5$. Values stated for Americas (A—A′), Africa (C—C′), Asia (D—D′), and global averages. Australian transect length < 3756 km and so values are undefined for that scale limit. $M$ = Mammalia, $Ca$ = Carnivora, $Ch$ = Chiroptera, $M^*$ = Mammlia excluding Carnivora, $Ct$ = Cetartiodactyla, $Eu$ = Eulipotyphla, $Pr$ = Primates, $Mr$ = Marsupialia, $Ro$ = Rodentia, $Pa$ = Passeriformes, $Tr$ = Trochilidae, $Ps$ = Psittaciformes, $Am$ = Amphibia, $An$ = Anura.

| Group | $Z_E$ | $R^2_{nE}$ | $Z_{Pn}$ | $R^2_{nPn}$ | $Z_{Tm}$ | $R^2_{nTm}$ | $Z_{\Delta T}$ | $R^2_{n\Delta T}$ |
|---|---|---|---|---|---|---|---|---|
| (Americas) | | | | | | | | |
| $M$ | $\mathbf{-22^{+11}_{-7.4}}$ | $\mathbf{0.71^{+0.09}_{-0.37}}$ | $\mathbf{36^{+14}_{-9.2}}$ | $\mathbf{0.81^{+0.07}_{-0.09}}$ | $3.3^{+2.7}_{-2.4}$ | $0.42^{+0.20}_{-0.23}$ | $-2.2^{+2.0}_{-1.3}$ | $0.25^{+0.31}_{-0.14}$ |
| $Ca$ | $\mathbf{-1.7^{+0.60}_{-0.90}}$ | $\mathbf{0.70^{+0.09}_{-0.20}}$ | $\mathbf{2.3^{+1.9}_{-0.81}}$ | $\mathbf{0.55^{+0.14}_{-0.22}}$ | $0.19^{+0.20}_{-0.30}$ | $0.25^{+0.14}_{-0.10}$ | $-0.10^{+0.23}_{-0.14}$ | $0.18^{+0.13}_{-0.09}$ |
| $Ch$ | $\mathbf{-15^{+7.6}_{-2.9}}$ | $\mathbf{0.77^{+0.09}_{-0.39}}$ | $\mathbf{25^{+7.0}_{-7.6}}$ | $\mathbf{0.88^{+0.04}_{-0.12}}$ | $2.1^{+1.7}_{-1.6}$ | $0.37^{+0.23}_{-0.23}$ | $-1.4^{+1.3}_{-0.91}$ | $0.23^{+0.36}_{-0.13}$ |
| $M^*$ | $\mathbf{-19^{+11}_{-7.3}}$ | $\mathbf{0.68^{+0.09}_{-0.37}}$ | $\mathbf{34^{+13}_{-8.4}}$ | $\mathbf{0.80^{+0.09}_{-0.10}}$ | $3.0^{+2.7}_{-2.0}$ | $0.41^{+0.21}_{-0.23}$ | $-2.0^{+1.8}_{-1.2}$ | $0.25^{+0.32}_{-0.14}$ |
| $Ct$ | $-0.20^{+0.24}_{-0.39}$ | $0.27^{+0.13}_{-0.17}$ | $0.68^{+0.64}_{-0.55}$ | $0.50^{+0.10}_{-0.12}$ | $0.01^{+0.10}_{-0.04}$ | $0.08^{+0.21}_{-0.05}$ | $0.0^{+0.04}_{-0.05}$ | $0.09^{+0.13}_{-0.05}$ |
| $Eu$ | $-0.02^{+0.37}_{-0.27}$ | $0.06^{+0.12}_{-0.13}$ | $-0.01^{+0.56}_{-0.45}$ | $0.08^{+0.14}_{-0.05}$ | $-0.03^{+0.05}_{-0.06}$ | $0.26^{+0.14}_{-0.12}$ | $0.02^{+0.05}_{-0.04}$ | $0.30^{+0.18}_{-0.16}$ |
| $Pr$ | $\mathbf{-1.0^{+1.0}_{-0.8}}$ | $\mathbf{0.53^{+0.11}_{-0.18}}$ | $\mathbf{1.6^{+0.49}_{-0.63}}$ | $\mathbf{0.76^{+0.12}_{-0.19}}$ | $0.08^{+0.26}_{-0.11}$ | $0.19^{+0.21}_{-0.10}$ | $-0.04^{+0.09}_{-0.12}$ | $0.14^{+0.21}_{-0.08}$ |
| $Mr$ | $-1.0^{+1.3}_{-1.0}$ | $0.49^{+0.07}_{-0.19}$ | $\mathbf{2.5^{+1.0}_{-1.8}}$ | $\mathbf{0.68^{+0.20}_{-0.28}}$ | $0.08^{+0.36}_{-0.19}$ | $0.17^{+0.18}_{-0.08}$ | $-0.10^{+0.16}_{-0.25}$ | $0.19^{+0.22}_{-0.12}$ |
| $Ro$ | $-1.3^{+2.6}_{-2.6}$ | $0.08^{+0.11}_{-0.06}$ | $2.2^{+4.1}_{-3.0}$ | $0.17^{+0.30}_{-0.12}$ | $\mathbf{0.45^{+0.51}_{-0.22}}$ | $\mathbf{0.51^{+0.24}_{-0.14}}$ | $-0.16^{+0.27}_{-0.28}$ | $0.28^{+0.24}_{-0.15}$ |
| $Pa$ | $\mathbf{-29^{+17}_{-23}}$ | $\mathbf{0.63^{+0.08}_{-0.37}}$ | $\mathbf{57^{+19}_{-12}}$ | $\mathbf{0.78^{+0.14}_{-0.11}}$ | $4.5^{+6.5}_{-2.9}$ | $0.43^{+0.16}_{-0.25}$ | $-3.0^{+2.6}_{-3.0}$ | $0.30^{+0.24}_{-0.15}$ |
| $Tr$ | $\mathbf{-4.2^{+3.0}_{-1.3}}$ | $\mathbf{0.74^{+0.07}_{-0.41}}$ | $\mathbf{7.4^{+1.1}_{-1.5}}$ | $\mathbf{0.90^{+0.08}_{-0.21}}$ | $0.42^{+0.71}_{-0.40}$ | $0.26^{+0.21}_{-0.16}$ | $-0.34^{+0.35}_{-0.37}$ | $0.21^{+0.29}_{-0.12}$ |
| $Ps$ | $\mathbf{-1.8^{+1.7}_{-1.0}}$ | $\mathbf{0.68^{+0.12}_{-0.30}}$ | $\mathbf{3.0^{+0.61}_{-0.90}}$ | $\mathbf{0.90^{+0.04}_{-0.14}}$ | $0.17^{+0.33}_{-0.19}$ | $0.22^{+0.22}_{-0.13}$ | $-0.11^{+0.14}_{-0.19}$ | $0.19^{+0.22}_{-0.10}$ |
| $Am$ | $\mathbf{-7.5^{+3.2}_{-2.9}}$ | $\mathbf{0.56^{+0.26}_{-0.28}}$ | $\mathbf{13^{+5.4}_{-2.9}}$ | $\mathbf{0.91^{+0.03}_{-0.27}}$ | $0.94^{+0.87}_{-0.96}$ | $0.27^{+0.21}_{-0.15}$ | $-0.52^{+0.94}_{-0.46}$ | $0.14^{+0.30}_{-0.08}$ |
| $An$ | $\mathbf{-7.0^{+3.3}_{-2.8}}$ | $\mathbf{0.58^{+0.24}_{-0.29}}$ | $\mathbf{12^{+4.2}_{-2.3}}$ | $\mathbf{0.93^{+0.03}_{-0.28}}$ | $0.80^{+0.89}_{-0.84}$ | $0.26^{+0.20}_{-0.15}$ | $-0.47^{+0.90}_{-0.44}$ | $0.15^{+0.29}_{-0.09}$ |
| (Africa) | $Z_E$ | $R^2_{nE}$ | $Z_{Pn}$ | $R^2_{nPn}$ | $Z_{Tm}$ | $R^2_{nTm}$ | $Z_{\Delta T}$ | $R^2_{n\Delta T}$ |
| $M$ | $-29^{+21}_{-37}$ | $0.28^{+0.13}_{-0.15}$ | $\mathbf{43^{+9.0}_{-13}}$ | $\mathbf{0.84^{+0.06}_{-0.08}}$ | $-6.0^{+10}_{-6.6}$ | $0.37^{+0.18}_{-0.15}$ | $\mathbf{-2.6^{+0.73}_{-0.97}}$ | $\mathbf{0.76^{+0.05}_{-0.06}}$ |

| | | | | | | | | |
|---|---|---|---|---|---|---|---|---|
| $Ca$ | $2.1^{+3.2}_{-3.5}$ | $0.23^{+0.18}_{-0.17}$ | $0.54^{+2.8}_{-3.2}$ | $0.15^{+0.17}_{-0.07}$ | $-0.31^{+0.60}_{-0.45}$ | $0.26^{+0.15}_{-0.09}$ | $-0.05^{+0.23}_{-0.16}$ | $0.11^{+0.12}_{-0.06}$ |
| $Ch$ | $\mathbf{-16^{+8.8}_{-16}}$ | $\mathbf{0.51^{+0.11}_{-0.28}}$ | $\mathbf{20^{+1.5}_{-1.4}}$ | $\mathbf{0.98^{+0.01}_{-0.01}}$ | $-1.4^{+4.2}_{-4.8}$ | $0.27^{+0.24}_{-0.14}$ | $\mathbf{-1.2^{+0.27}_{-0.44}}$ | $\mathbf{0.84^{+0.04}_{-0.05}}$ |
| $M$* | $-31^{+19}_{-34}$ | $0.36^{+0.11}_{-0.24}$ | $\mathbf{43^{+5.8}_{-10}}$ | $\mathbf{0.89^{+0.04}_{-0.04}}$ | $-4.9^{+9.3}_{-7.7}$ | $0.33^{+0.19}_{-0.14}$ | $\mathbf{-2.5^{+0.61}_{-0.86}}$ | $\mathbf{0.81^{+0.05}_{-0.05}}$ |
| $Ct$ | $-1.0^{+3.2}_{-5.3}$ | $0.12^{+0.13}_{-0.06}$ | $3.8^{+2.1}_{-3.1}$ | $0.45^{+0.20}_{-0.31}$ | $-0.57^{+0.81}_{-0.58}$ | $0.43^{+0.12}_{-0.14}$ | $-0.24^{+0.20}_{-0.15}$ | $0.45^{+0.15}_{-0.28}$ |
| $Eu$ | $\mathbf{-6.8^{+1.5}_{-2.1}}$ | $\mathbf{0.67^{+0.11}_{-0.17}}$ | $\mathbf{5.6^{+3.1}_{-1.5}}$ | $\mathbf{0.74^{+0.02}_{-0.02}}$ | $0.31^{+1.1}_{-1.7}$ | $0.19^{+0.21}_{-0.15}$ | $\mathbf{-0.43^{+0.07}_{-0.10}}$ | $\mathbf{0.89^{+0.04}_{-0.11}}$ |
| $Pr$ | $\mathbf{-5.1^{+2.2}_{-3.6}}$ | $\mathbf{0.59^{+0.07}_{-0.40}}$ | $\mathbf{5.6^{+1.7}_{-0.57}}$ | $\mathbf{0.95^{+0.01}_{-0.03}}$ | $-0.31^{+1.3}_{-1.3}$ | $0.24^{+0.18}_{-0.15}$ | $\mathbf{-0.36^{+0.04}_{-0.08}}$ | $\mathbf{0.89^{+0.05}_{-0.05}}$ |
| $Mr$ | - | - | - | - | - | - | - | - |
| $Ro$ | $-0.88^{+4.9}_{-9.1}$ | $0.14^{+0.14}_{-0.09}$ | $\mathbf{5.7^{+2.2}_{-4.7}}$ | $\mathbf{0.51^{+0.19}_{-0.26}}$ | $\mathbf{-1.4^{+1.0}_{-0.6}}$ | $\mathbf{0.53^{+0.14}_{-0.18}}$ | $-0.03^{+0.01}_{-0.02}$ | $0.42^{+0.16}_{-0.20}$ |
| $Pa$ | $-50^{+28}_{-48}$ | $0.43^{+0.11}_{-0.29}$ | $\mathbf{65^{+7.6}_{-11}}$ | $\mathbf{0.94^{+0.03}_{-0.05}}$ | $-7.6^{+15}_{-11}$ | $0.31^{+0.23}_{-0.16}$ | $\mathbf{-4.0^{+0.96}_{-1.1}}$ | $\mathbf{0.85^{+0.04}_{-0.04}}$ |
| $Tr$ | - | - | - | - | - | - | - | - |
| $Ps$ | $-0.30^{+0.33}_{-0.29}$ | $0.14^{+0.06}_{-0.08}$ | $0.44^{+0.37}_{-0.13}$ | $0.50^{+0.10}_{-0.12}$ | $-0.04^{+0.11}_{-0.07}$ | $0.20^{+0.16}_{-0.12}$ | $-0.03^{+0.01}_{-0.02}$ | $0.38^{+0.06}_{-0.09}$ |
| $Am$ | $-17^{+9.3}_{-15}$ | $0.48^{+0.11}_{-0.35}$ | $\mathbf{20^{+4.5}_{-3.2}}$ | $\mathbf{0.93^{+0.02}_{-0.02}}$ | $-1.8^{+4.6}_{-4.0}$ | $0.29^{+0.24}_{-0.16}$ | $\mathbf{-1.3^{+0.18}_{-0.32}}$ | $\mathbf{0.89^{+0.03}_{-0.06}}$ |
| $An$ | $-16^{+9.0}_{-15}$ | $0.45^{+0.12}_{-0.33}$ | $\mathbf{20^{+4.8}_{-3.7}}$ | $\mathbf{0.91^{+0.03}_{-0.04}}$ | $-1.8^{+4.4}_{-3.9}$ | $0.30^{+0.23}_{-0.15}$ | $\mathbf{-1.2^{+0.19}_{-0.33}}$ | $\mathbf{0.88^{+0.04}_{-0.07}}$ |

| (Asia) | $Z_E$ | $R^2_{nE}$ | $Z_{Pn}$ | $R^2_{nPn}$ | $Z_{Tm}$ | $R^2_{nTm}$ | $Z_{\Delta T}$ | $R^2_{n\Delta T}$ |
|---|---|---|---|---|---|---|---|---|
| $M$ | $-8.0^{+9.0}_{-15}$ | $0.30^{+0.13}_{-0.12}$ | $\mathbf{43^{+21}_{-23}}$ | $\mathbf{0.51^{+0.10}_{-0.15}}$ | $\mathbf{0.95^{+0.25}_{-0.11}}$ | $\mathbf{0.94^{+0.02}_{-0.05}}$ | $\mathbf{-1.5^{+0.43}_{-0.36}}$ | $\mathbf{0.83^{+0.04}_{-0.16}}$ |
| $Ca$ | $-1.2^{+1.6}_{-2.5}$ | $0.26^{+0.15}_{-0.10}$ | $\mathbf{8.1^{+3.9}_{-4.0}}$ | $\mathbf{0.57^{+0.08}_{-0.12}}$ | $\mathbf{0.17^{+0.03}_{-0.01}}$ | $\mathbf{0.96^{+0.01}_{-0.02}}$ | $\mathbf{-0.28^{+0.07}_{-0.06}}$ | $\mathbf{0.86^{+0.03}_{-0.09}}$ |
| $Ch$ | $-3.8^{+4.1}_{-5.1}$ | $0.16^{+0.12}_{-0.08}$ | $\mathbf{26^{+7.6}_{-6.8}}$ | $\mathbf{0.74^{+0.06}_{-0.08}}$ | $\mathbf{0.44^{+0.07}_{-0.07}}$ | $\mathbf{0.95^{+0.01}_{-0.03}}$ | $\mathbf{-0.77^{+0.14}_{-0.12}}$ | $\mathbf{0.90^{+0.05}_{-0.06}}$ |
| $M$* | $-6.8^{+7.2}_{-12}$ | $0.31^{+0.13}_{-0.12}$ | $35^{+17}_{-20}$ | $0.48^{+0.10}_{-0.15}$ | $\mathbf{0.77^{+0.22}_{-0.09}}$ | $\mathbf{0.93^{+0.02}_{-0.06}}$ | $\mathbf{-1.2^{+0.36}_{-0.31}}$ | $\mathbf{0.82^{+0.04}_{-0.17}}$ |
| $Ct$ | $-2.3^{+1.7}_{-2.0}$ | $0.30^{+0.10}_{-0.13}$ | $10^{+2.9}_{-3.9}$ | $0.47^{+0.19}_{-0.18}$ | $\mathbf{0.18^{+0.03}_{-0.02}}$ | $\mathbf{0.89^{+0.05}_{-0.11}}$ | $\mathbf{-0.29^{+0.08}_{-0.07}}$ | $\mathbf{0.76^{+0.10}_{-0.23}}$ |
| $Eu$ | $-0.67^{+1.3}_{-1.4}$ | $0.35^{+0.06}_{-0.06}$ | $-2.8^{+2.0}_{-4.0}$ | $0.24^{+0.09}_{-0.12}$ | $-0.01^{+0.07}_{-0.05}$ | $0.17^{+0.13}_{-0.09}$ | $0.05^{+0.15}_{-0.07}$ | $0.17^{+0.14}_{-0.11}$ |
| $Pr$ | $-0.97^{+0.54}_{-0.56}$ | $0.32^{+0.13}_{-0.23}$ | $\mathbf{4.6^{+0.89}_{-1.2}}$ | $\mathbf{0.72^{+0.07}_{-0.13}}$ | $\mathbf{0.07^{+0.02}_{-0.02}}$ | $\mathbf{0.76^{+0.10}_{-0.10}}$ | $\mathbf{-0.10^{+0.05}_{-0.03}}$ | $\mathbf{0.64^{+0.16}_{-0.20}}$ |
| $Mr$ | - | - | - | - | - | - | - | - |
| $Ro$ | $-0.39^{+2.1}_{-4.5}$ | $0.27^{+0.10}_{-0.12}$ | $-2.7^{+6.1}_{-8.8}$ | $0.31^{+0.10}_{-0.16}$ | $\mathbf{0.10^{+0.19}_{-0.11}}$ | $\mathbf{0.57^{+0.08}_{-0.15}}$ | $\mathbf{-0.10^{+0.23}_{-0.20}}$ | $\mathbf{0.52^{+0.12}_{-0.22}}$ |
| $Pa$ | $-7.7^{+19}_{-26}$ | $0.20^{+0.18}_{-0.12}$ | $\mathbf{80^{+44}_{-45}}$ | $\mathbf{0.59^{+0.08}_{-0.13}}$ | $\mathbf{1.7^{+0.20}_{-0.18}}$ | $\mathbf{0.89^{+0.04}_{-0.02}}$ | $\mathbf{-2.8^{+0.68}_{-0.65}}$ | $\mathbf{0.81^{+0.08}_{-0.06}}$ |

| | | | | | | | | |
|---|---|---|---|---|---|---|---|---|
| *Tr* | - | - | - | - | - | - | - | - |
| *Ps* | $-0.83^{+0.70}_{-0.83}$ | $0.23^{+0.13}_{-0.12}$ | $\mathbf{5.4^{+0.74}_{-0.83}}$ | $\mathbf{0.85^{+0.04}_{-0.03}}$ | $\mathbf{0.08^{+0.02}_{-0.02}}$ | $\mathbf{0.86^{+0.03}_{-0.05}}$ | $\mathbf{-0.13^{+0.04}_{-0.04}}$ | $\mathbf{0.79^{+0.07}_{-0.09}}$ |
| *Am* | $-4.2^{+3.6}_{-5.3}$ | $0.28^{+0.10}_{-0.10}$ | $\mathbf{26^{+4.3}_{-7.0}}$ | $\mathbf{0.78^{+0.06}_{-0.10}}$ | $\mathbf{0.44^{+0.03}_{-0.06}}$ | $\mathbf{0.95^{+0.02}_{-0.03}}$ | $\mathbf{-0.76^{+0.16}_{-0.07}}$ | $\mathbf{0.95^{+0.02}_{-0.05}}$ |
| *An* | $-4.0^{+3.7}_{-5.4}$ | $0.26^{+0.10}_{-0.10}$ | $\mathbf{26^{+4.6}_{-7.3}}$ | $\mathbf{0.79^{+0.06}_{-0.10}}$ | $\mathbf{0.43^{+0.03}_{-0.05}}$ | $\mathbf{0.95^{+0.02}_{-0.03}}$ | $\mathbf{-0.76^{+0.13}_{-0.06}}$ | $\mathbf{0.96^{+0.01}_{-0.04}}$ |

| (Global) | $Z_E$ | $R^2_{nE}$ | $Z_{Pn}$ | $R^2_{nPn}$ | $Z_{Tm}$ | $R^2_{nTm}$ | $Z_{\Delta T}$ | $R^2_{n\Delta T}$ |
|---|---|---|---|---|---|---|---|---|
| *M* | $-42^{+42}_{-66}$ | $0.20^{+0.12}_{-0.08}$ | $38^{+26}_{-16}$ | $0.48^{+0.19}_{-0.16}$ | $0.40^{+2.2}_{-3.4}$ | $0.21^{+0.28}_{-0.13}$ | $-1.6^{+1.4}_{-1.3}$ | $0.26^{+0.15}_{-0.13}$ |
| *Ca* | $-3.1^{+4.0}_{-5.2}$ | $0.25^{+0.15}_{-0.13}$ | $2.3^{+2.2}_{-1.5}$ | $0.34^{+0.15}_{-0.18}$ | $0.00^{+0.17}_{-0.31}$ | $0.21^{+0.14}_{-0.08}$ | $-0.09^{+0.14}_{-0.13}$ | $0.15^{+0.10}_{-0.09}$ |
| *Ch* | $-21^{+33}_{-43}$ | $0.19^{+0.15}_{-0.09}$ | $\mathbf{21^{+18}_{-7.6}}$ | $\mathbf{0.56^{+0.17}_{-0.08}}$ | $0.24^{+1.8}_{-2.3}$ | $0.35^{+0.27}_{-0.18}$ | $-0.95^{+0.71}_{-0.82}$ | $0.35^{+0.15}_{-0.20}$ |
| *M** | $-45^{+44}_{-56}$ | $0.22^{+0.12}_{-0.09}$ | $\mathbf{37^{+22}_{-14}}$ | $\mathbf{0.53^{+0.18}_{-0.14}}$ | $0.33^{+2.0}_{-3.3}$ | $0.23^{+0.30}_{-0.15}$ | $-1.5^{+1.2}_{-1.3}$ | $0.30^{+0.14}_{-0.13}$ |
| *Ct* | $-3.1^{+3.0}_{-7.3}$ | $0.18^{+0.10}_{-0.07}$ | $1.6^{+1.4}_{-1.0}$ | $0.36^{+0.15}_{-0.14}$ | $0.01^{+0.17}_{-0.19}$ | $0.27^{+0.24}_{-0.16}$ | $-0.11^{+0.10}_{-0.08}$ | $0.26^{+0.14}_{-0.13}$ |
| *Eu* | $-2.4^{+3.8}_{-3.0}$ | $0.31^{+0.14}_{-0.13}$ | $0.91^{+2.2}_{-3.2}$ | $0.24^{+0.21}_{-0.13}$ | $-0.07^{+0.11}_{-0.12}$ | $0.12^{+0.16}_{-0.08}$ | $0.07^{+0.08}_{-0.18}$ | $0.12^{+0.20}_{-0.07}$ |
| *Pr* | $-5.0^{+5.4}_{-7.1}$ | $0.20^{+0.20}_{-0.14}$ | $\mathbf{4.2^{+0.21}_{-0.48}}$ | $\mathbf{0.99^{+0.01}_{-0.01}}$ | $-0.03^{+0.72}_{-0.56}$ | $0.20^{+0.14}_{-0.10}$ | $\mathbf{-0.31^{+0.14}_{-0.10}}$ | $\mathbf{0.80^{+0.07}_{-0.20}}$ |
| *Mr* | $-0.50^{+2.6}_{-10}$ | $0.10^{+0.13}_{-0.07}$ | $0.29^{+2.6}_{-1.4}$ | $0.22^{+0.11}_{-0.10}$ | $0.30^{+0.54}_{-0.42}$ | $0.09^{+0.19}_{-0.07}$ | $0.00^{+0.10}_{-0.20}$ | $0.33^{+0.07}_{-0.13}$ |
| *Ro* | $-3.5^{+7.1}_{-6.6}$ | $0.23^{+0.12}_{-0.12}$ | $4.5^{+3.8}_{-3.1}$ | $0.31^{+0.26}_{-0.11}$ | $0.04^{+0.20}_{-0.53}$ | $0.10^{+0.17}_{-0.06}$ | $-0.11^{+0.22}_{-0.32}$ | $0.16^{+0.10}_{-0.11}$ |
| *Pa* | $-2.5^{+30}_{-39}$ | $0.10^{+0.12}_{-0.07}$ | $55^{+82}_{-44}$ | $0.39^{+0.25}_{-0.15}$ | $-0.02^{+2.8}_{-3.1}$ | $0.20^{+0.24}_{-0.13}$ | $-2.0^{+2.5}_{-2.4}$ | $0.19^{+0.14}_{-0.10}$ |
| *Tr* | $-8.4^{+5.4}_{-14}$ | $0.37^{+0.11}_{-0.21}$ | $\mathbf{6.1^{+9.0}_{-3.0}}$ | $\mathbf{0.52^{+0.07}_{-0.07}}$ | $0.33^{+0.96}_{-0.84}$ | $0.41^{+0.22}_{-0.20}$ | $-0.31^{+0.24}_{-0.26}$ | $0.45^{+0.10}_{-0.16}$ |
| *Ps* | $1.2^{+3.7}_{-3.8}$ | $0.29^{+0.14}_{-0.15}$ | $0.20^{+1.5}_{-5.5}$ | $0.09^{+0.10}_{-0.05}$ | $-0.25^{+0.29}_{-0.30}$ | $0.18^{+0.13}_{-0.08}$ | $-0.08^{+0.12}_{-0.12}$ | $0.20^{+0.08}_{-0.08}$ |
| *Am* | $-7.3^{+24}_{-50}$ | $0.14^{+0.10}_{-0.06}$ | $13^{+11}_{-6.2}$ | $0.44^{+0.14}_{-0.14}$ | $0.23^{+1.4}_{-1.8}$ | $0.35^{+0.22}_{-0.18}$ | $-0.74^{+0.65}_{-0.52}$ | $0.30^{+0.19}_{-0.19}$ |
| *An* | $-8.2^{+23}_{-52}$ | $0.14^{+0.11}_{-0.07}$ | $12^{+11}_{-5.8}$ | $0.46^{+0.12}_{-0.14}$ | $0.17^{+1.4}_{-1.6}$ | $0.36^{+0.20}_{-0.18}$ | $-0.72^{+0.65}_{-0.51}$ | $0.32^{+0.19}_{-0.20}$ |

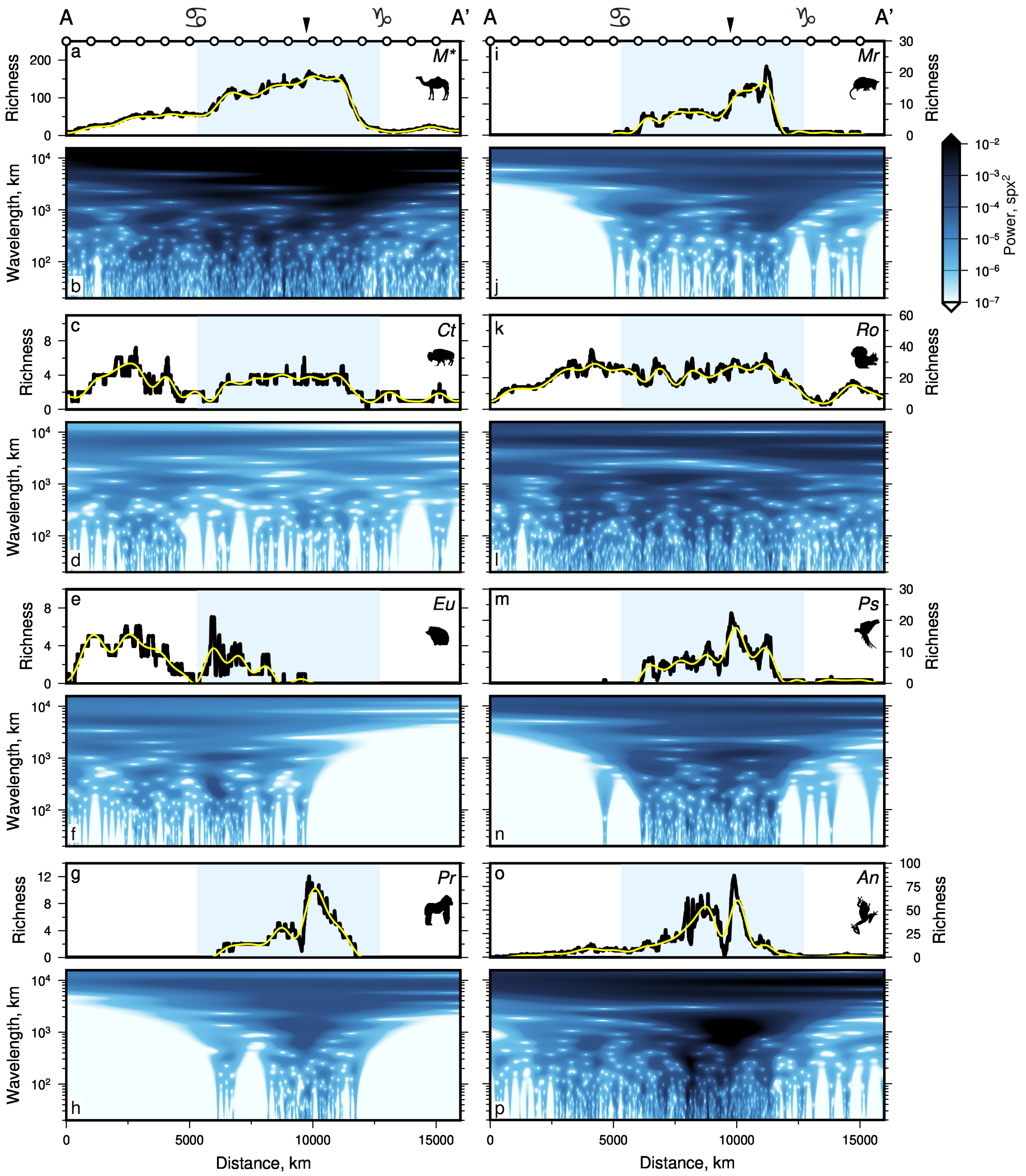


**Figure 1: Spectral analyses of other taxonomic groups**. As Figure 2 of main text but for other taxonomic groups not in main text. Mean differences between signals and inverse transforms filtered to remove wavelengths < 1000 km = 3.6 spx (*M**, Mammalia without Carnivora), 0.3 spx (*Ct*, Cetartiodactyla), 0.6 spx (*Eu*, Eulipotyphla), 0.5 spx (*Pr*, Primates), 0.6 spx (*Mr*, Marsupialia), 1.7 spx (*Ro*, Rodentia), 1.1 spx (*Ps*, Psittaciformes), and 2.7 spx (*An*, Anura).

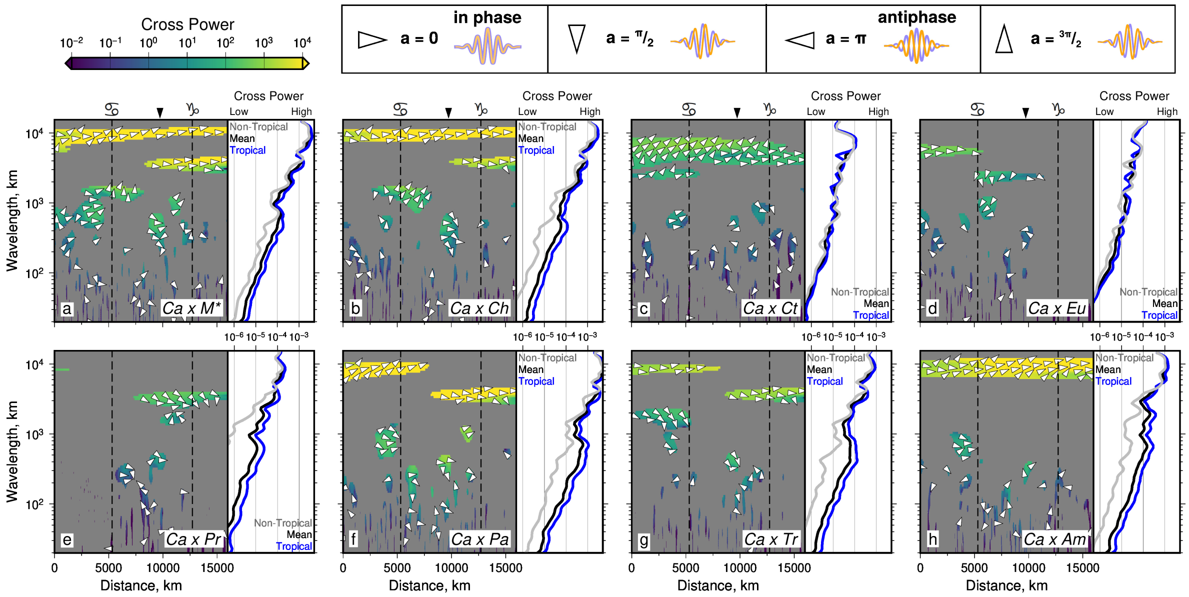


**Figure 2: Species-species interactions presented in the scale-distance domain.** As Figure 3 of main text but for species-species interactions between carnivorans ($Ca$) and all other mammals ($M^*$), Chiroptera ($Ch$), Cetartiodactyla ($Ct$), Eulipotyphla ($Eu$), Primates ($Pr$), Passeriformes ($Pa$), Trochilidae ($Tr$), and Amphibia ($Am$).

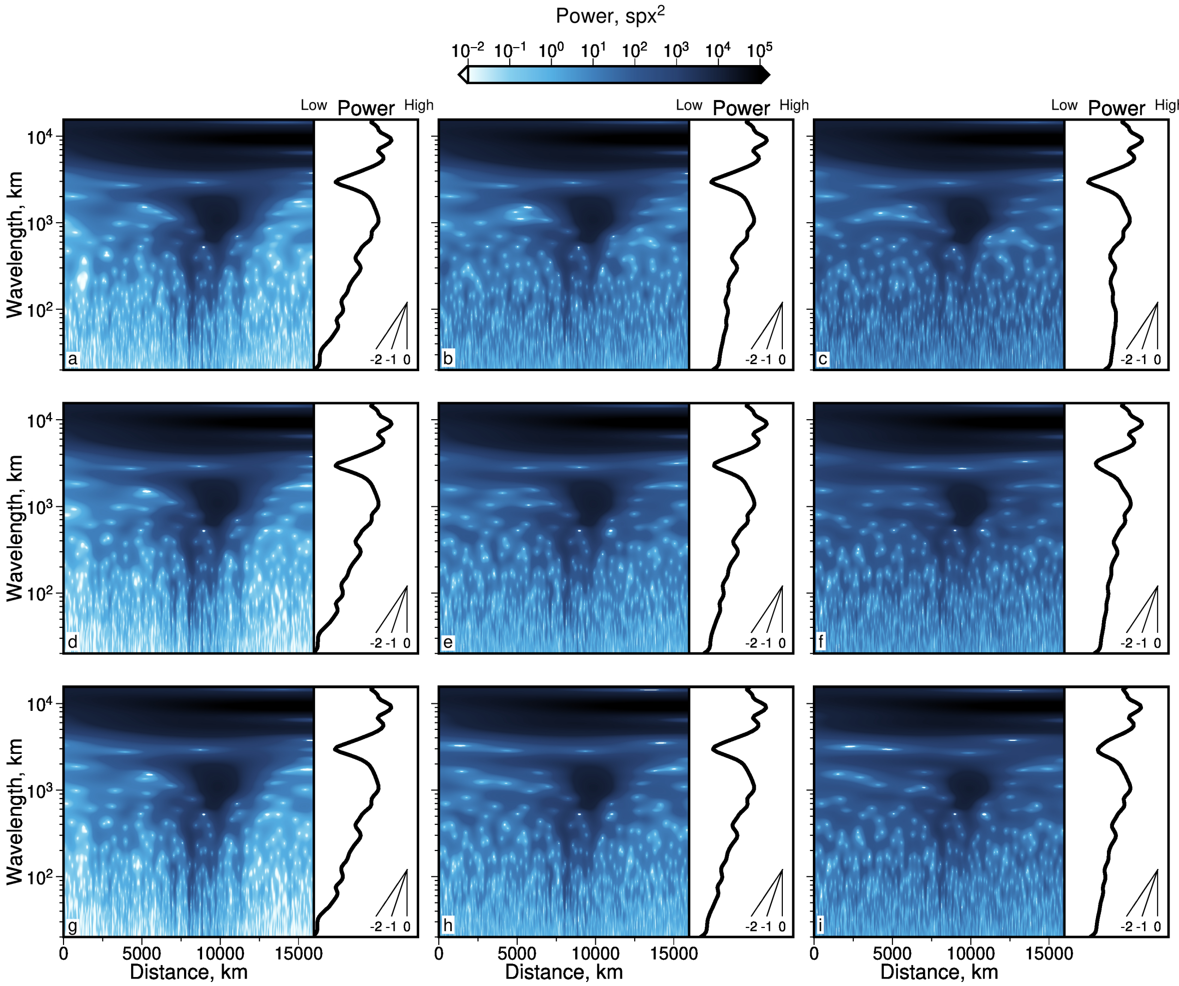


**Figure 3: Testing impact of noise on signals used for continuous wavelet transforms.** (a) Species richness power of Amphibia along transect A—A′, where white noise was first added to the input signal. Minimum wavelength of noise $\lambda_{\min}$ was 20 km in each case. Added noise has maximum wavelength $\lambda_{\max}$ of 100 km, and maximum amplitudes $\delta_{\max}$ of $0.1\sigma$, where $\sigma$ = standard deviation of noise-free signal (= 22.3 spx). Side panel = distance-averaged power as a function of wavelength, across entire transect; see inset slope guide for spectral slopes of $-2$ (red noise), $-1$ (pink noise) and 0 (white noise). (b) As (a) but $\delta_{\max} = 0.5\sigma$. (c) As (a) but $\delta_{\max} = 1\sigma$. (d)–(f) As (a)–(c) but $\lambda_{\max}$ = 1,000 km. (g)–(i) As (a)–(c) but $\lambda_{\max}$ = 10,000 km.

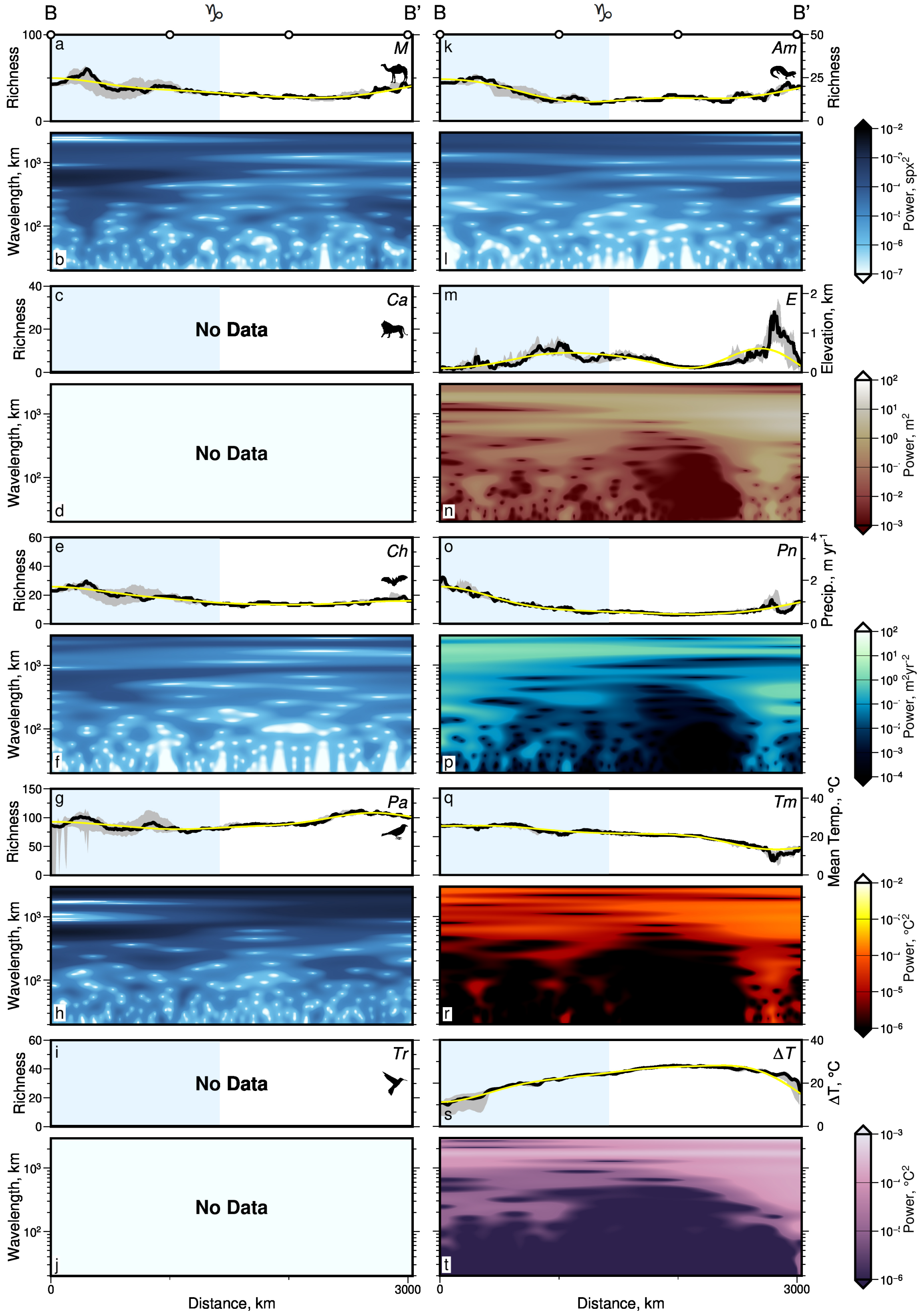
B
B'
Richness
Wavelength, km
No Data
Distance, km
Elevation, km
Precip., m yr$^{-1}$
Mean Temp., °C
ΔT, °C
Power, spx$^2$
Power, m$^2$
Power, m$^2$yr$^{-2}$
Power, °C$^2$
M
Am
Ca
E
Ch
Pn
Pa
Tm
Tr
ΔT
0
3000

**Figure 4**

**Figure 4 *(preceding page)*: Spectral analyses of Australian species richness and environmental variables.** (a) Black line = species richness of Mammalia ($M$) along transect B—B′ (Australia); gray bands = 100 km wide swaths centred on B—B′; blue bands = tropical latitudes; white circles are shown every 1,000 km, see transect B—B′ on Figure 1 of main text; symbol above top axis = Tropic of Capricorn. Yellow line = inverse wavelet transform of signal, filtering to pass only wavelengths > 1000 km; mean difference to input signal = 2.2 spx. (b) Continuous wavelet transform of Mammalia spatial series (black line in panel a). Colors = spectral power as a function of location and scale (wavelength); spx = species per pixel. (c)–(t) As (a)–(b) but for Carnivora ($Ca$), Chiroptera ($Ch$), Passeriformes ($Pa$), Trochilidae ($Tr$), Amphibia ($Am$), elevation ($E$), mean annual precipitation rate ($Pn$), temperature ($Tm$) and temperature range ($\Delta T$) along transect B—B′ Amante & Eakins (2009); Jenkins *et al.* (2013); Karger *et al.* (2017). Panels (c)–(d) and (i)–(j) empty due to absence of Carnivora and Trochilidae in the region. Mean differences between signals and inverse transforms filtered to remove wavelengths < 1000 km = 1.1 spx ($Ch$), 3.2 spx ($Pa$), 1.1 spx ($Am$), 0.11 km ($E$), 0.06 m/yr ($Pn$), 0.8 °C ($Tm$), and 0.7 °C ($\Delta T$).

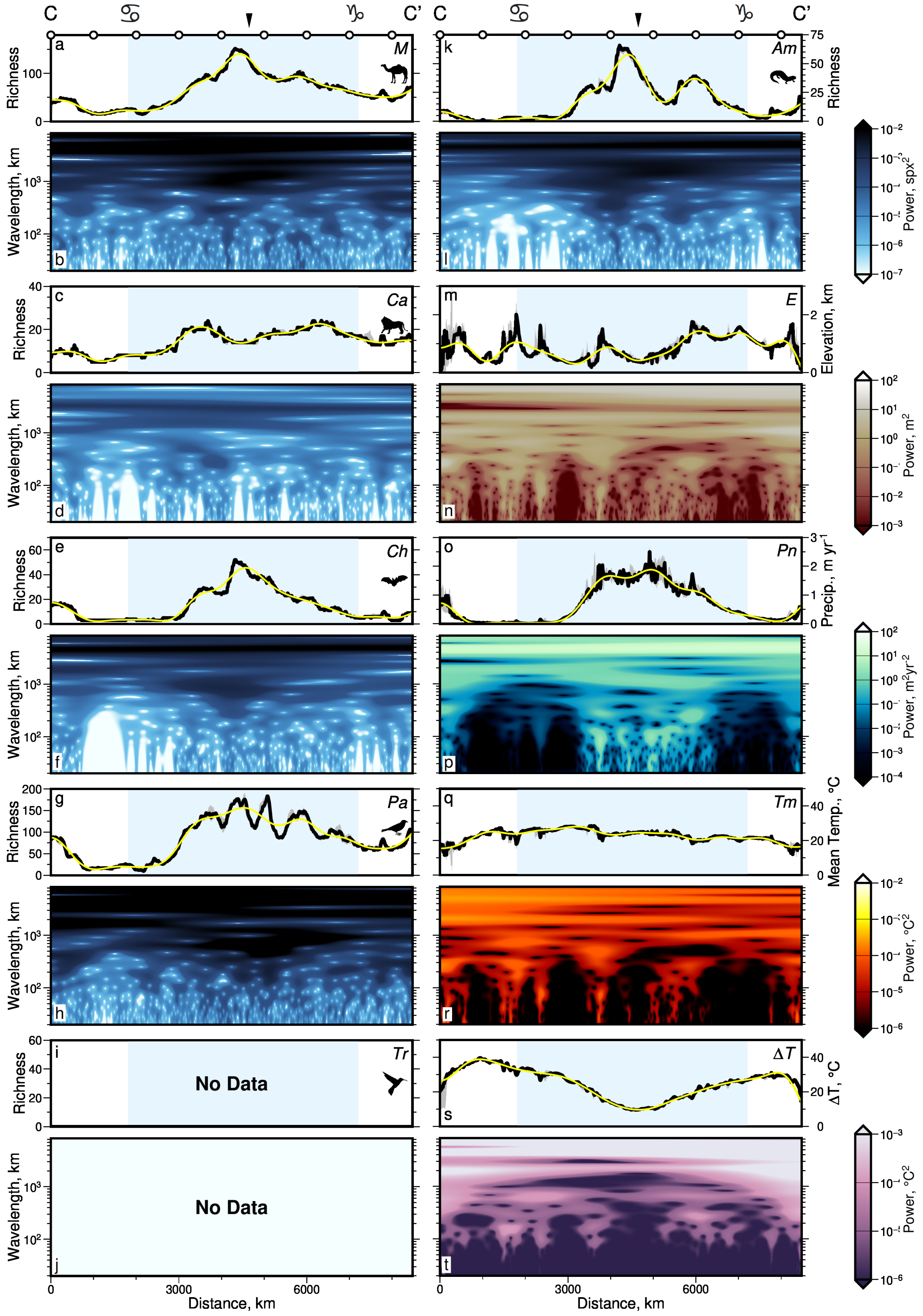
C
C'
a
M
Richness
b
Wavelength, km
c
Ca
d
e
Ch
f
g
Pa
h
i
Tr
No Data
j
No Data
Distance, km
k
Am
l
Power, spx$^2$
m
E
Elevation, km
n
Power, m$^2$
o
Pn
Precip., m yr$^{-1}$
p
Power, m$^2$yr$^{-2}$
q
Tm
Mean Temp., °C
r
Power, °C$^2$
s
ΔT
ΔT, °C
t
Power, °C$^2$

**Figure 5**

**Figure 5** ***(preceding page)*****: Spectral analyses of African species richness and environmental variables.** As Figure 4 but for transect C—C′ (Africa). Symbols as in Figure 1 of main text. Panels (i)–(j) empty due to absence of Trochilidae in the region. Mean differences between signals and inverse transforms filtered to remove wavelengths $< 1000$ km $= 3.6$ spx ($Ma$), 0.8 spx ($Ca$), 1.7 spx ($Ch$), 8.7 spx ($Pa$), 2.2 spx ($Am$), 0.14 km ($E$), 0.07 m/yr ($Pn$), 0.8 °C ($Tm$), and 0.9 °C ($\Delta T$).

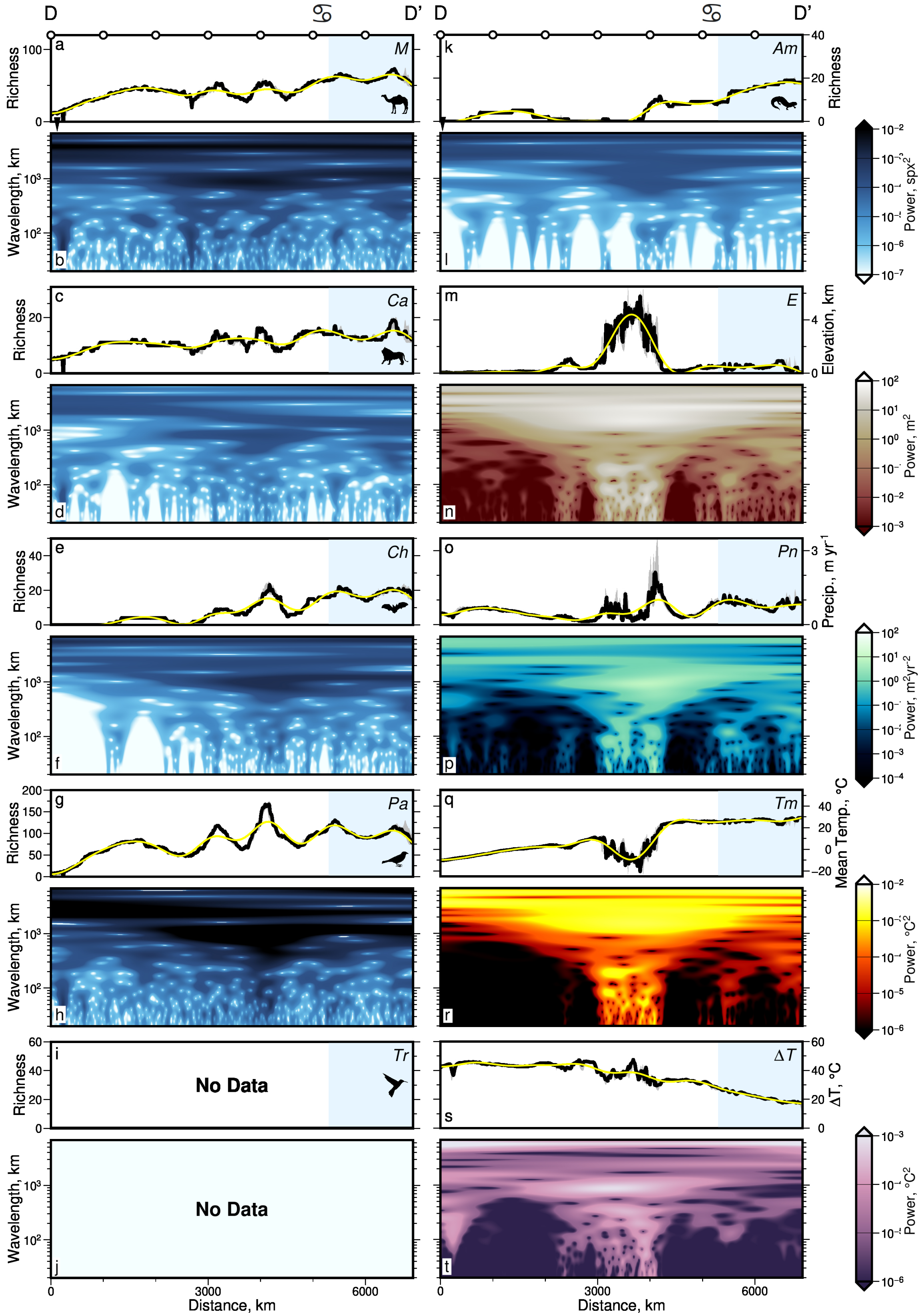

D
D'
Richness
Wavelength, km
Distance, km
Power, spx²
Power, m²
Power, m²yr⁻²
Power, °C²
M
Ca
Ch
Pa
Tr
Am
E
Pn
Tm
ΔT
Elevation, km
Precip., m yr⁻¹
Mean Temp., °C
ΔT, °C
No Data
No Data


**Figure 6**

**Figure 6** ***(preceding page)*****: Spectral analyses of Eurasian species richness and environmental variables.** As Figure 4 but for transect D—D′ (Eurasia). Panels (i)–(j) empty due to absence of Trochilidae in the region. Mean differences between signals and inverse transforms filtered to remove wavelengths $<$ 1000 km = 2.6 spx ($Ma$), 0.9 spx ($Ca$), 1.4 spx ($Ch$), 6.2 spx ($Pa$), 0.7 spx ($Am$), 0.19 km ($E$), 0.10 m/yr ($Pn$), 1.3 °C ($Tm$), and 1.1 °C ($\Delta T$).

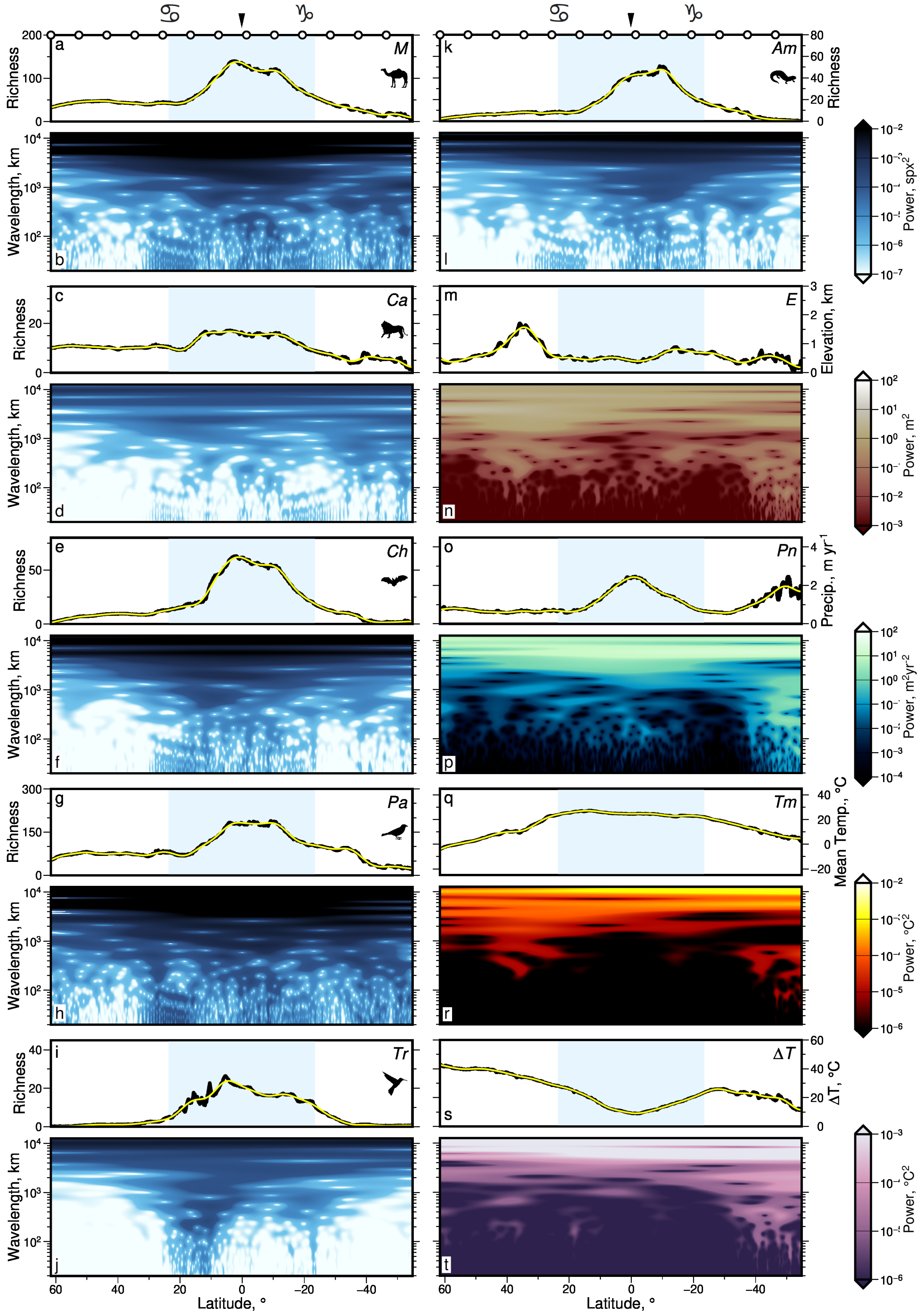

Richness
Wavelength, km
Latitude, °
M
Am
Ca
E
Ch
Pn
Pa
Tm
Tr
ΔT
Elevation, km
Precip., m yr-1
Mean Temp., °C
ΔT, °C
Power, m2
Power, °C2


**Figure 7**

**Figure 7** ***(preceding page)*****: Spectral analyses of global averages of species richness and environmental variables.** As Figure 4 but for global latitudinally-averaged transects. Mean differences between signals and inverse transforms filtered to remove wavelengths < 1000 km = 1.2 spx ($Ma$), 0.3 spx ($Ca$), 0.6 spx ($Ch$), 2.3 spx ($Pa$), 0.4 spx ($Tr$), 0.5 spx ($Am$), 0.03 km ($E$), 0.04 m/yr ($Pn$), 0.3 °C ($Tm$), and 0.3 °C ($\Delta T$).

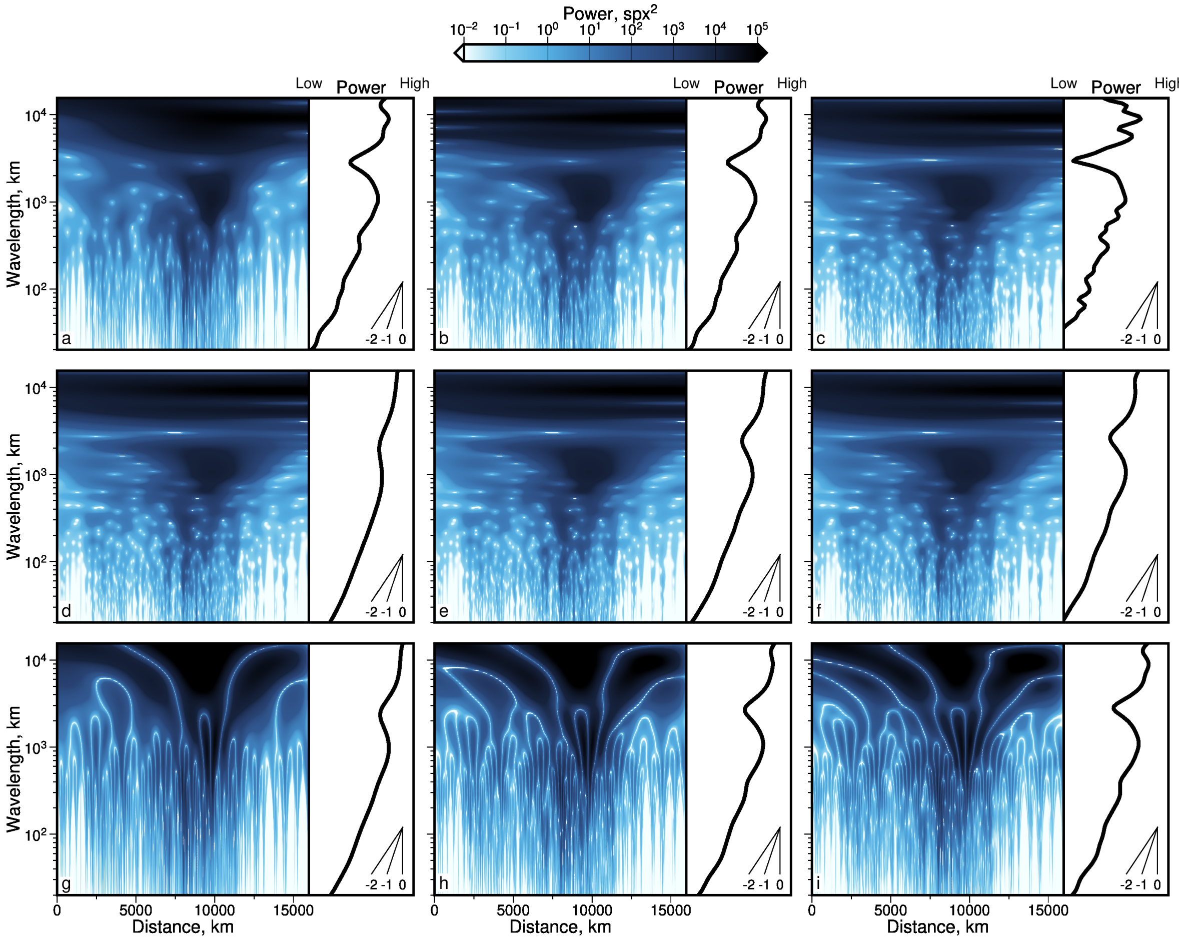


**Figure 8: Testing alternative mother wavelets.** Continuous wavelet transforms of species richness of Amphibia along transect A—A′, using different mother wavelets. (a) Mother wavelet = Morlet, $\omega_0 = 4$; see Materials and Methods in main text, Torrence & Compo (1998). Side panel = distance-averaged power as a function of wavelength, across entire transect; see inset slope guide for spectral slopes of −2, −1 and 0. (b) As (a) but $\omega_0 = 6$, i.e. same as main text Figure 2l. (c) As (a) but $\omega_0 = 8$. (d) As (a) but mother wavelet = Paul, order $m = 2$. (e) As (d) but $m = 4$. (f) As (d) but $m = 6$. (g) As (a) but mother wavelet = derivative of Gaussian, order $m = 2$. (h) As (g) but $m = 4$. (i) As (g) but $m = 6$.

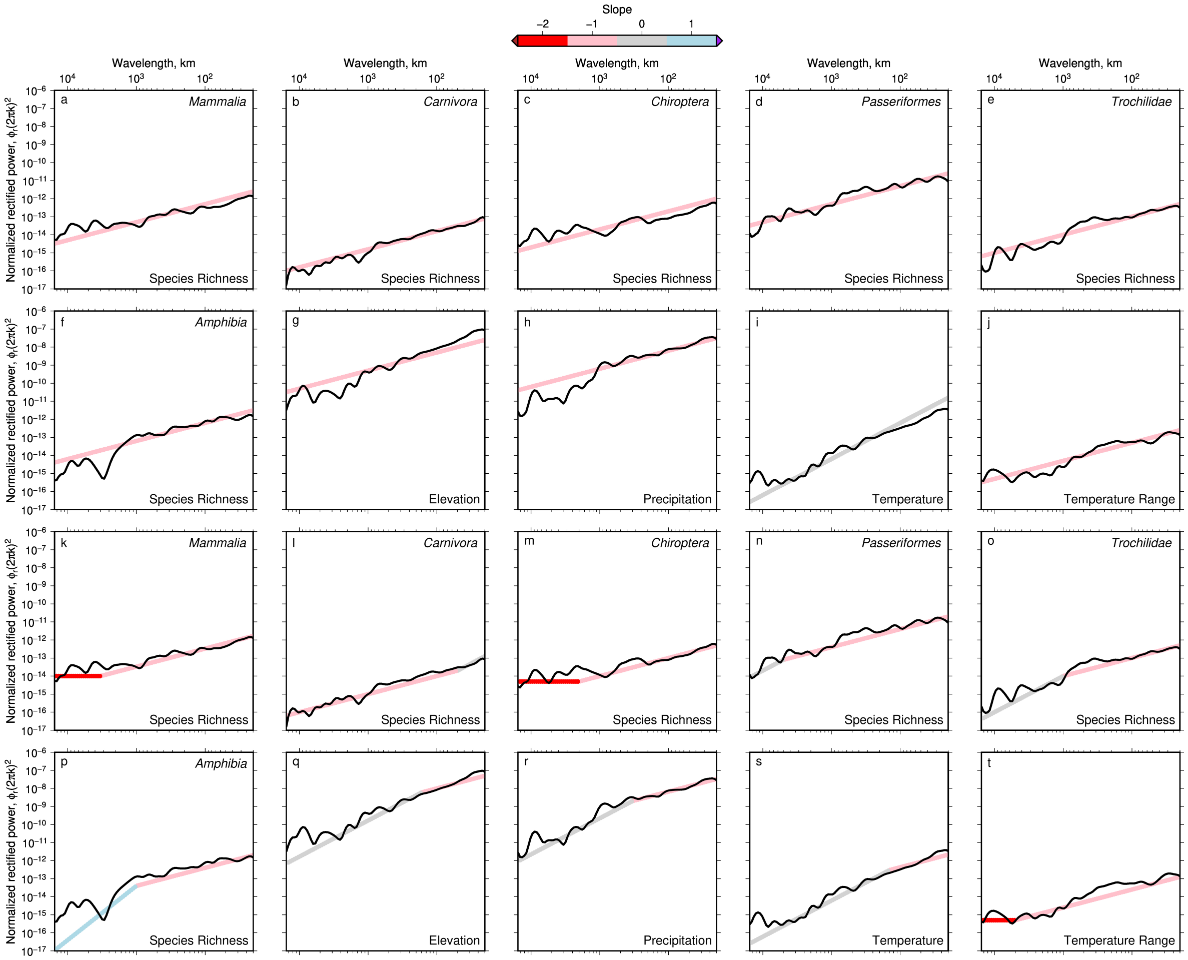


**Figure 9: Fitting theoretical integer spectral slopes to distance-averaged power spectra of observed American series.** (a) Black line = rectified distance-averaged power of Mammalia species richness along transect A—A′ (Jenkins *et al.*, 2013), i.e. distance average of Figure 2a of main text, normalized so a slope of −2 is horizontal. Colored band = best-fitting integer power law relationship between species richness power and wavenumber i.e. slope = $\gamma$ where $\phi_r \propto k^\gamma$ (see scale bar atop figure). (b)–(f) As (a) but for power of species richness of Carnivora, Chiroptera, Passeriformes, Trochilidae and Amphibia. (g)–(j) As (a)–(f) but for power of elevation, precipitation rate, temperature and temperature range (data from Amante & Eakins, 2009; Karger *et al.*, 2017). (k)–(t) As (a)–(j) but for best-fitting integer two-slope relationships between power and wavenumber i.e. slopes = $\alpha$, $\beta$ where $\phi_r \propto k^\alpha$ if $k \geq k_x$ and $\phi_r \propto k^\beta$ if $k < k_x$, where $k_x$ is best-fitting crossover wavenumber (see Roberts *et al.*, 2019).

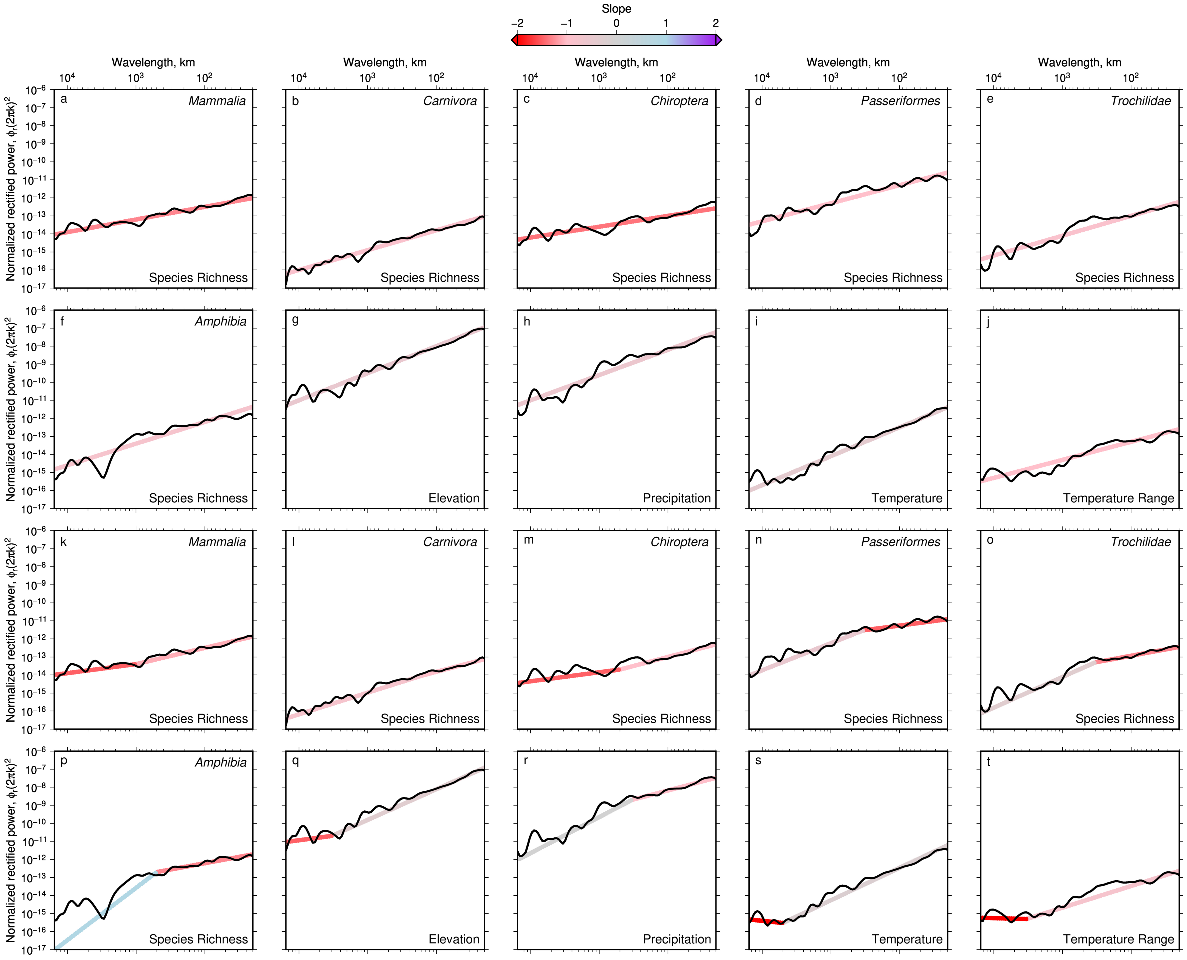


**Figure 10: Fitting theoretical non-integer spectral slopes to distance-averaged power spectra of observed American series.** As Figure 9 but fitted slopes are not restricted to be integers.

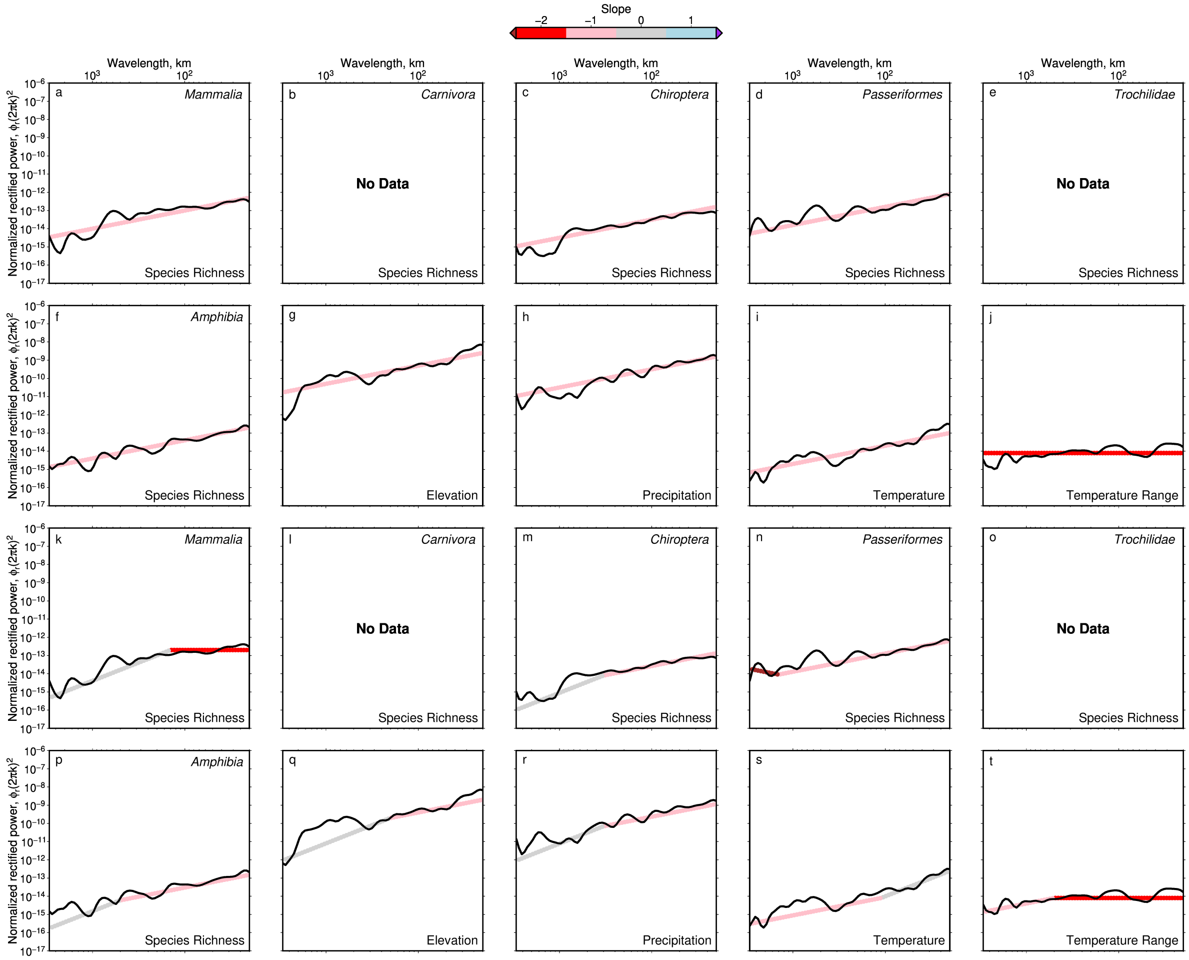


**Figure 11: Fitting theoretical integer spectral slopes to distance-averaged power spectra of observed Australian series.** As Figure 9 but for transect B—B′ (see Figure 4). Panels (b), (e), (l) and (o) empty due to absence of Carnivora and Trochilidae in the region.

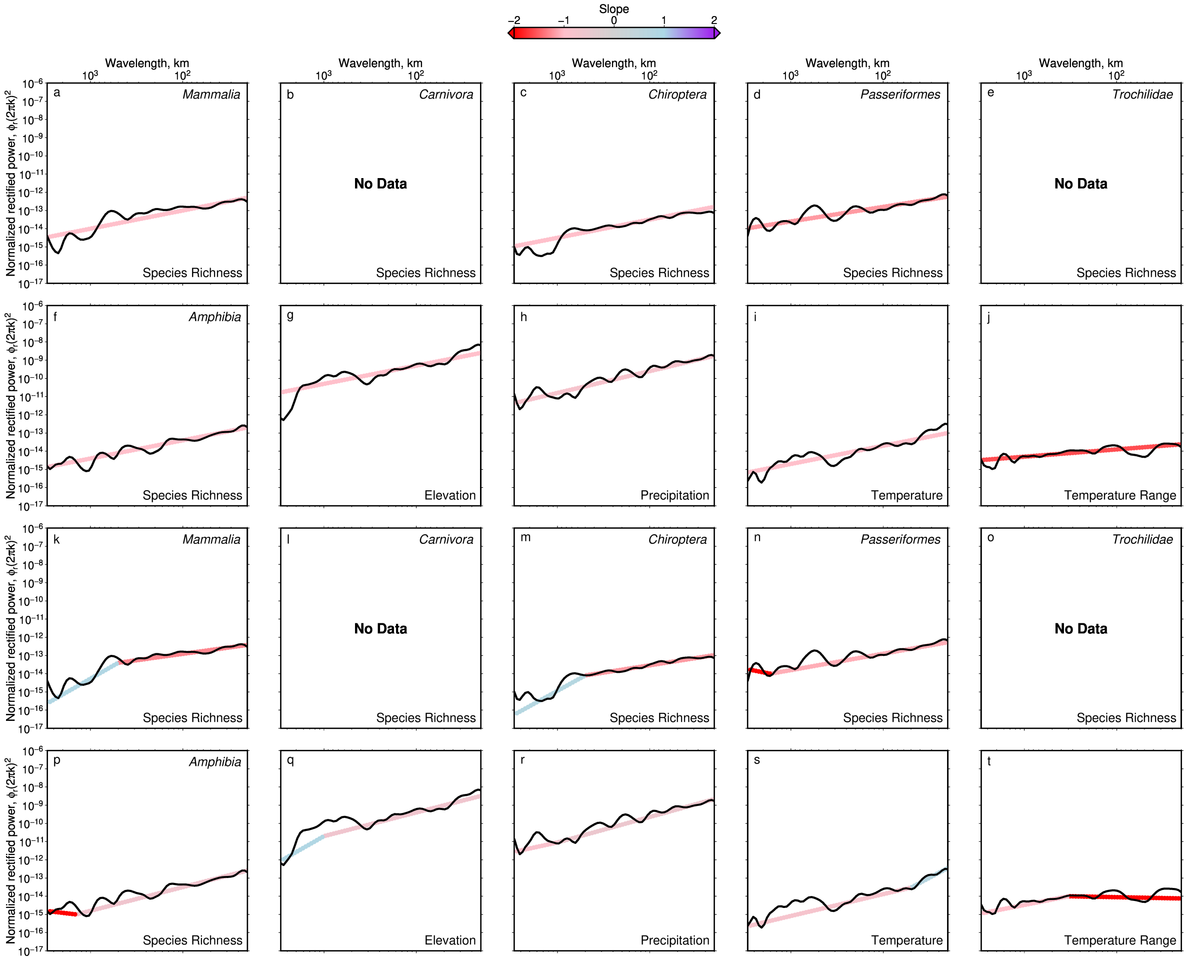


**Figure 12: Fitting theoretical non-integer spectral slopes to distance-averaged power spectra of observed Australian series.** As Figure 11 but fitted slopes are not restricted to be integers.

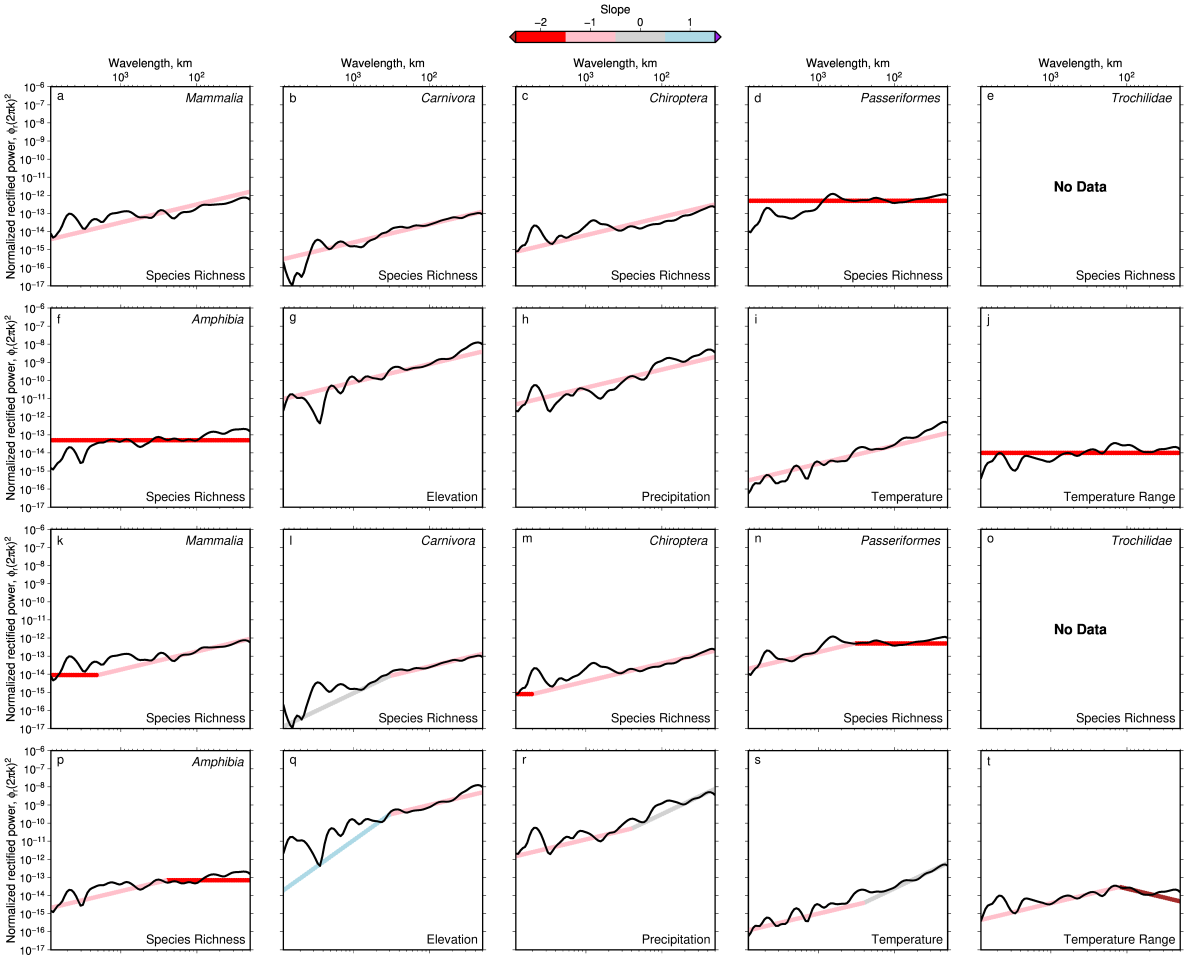


**Figure 13: Fitting theoretical integer spectral slopes to distance-averaged power spectra of observed African series.** As Figure 9 but for transect C—C′ (see Figure 5). Panels (e) and (o) empty due to absence of Trochilidae in the region.

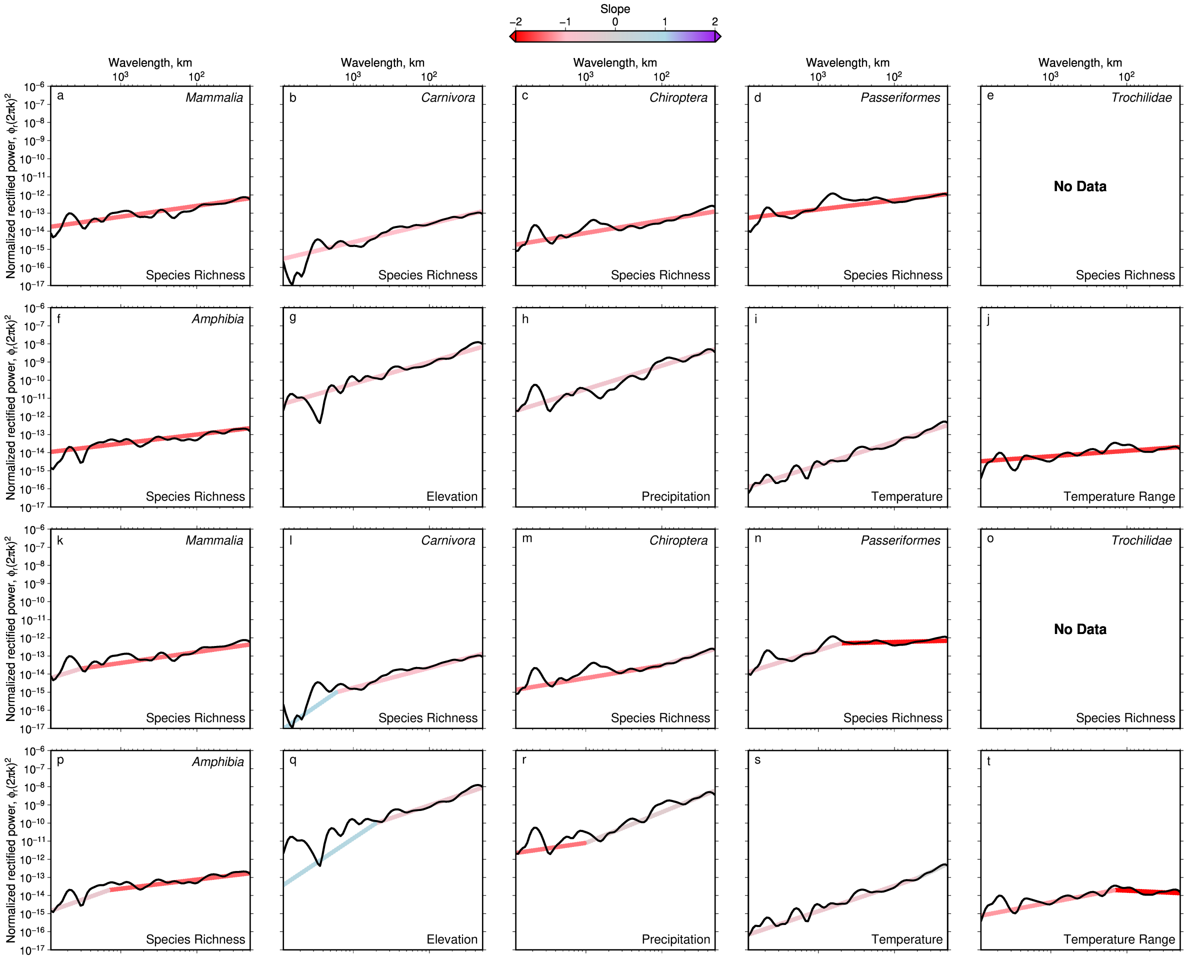


**Figure 14: Fitting theoretical non-integer spectral slopes to distance-averaged power spectra of observed African series.** As Figure 13 but fitted slopes are not restricted to be integers.

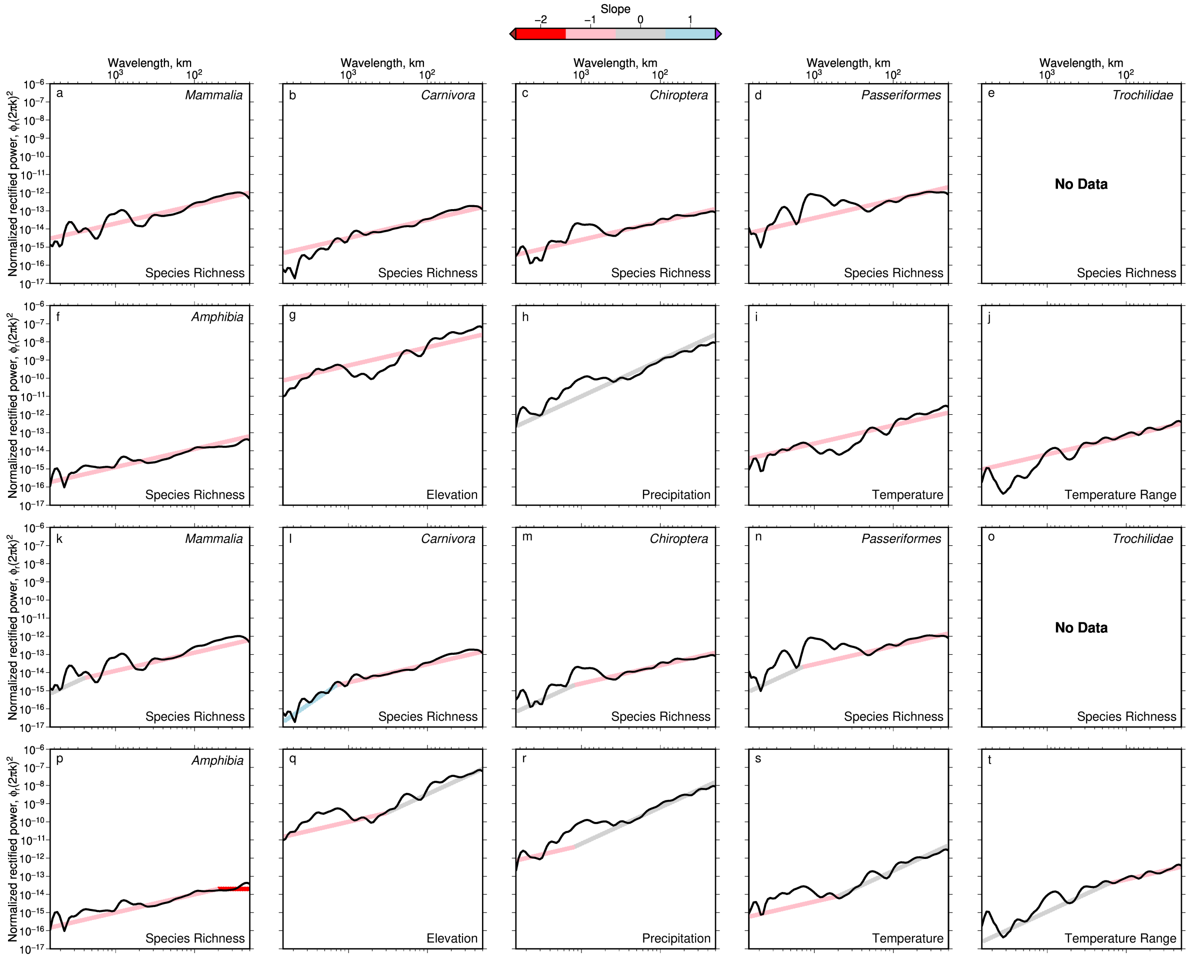


**Figure 15: Fitting theoretical integer spectral slopes to distance-averaged power spectra of observed Eurasian series.** As Figure 9 but for transect D—D′ (see Figure 6). Panels (e) and (o) empty due to absence of Trochilidae in the region.

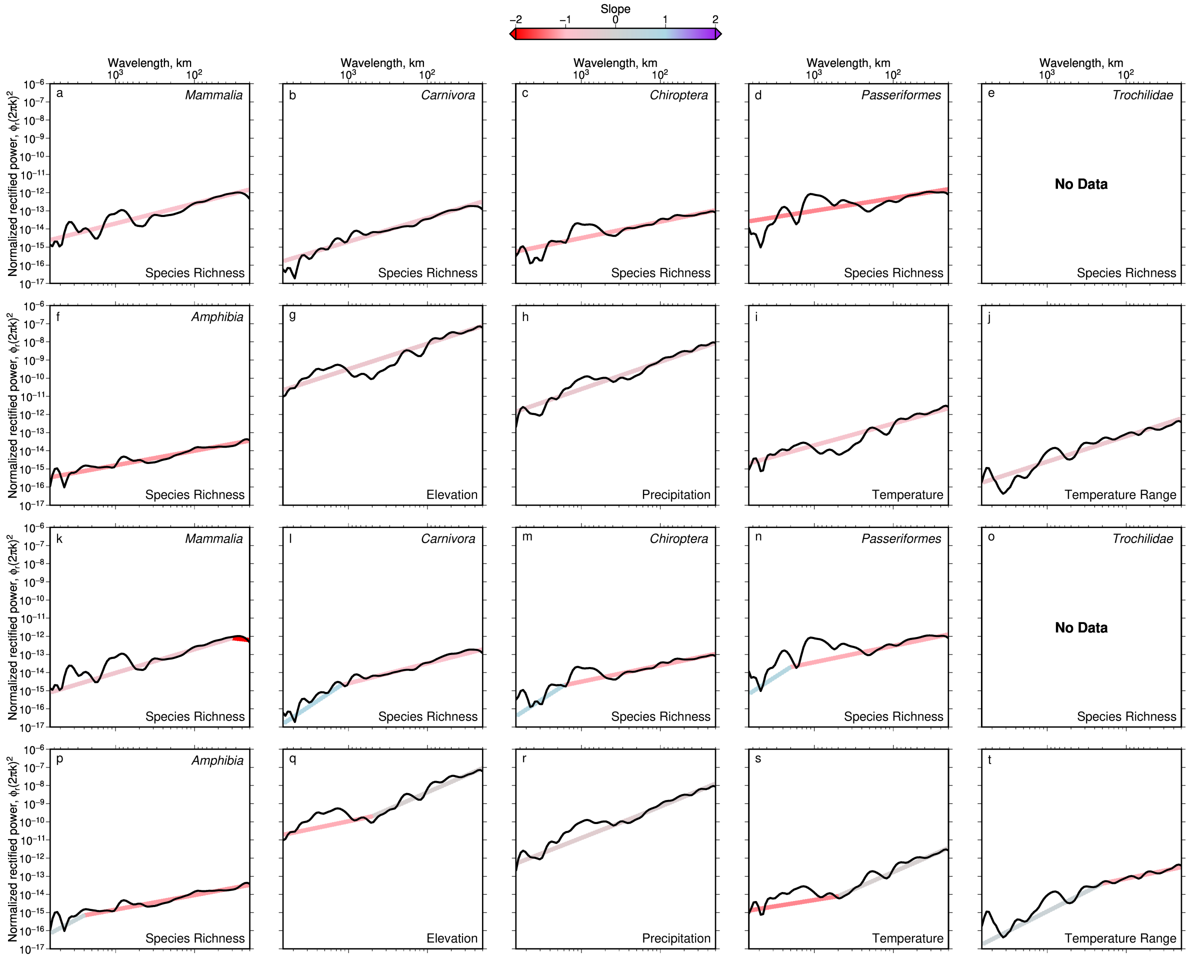


**Figure 16: Fitting theoretical non-integer spectral slopes to distance-averaged power spectra of observed Eurasian series.** As Figure 15 but fitted slopes are not restricted to be integers.

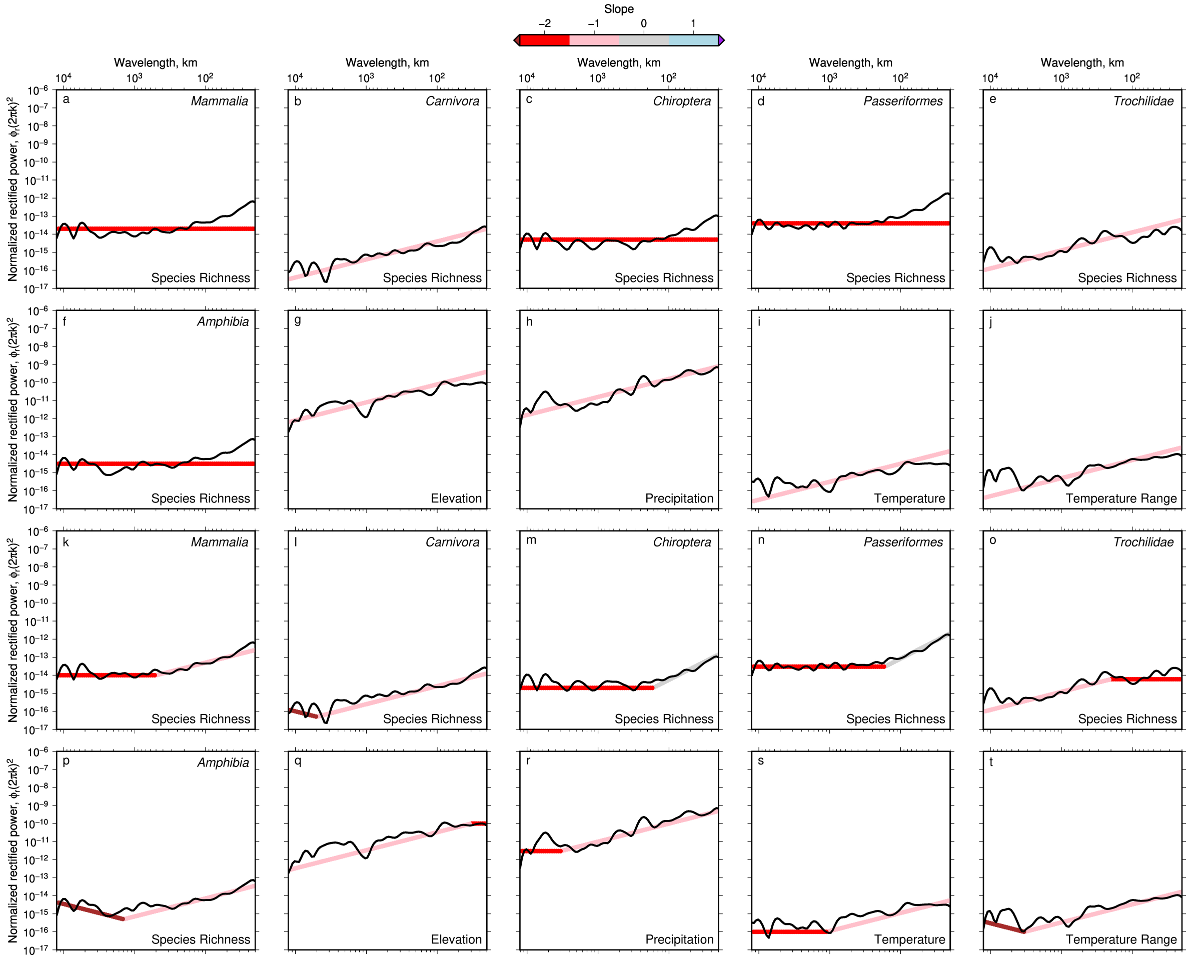


**Figure 17: Fitting theoretical integer spectral slopes to distance-averaged power spectra of observed global series.** As Figure 9 but for latitudinal trend averaged across all longitudes (see Figure 7).

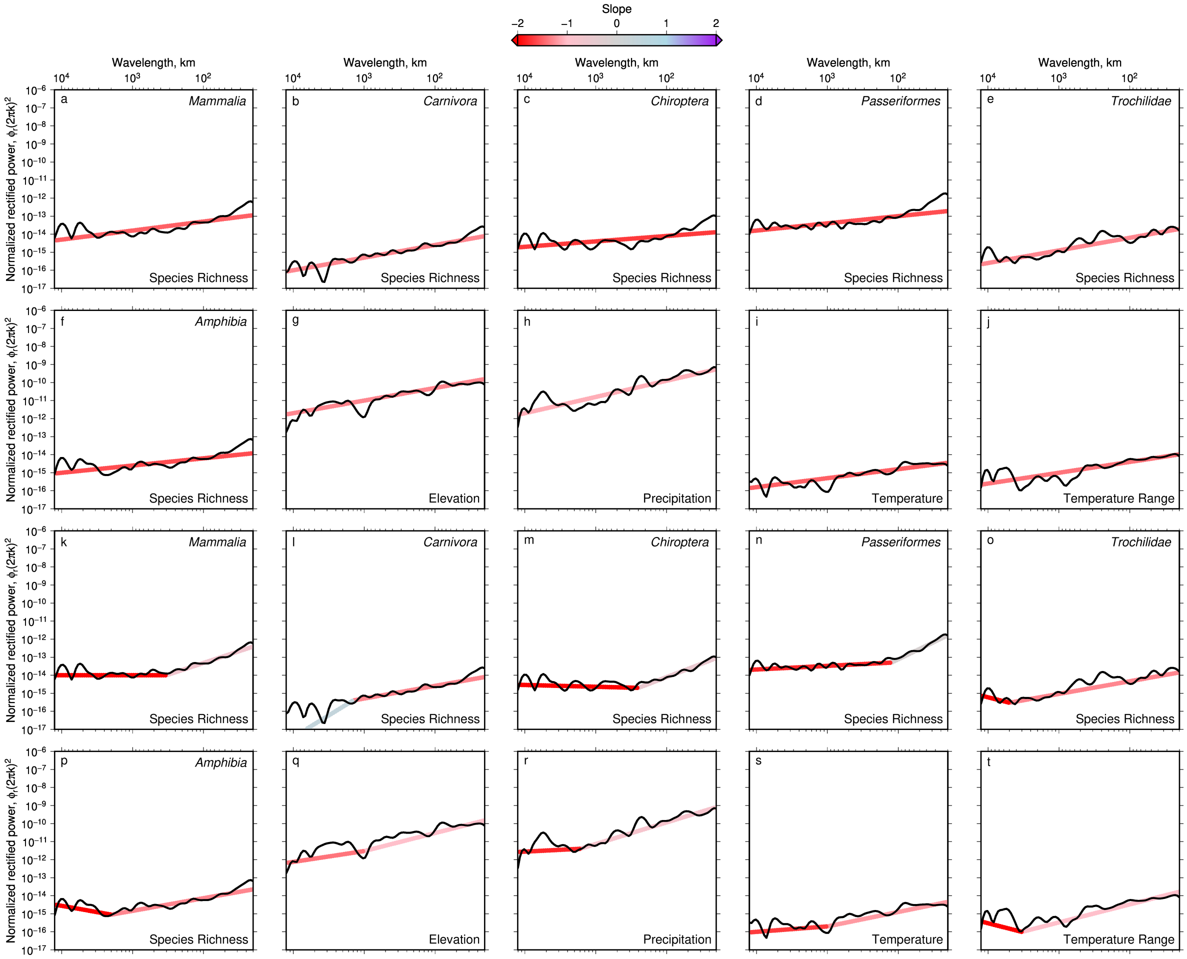


**Figure 18: Fitting theoretical non-integer spectral slopes to distance-averaged power spectra of observed global series.** As Figure 17 but fitted slopes are not restricted to be integers.

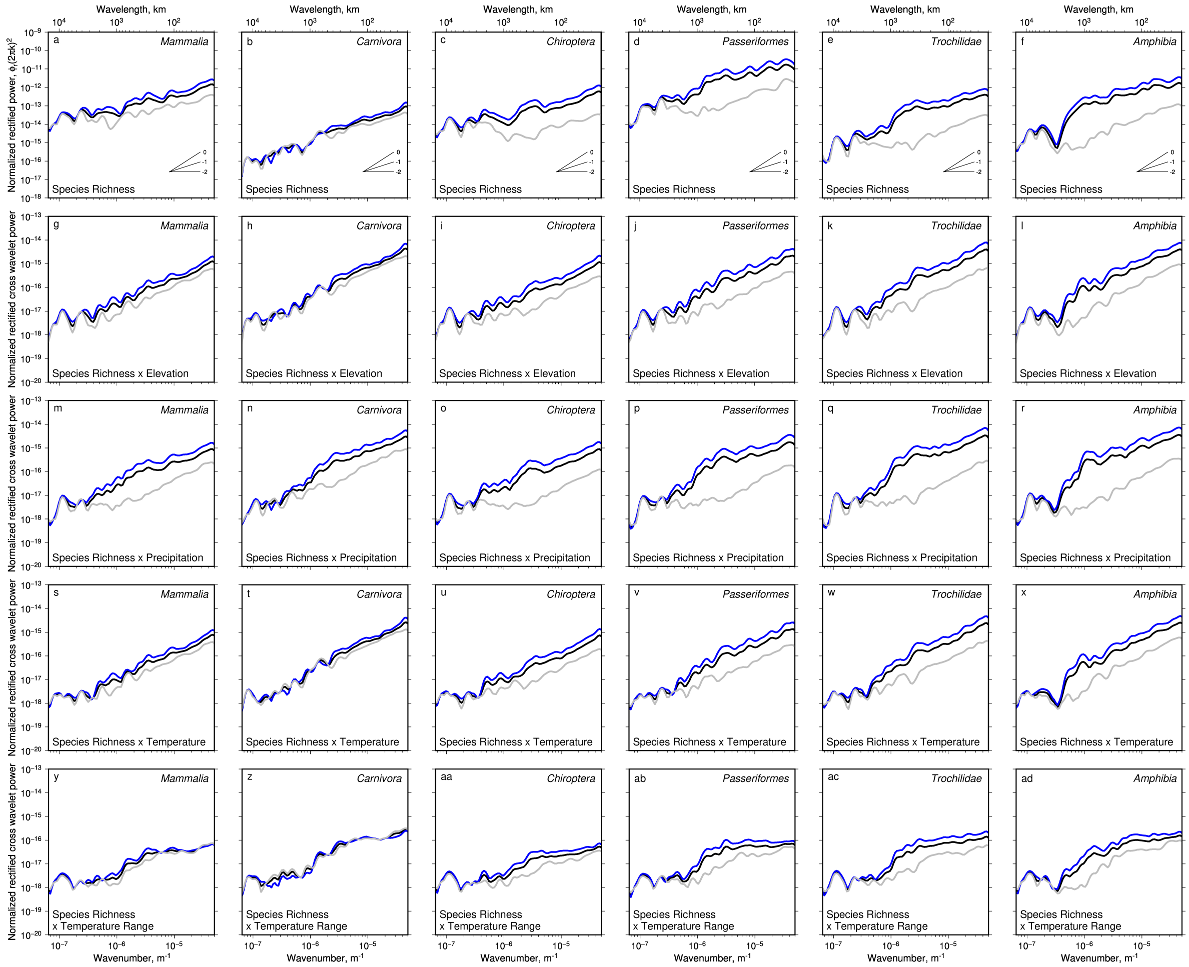


**Figure 19: Comparisons of distance-averaged American spectra within and outside the tropics.** (a) Black line = distance-averaged power from continuous wavelet transform of transect A—A′ (Americas) through species richness of Mammalia (Figure 2a of main text). Blue line = same but only for distance-averaged power of species richness within tropical latitudes. Gray line = same but only for distance-averaged power outside of tropical latitudes. Slopes normalized so that slopes of −2 (red noise) are flat; see inset slope guide for slopes of −1 and 0, i.e. pink and white noise. (b)–(f) As (a) but for species richness of Carnivora, Chiroptera, Passeriformes, Trochilidae and Amphibia. (g) Black line = distance-averaged cross-wavelet power between species richness of Mammalia and elevation. Blue line = same but only for distance-averaged cross power within tropical latitudes. Gray line = same but only for distance-averaged cross power outside of tropical latitudes. Slopes normalized so that slopes of −2 (red noise) are flat. (h)–(l) As (g) but for cross power between elevation and species richness of Carnivora, Chiroptera, Passeriformes, Trochilidae and Amphibia (Amante & Eakins, 2009). (m)–(r) As (g)–(l) but for cross power between mean annual precipitation rate and species richness (Karger *et al.*, 2017). (s)–(x) As (g)–(l) but for cross power between mean annual temperature and species richness (Karger *et al.*, 2017). (y)–(ad) As (g)–(l) but for cross power between mean annual temperature range and species richness (Karger *et al.*, 2017). See body text of main manuscript for discussion of these results.

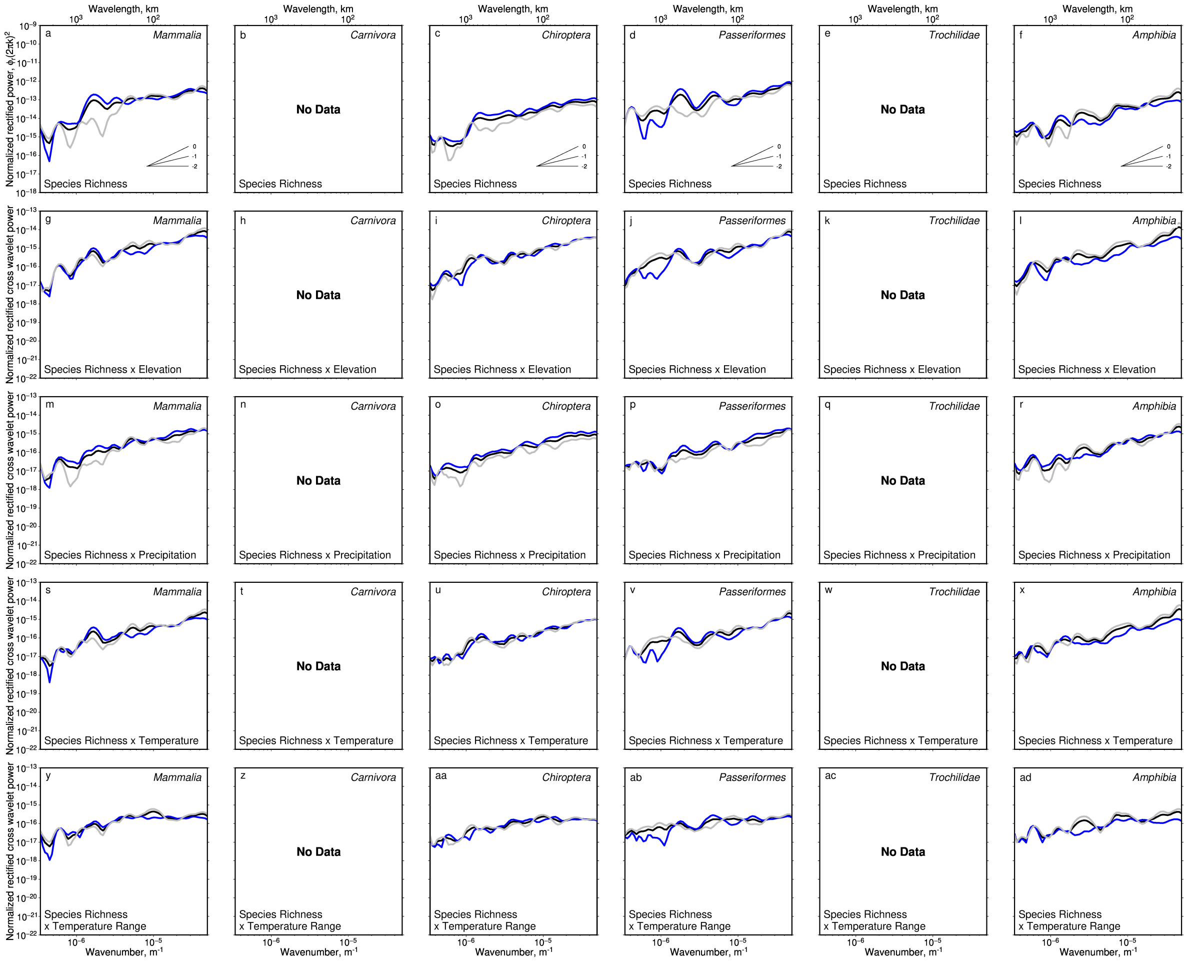


**Figure 20: Comparisons of distance-averaged Australian spectra within and outside the tropics.** As Figure 20 but for transect B—B′ (Australia). Note Carnivora and Trochilidae are absent in Australia.

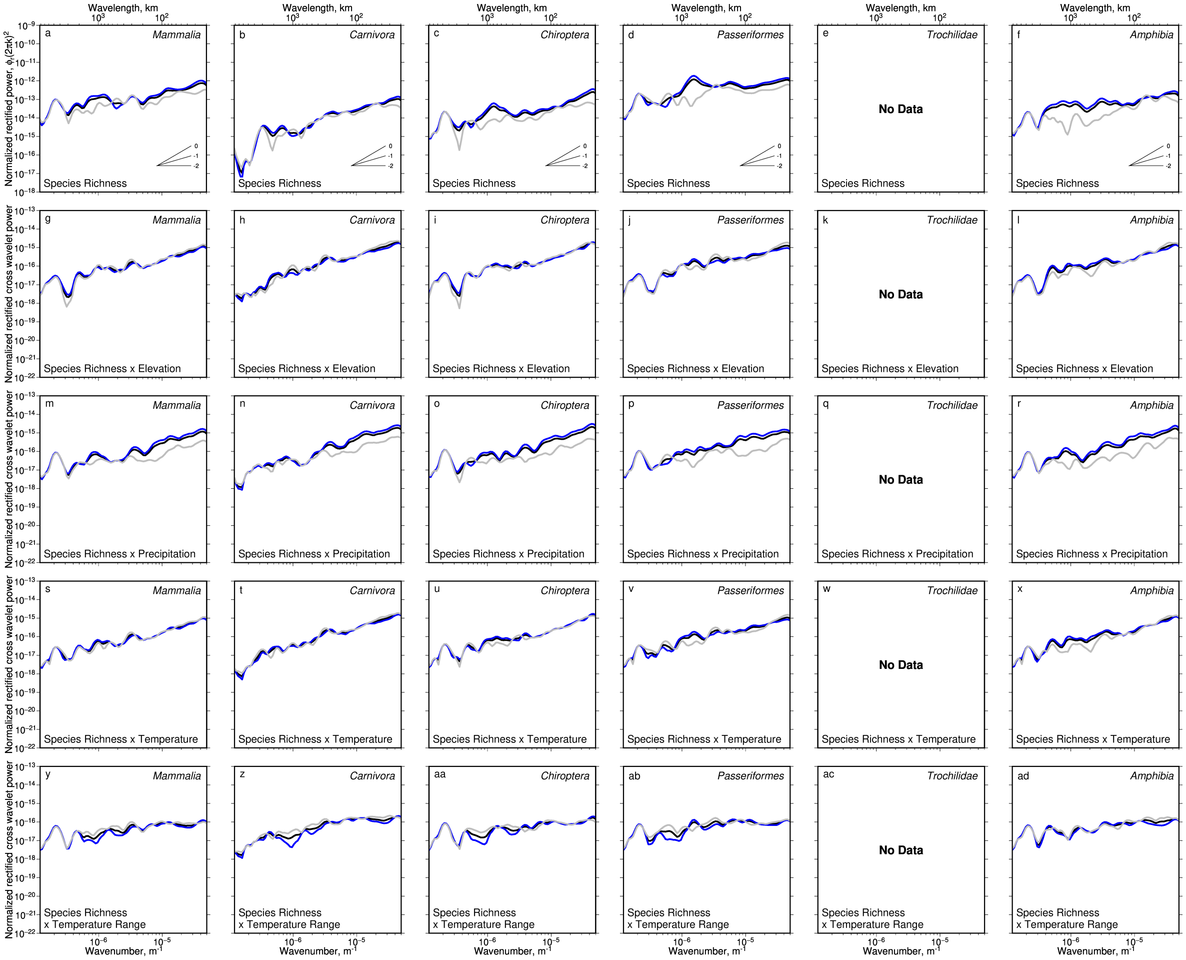


**Figure 21: Comparisons of distance-averaged African spectra within and outside the tropics.** As Figure 20 but for transect C—C′ (Africa). Note Trochilidae are absent in Africa.

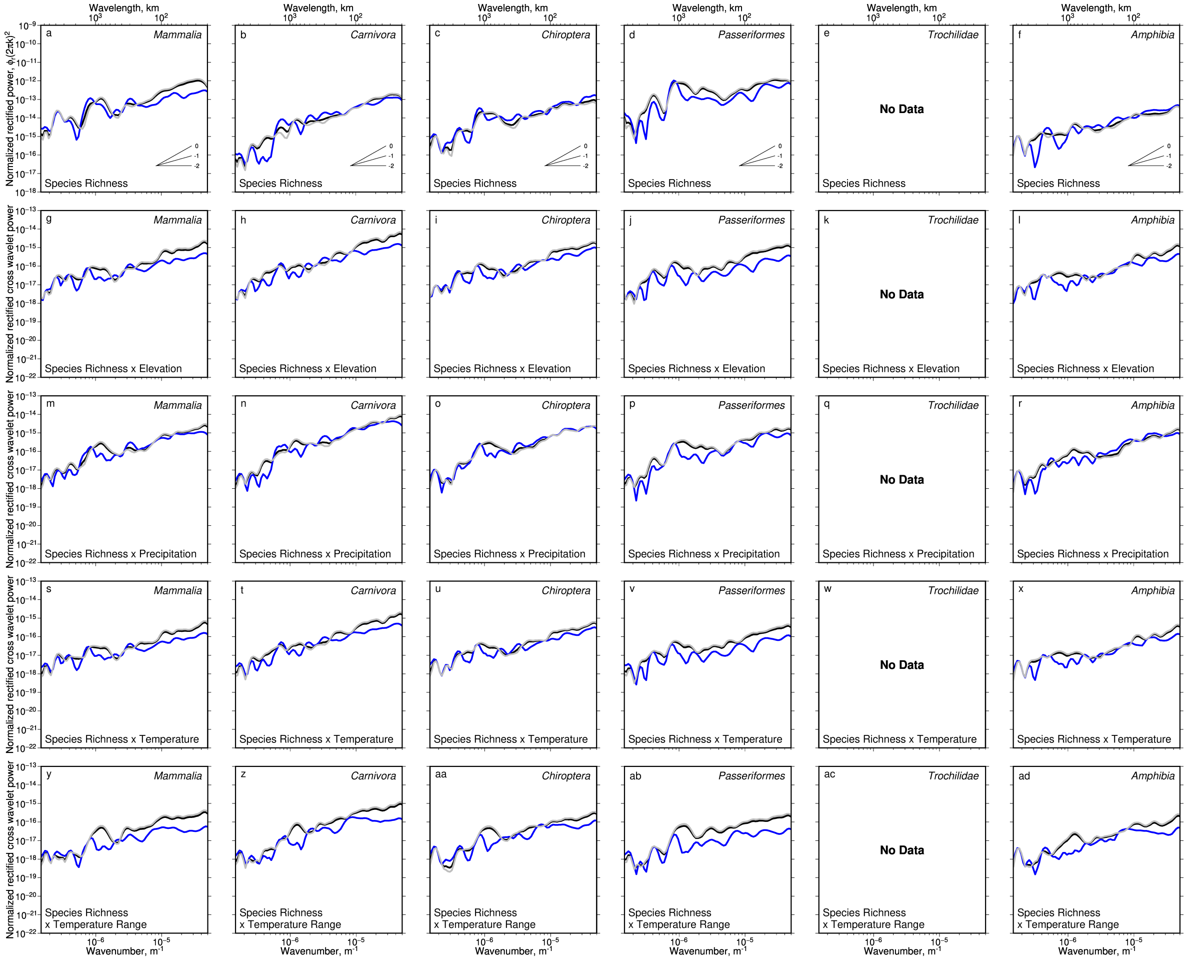


**Figure 22: Comparisons of distance-averaged Eurasian spectra within and outside the tropics.** As Figure 20 but for transect D—D′ (Eurasia). Note Trochilidae are absent in Eurasia.

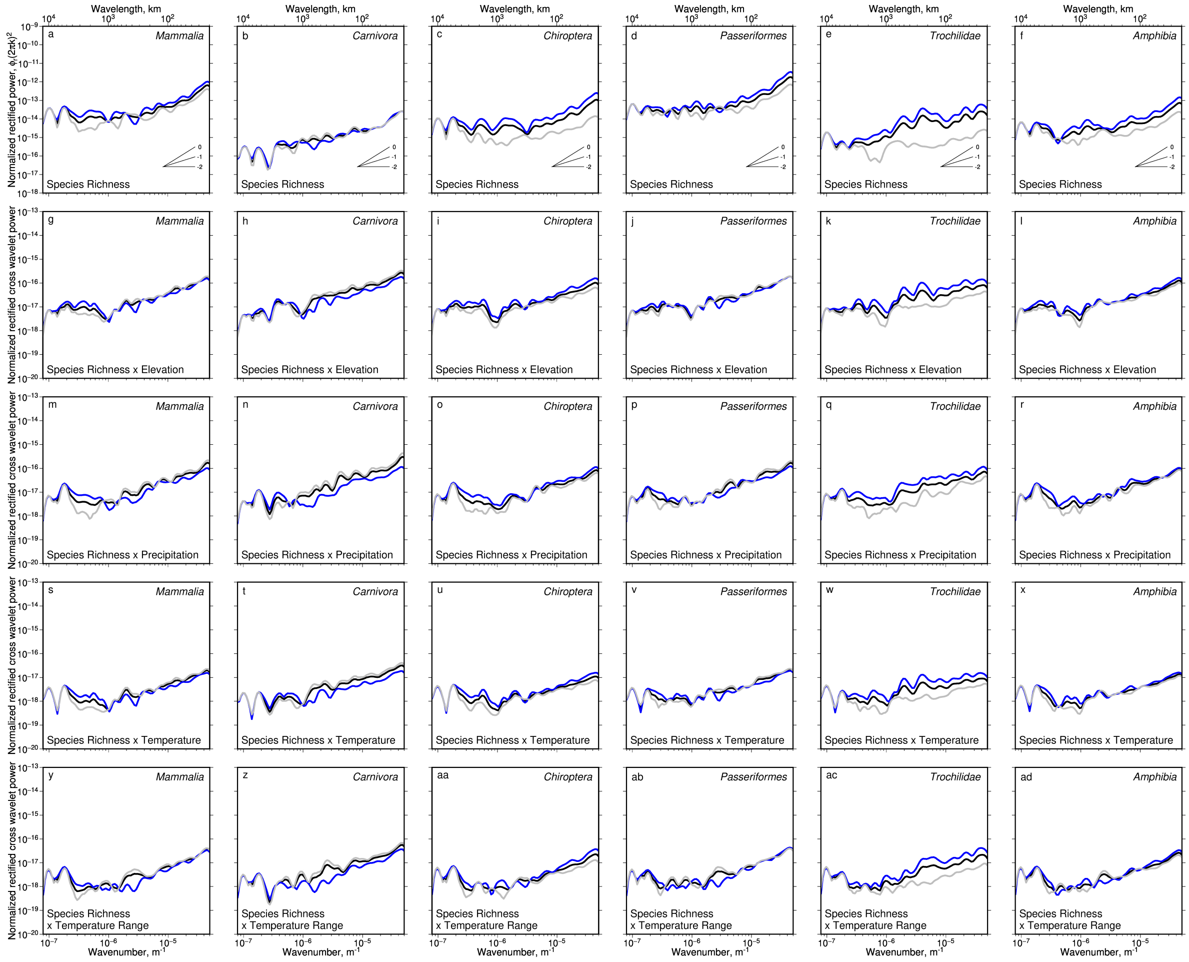

**Figure 23: Comparisons of distance-averaged global spectra within and outside the tropics.** As Figure 20 but for global latitudinal trend averaged across all longitudes. Note that cross-power is calculated between mean global latitudinal transects, rather than being evaluated across multiple individual transects and then averaged.

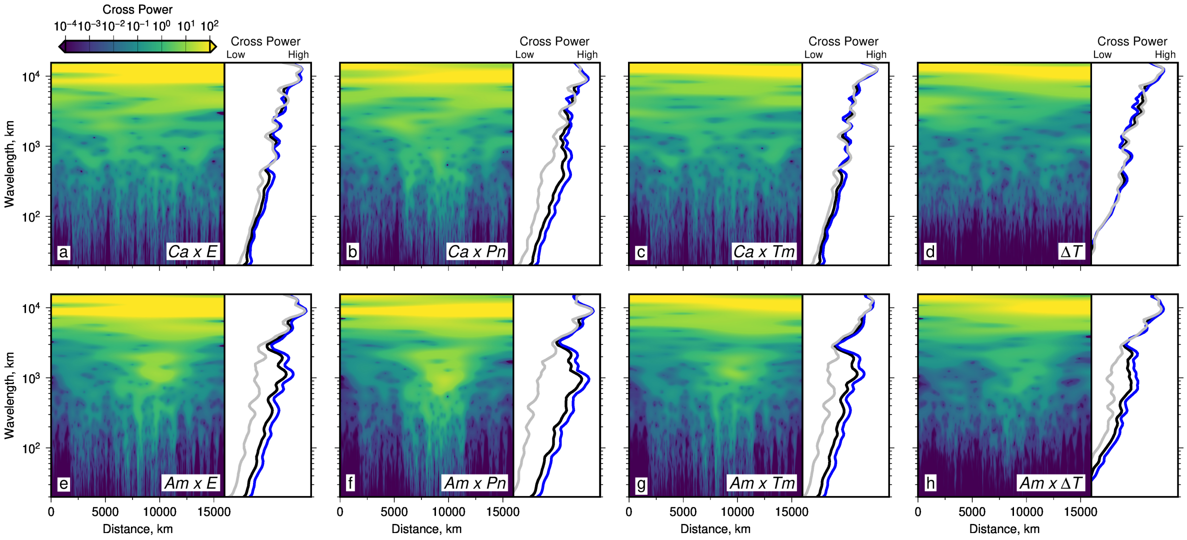


**Figure 24: Unmasked cross wavelet power between American species richness and environment**. As Figure 3 of main text except areas of coherence below the 90% confidence threshold are not masked. $Ca$ = carnivoran richness, $Am$ = amphibian richness, $E$ = elevation, $Pn$ = mean annual precipitation, $Tm$ = mean annual temperature, $\Delta T$ = annual temperature range.

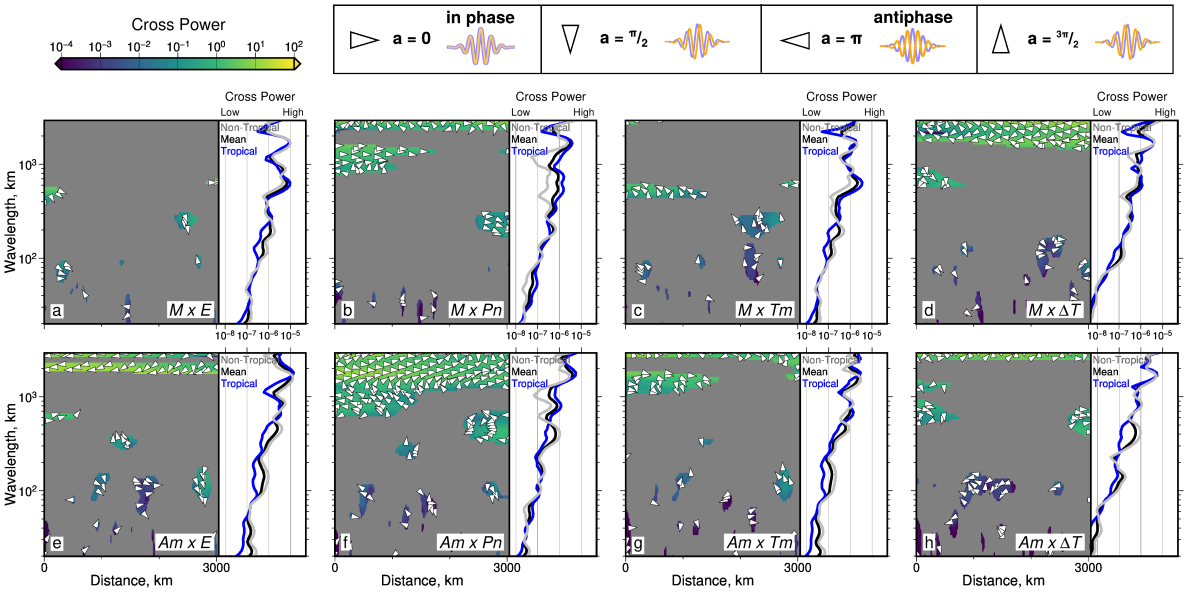


**Figure 25: Cross wavelet power between Australian species richness and environment**. (a) Cross wavelet power between species richness of Mammalia and elevation along section B—B′ (Carnivora are not endemic to Australia). Regions with wavelet coherence below 90% significance level above random red noise are masked in gray. Arrows = direction of phase difference measured clockwise from horizontal, i.e. right-facing = completely in phase, left-facing = completely antiphase (see guide above panel b). Black line in side panel = distance-averaged cross wavelet power; blue line = as black line but for distance-averaged cross wavelet power only within tropical latitudes; gray line = as black line but for distance-averaged cross wavelet power only outside of tropical latitudes. (b)–(d) As (a) but for cross wavelet power between species richness of Mammalia and mean annual precipitation rate, temperature and annual temperature range respectively. (e)–(h) As (a)–(d) but for cross wavelet power between species richness of Amphibia and other variables.

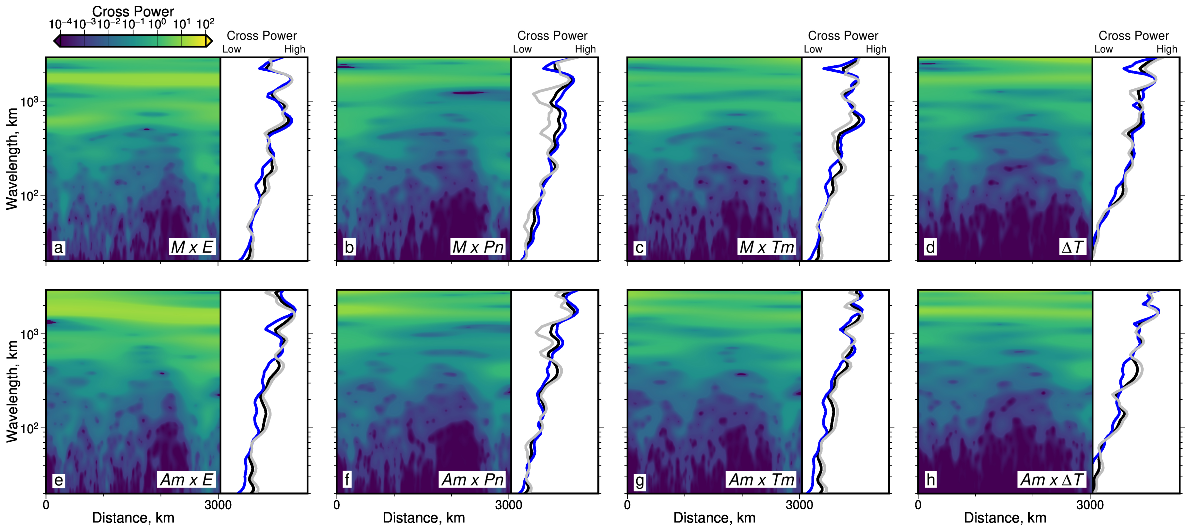


**Figure 26: Unmasked cross wavelet power between Australian species richness and environment**. As Figure 25 except areas of coherence below the 90% confidence threshold are not masked.

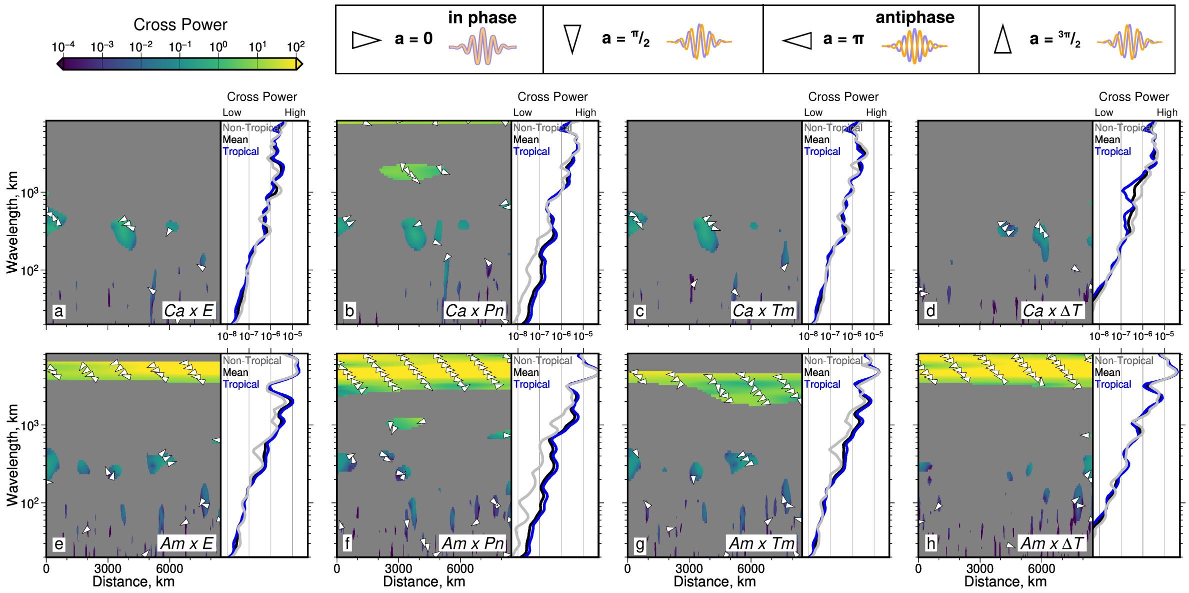


**Figure 27: Cross wavelet power between African species richness and environment**. As Figure 25 but for transect C—C′, and Carnivora species richness replaces Mammalia species richness.

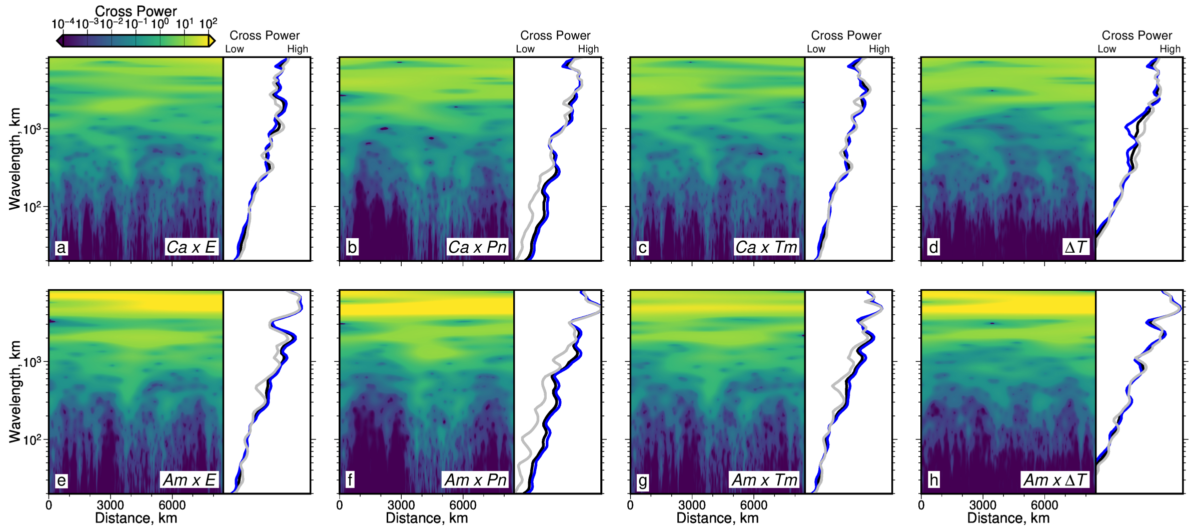


**Figure 28: Unmasked cross wavelet power between African species richness and environment**. As Figure 27 except areas of coherence below the 90% confidence threshold are not masked.

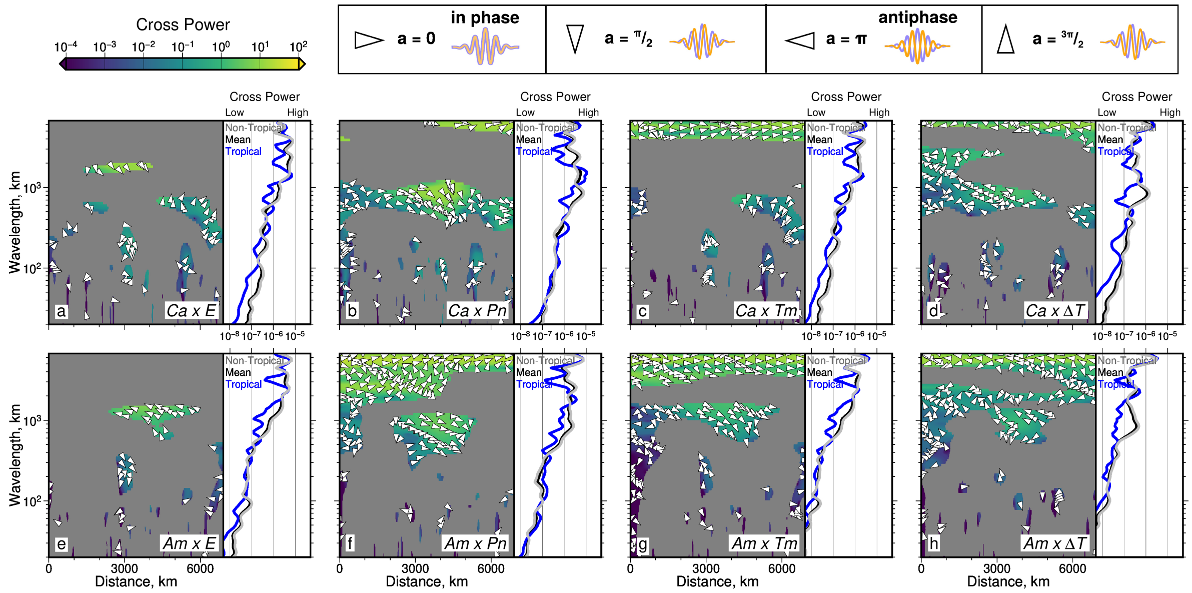


**Figure 29: Cross wavelet power between Eurasian species richness and environment**. As Figure 27 but for transect D—D′.

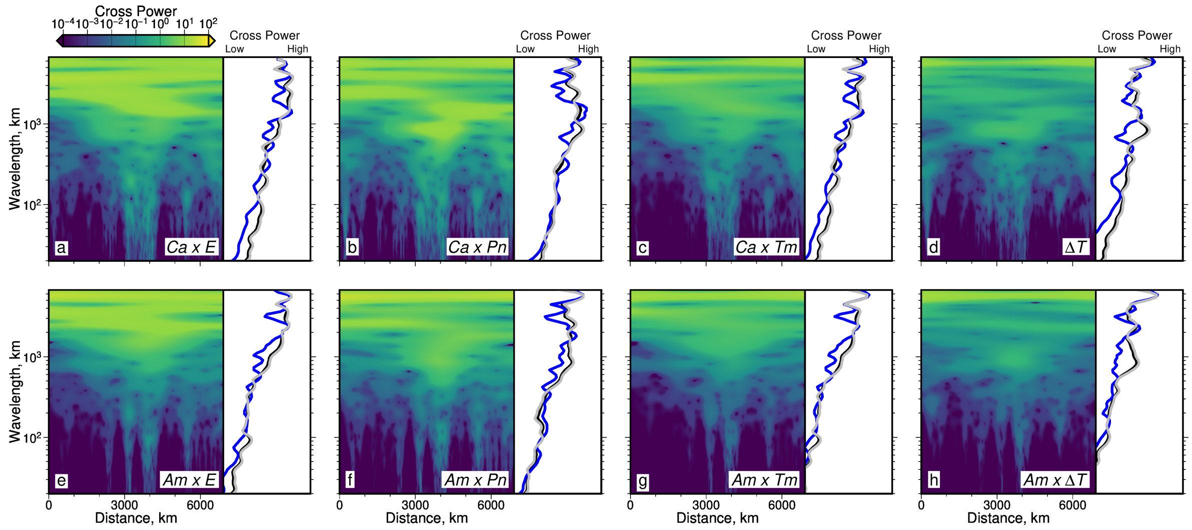

**Figure 30: Unmasked Cross wavelet power between Eurasian species richness and environment**. As Figure 29 except areas of coherence below the 90% confidence threshold are not masked.

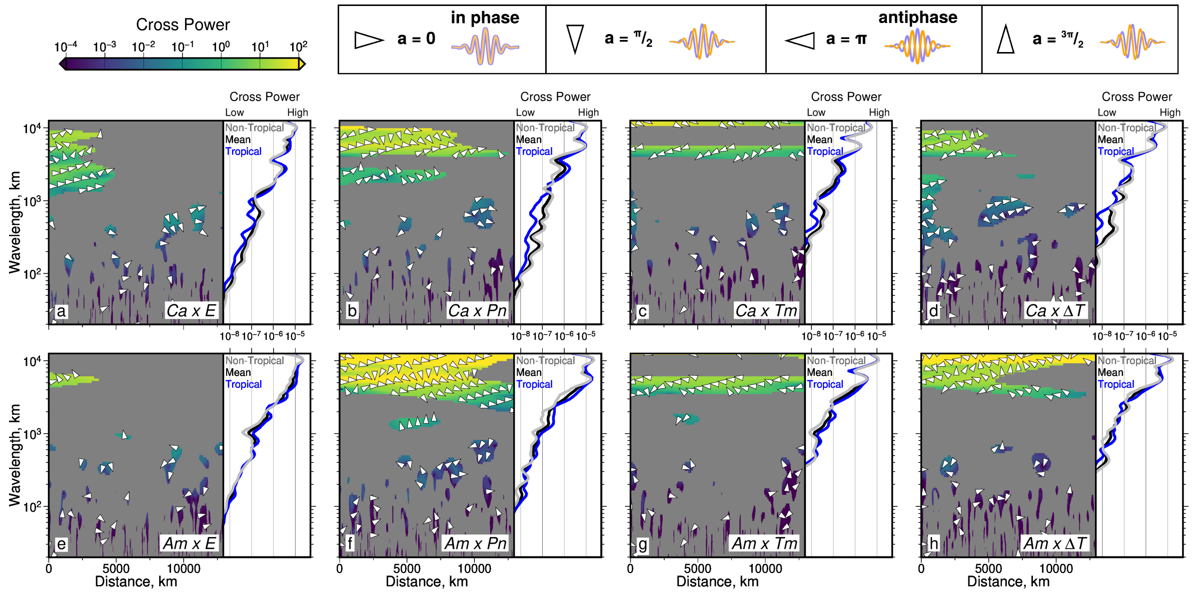

**Figure 31: Cross wavelet power between global species richness and environment**. As Figure 3 of main text but for global latitudinal trends averaged across all latitudes.

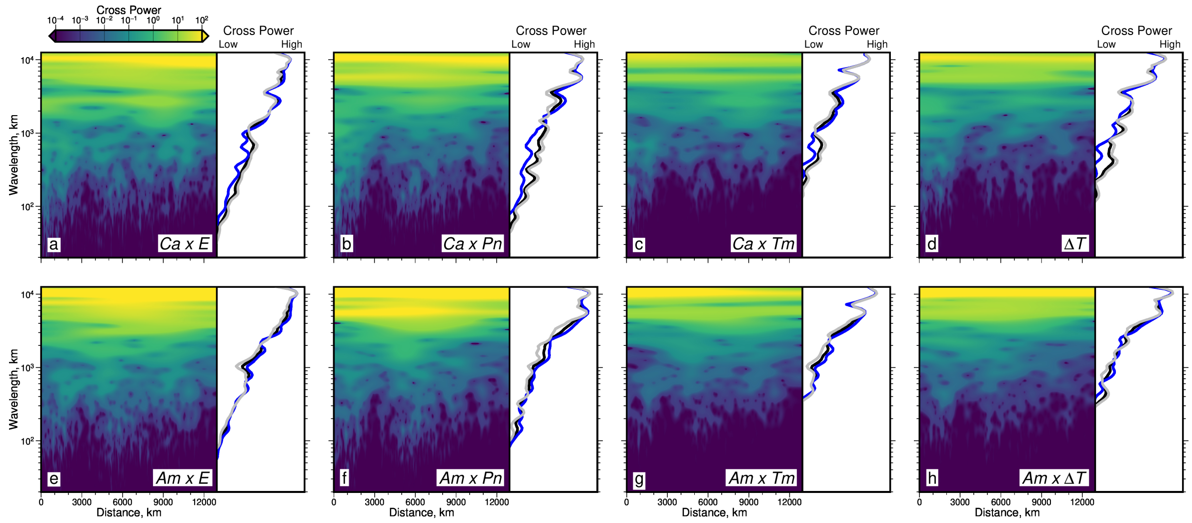


**Figure 32: Unmasked cross wavelet power between global species richness and environment**. As Figure 31 except areas of coherence below the 90% confidence threshold are not masked.

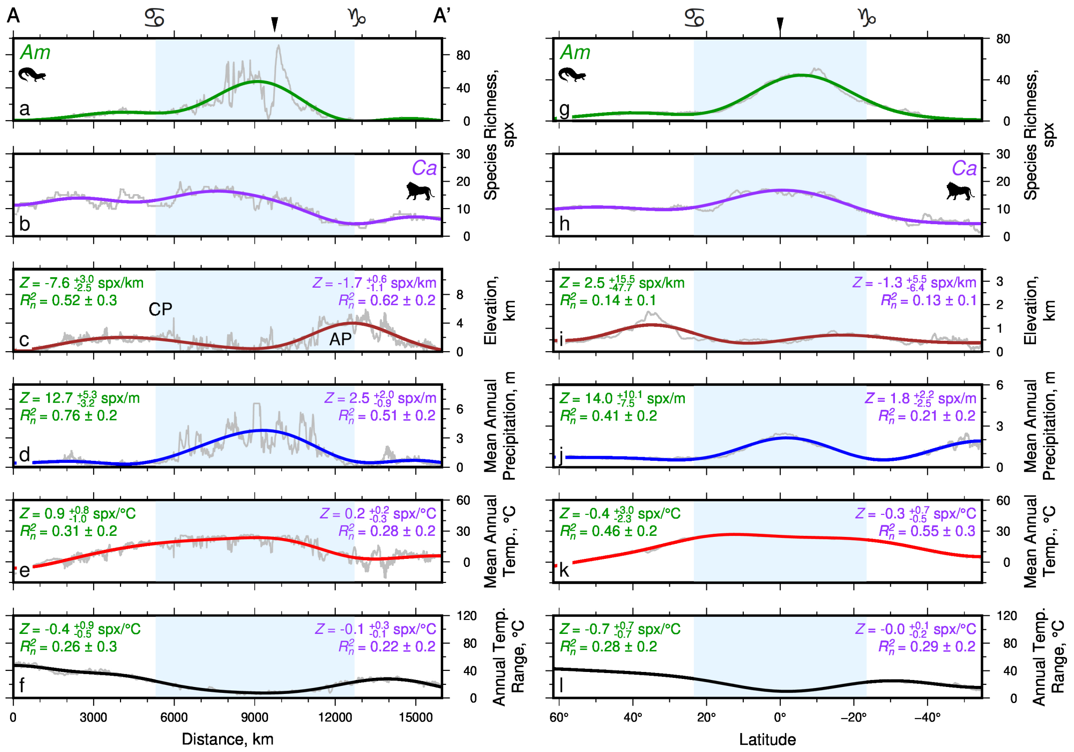

**Figure 33: Coherence and admittance between species richness and environmental variables at large scales**. (a-f) Species richness and environmental variables along Americas transect A—A′ (see Figures 1–2 of main text). Blue band = tropical latitudes; black arrow = Equator; symbols above $x$ axis = Tropics of Cancer and Capricorn. (a) Amphibian species richness. Gray = full-resolution observed species richness trend (see Figure 2k of main text). Green = inverse wavelet transform showing filtered amphibian species richness at wavelengths $\geq$ 3756 km (i.e. one quarter of transect length scale). Mean difference between gray and green lines = 4.4 spx. (b) As (a) but for carnivoran species richness. Mean difference between gray and green lines = 1.2 spx. (c)–(f) As (a)–(b) but for elevation, mean annual precipitation, mean temperature, and annual temperature range, respectively. $Z$, $R_n^2$ = mean admittance and coherence, respectively, between species richness and given environmental variable. Green text = admittance for amphibian species richness; purple text = admittance for carnivoran species richness. (g)–(l) As (a)–(f) but for mean global latitudinal transects.

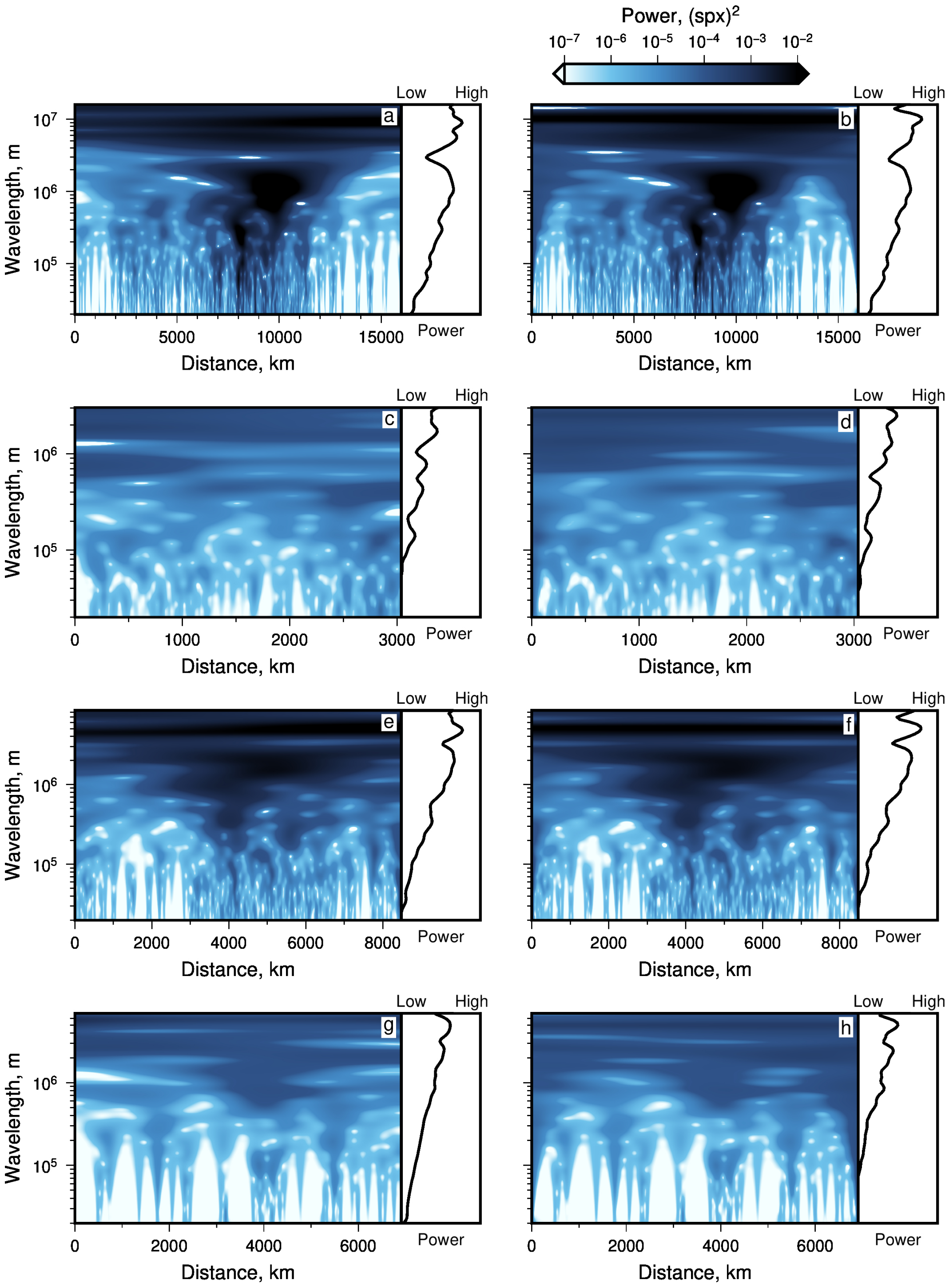


**Figure 34: Effect of mirroring signals to reduce edge effects on wavelet power**. (a) Rectified wavelet power for amphibian species richness signal along A—A′, mirrored to reduce edge effects, as in Figure 2l of main text. Side panel = distance-averaged rectified power. (b) As (a) except signal was not mirrored before wavelet transformation. Note slight increase in power with respect to power spectrum for mirrored signal, at long wavelengths near start ($x = 0$) and end of spectrum. Distance-averaged power is largely unaffected. (c)–(d) As (a)–(b) but for amphibian species richness across Australia (B—B′). (e)–(f) As (a)–(b) but for species richness across Africa (C—C′). (g)–(h) As (a)–(b) but for species richness across Eurasia (D—D′). In all cases distance-averaged power $x$ axes have the same limits.

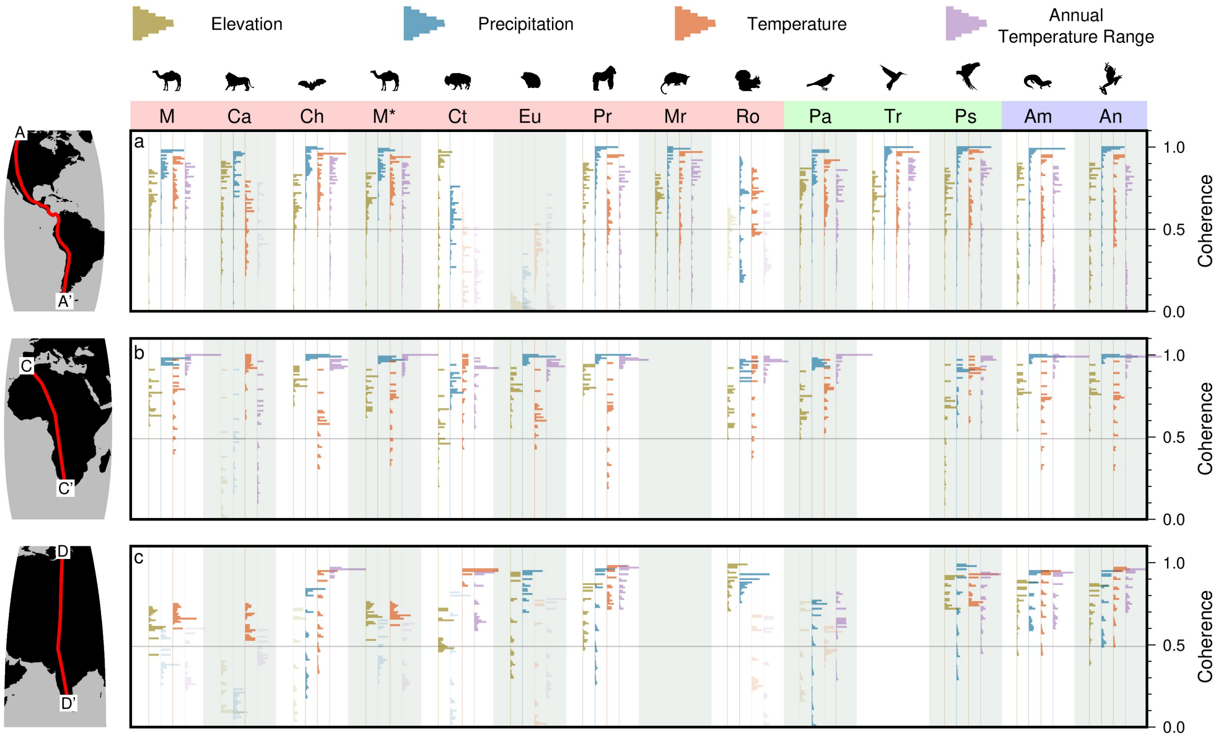


**Figure 35: Effect of mirroring signals to reduce edge effects on coherence with environment.** As Figure 4 of main text except signals were not mirrored before coherence was calculated. (a) Histograms = distribution of coherence values (see right-hand $y$ axis) between species richness and climate, for scales $\geq$3756 km (largest 25% of scales) and different taxa (see top $x$ axis), across the Americas (transect A—A′). Brown/blue/red/purple histograms = coherence between species richness and elevation/mean annual precipitation/mean annual temperature/annual temperature range respectively (see guide above plot; Jenkins *et al.*, 2013; Karger *et al.*, 2017). $M$ = Mammalia, $Ca$ = Carnivora, $Ch$ = Chiroptera, $M^*$ = Mammlia excluding Carnivora, $Ct$ = Cetartiodactyla, $Eu$ = Eulipotyphla, $Pr$ = Primates, $Mr$ = Marsupialia, $Ro$ = Rodentia, $Pa$ = Passeriformes, $Tr$ = Trochilidae, $Ps$ = Psittaciformes, $Am$ = Amphibia, $An$ = Anura. Pink/green/blue bands behind labels cover mammal/bird/amphibian groups respectively. (b)–(c) As (a) but for transects across Africa (C—C′) and Eurasia (D—D′) respectively– see side panel maps for transect locations. Note that compared with Figure 4 of main text, coherence is generally higher "artificially" for un-mirorred signals, due to coherence of edge effects.

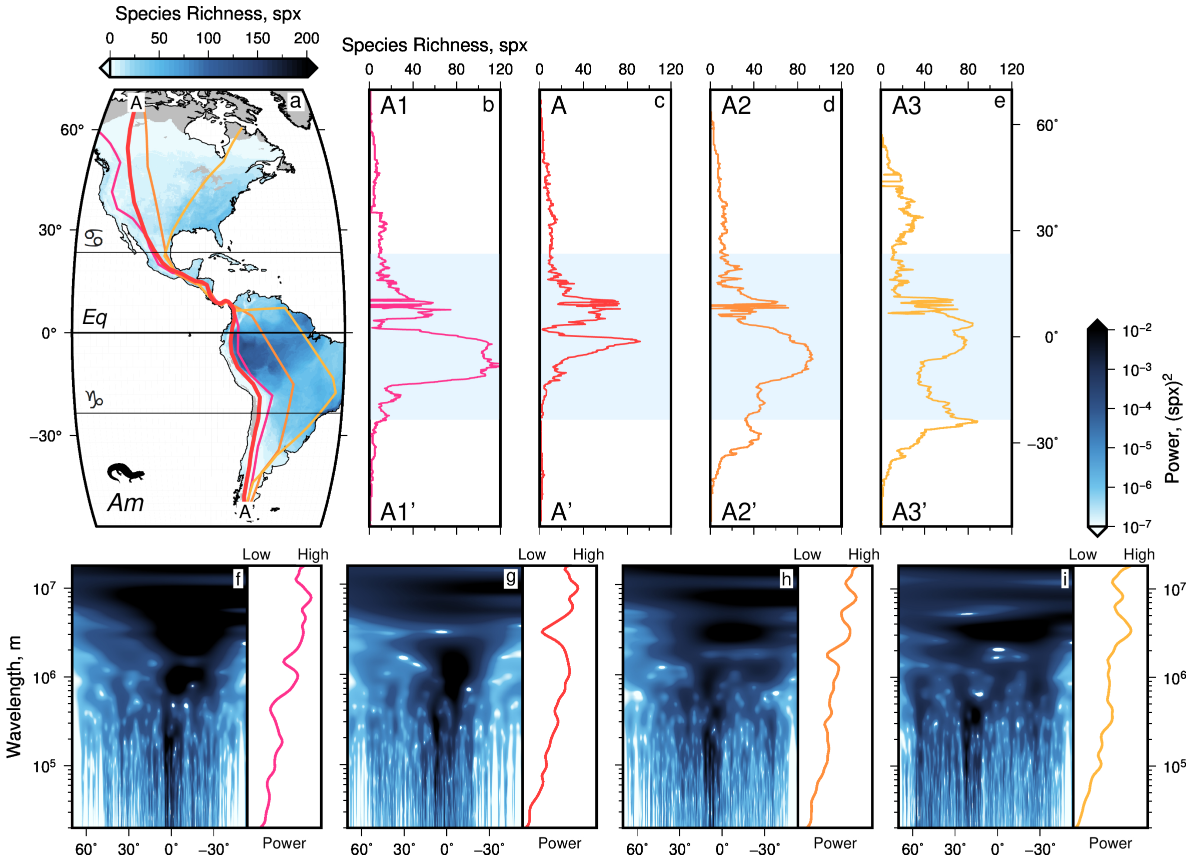


**Figure 36: Analysis of alternative transects through amphibian species richness across the Americas**. (a) Amphibian species richness map (Jenkins *et al.*, 2013). Pink/orange/yellow lines = locations of new transects shown in (b), (d) and (e). Red line = transect A—A′. Eq = Equator. (b)–(e) Amphibian species richness as a function of latitude along each transect. (f)–(i) Rectified wavelet power of species richness, for signals shown in (b)–(e) respectively. Side panel = distance-averaged power.

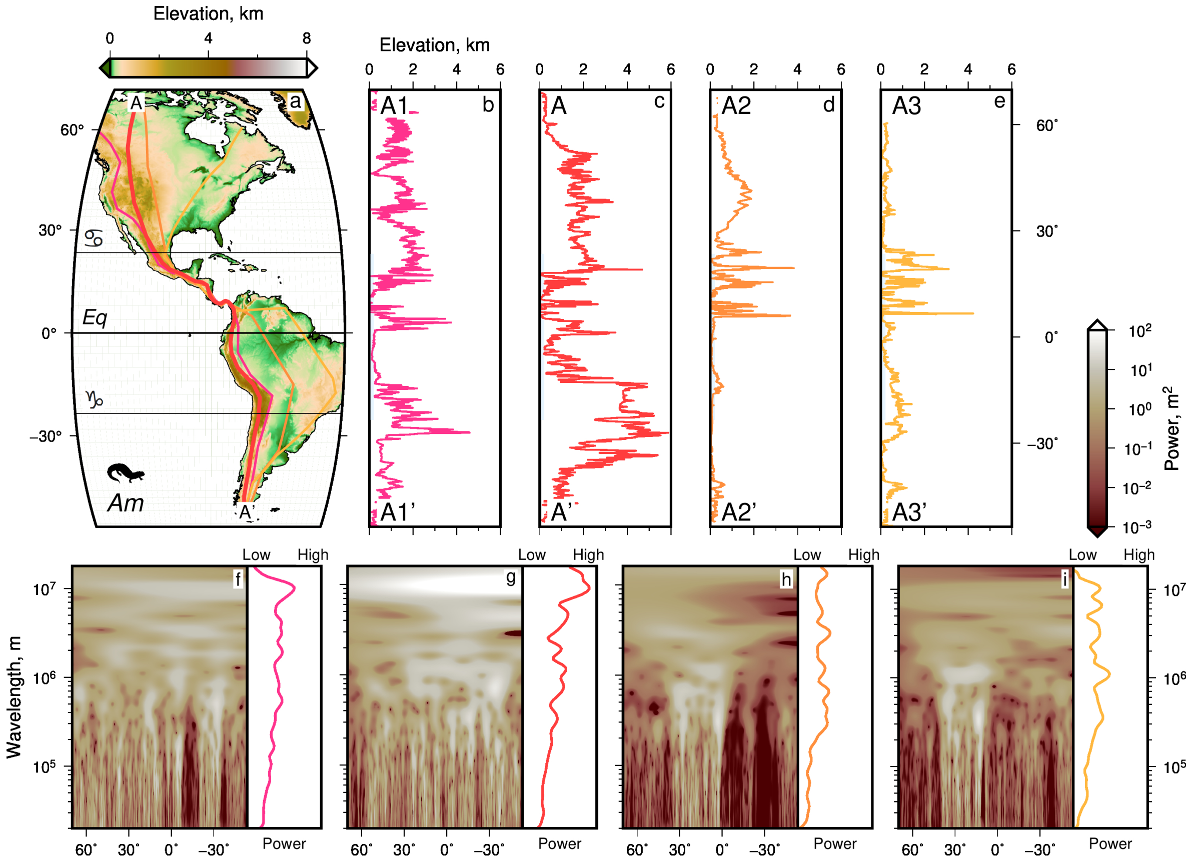


**Figure 37: Analysis of alternative transects through elevation across the Americas**. As Figure 36 but for elevation instead of species richness (Amante & Eakins, 2009).

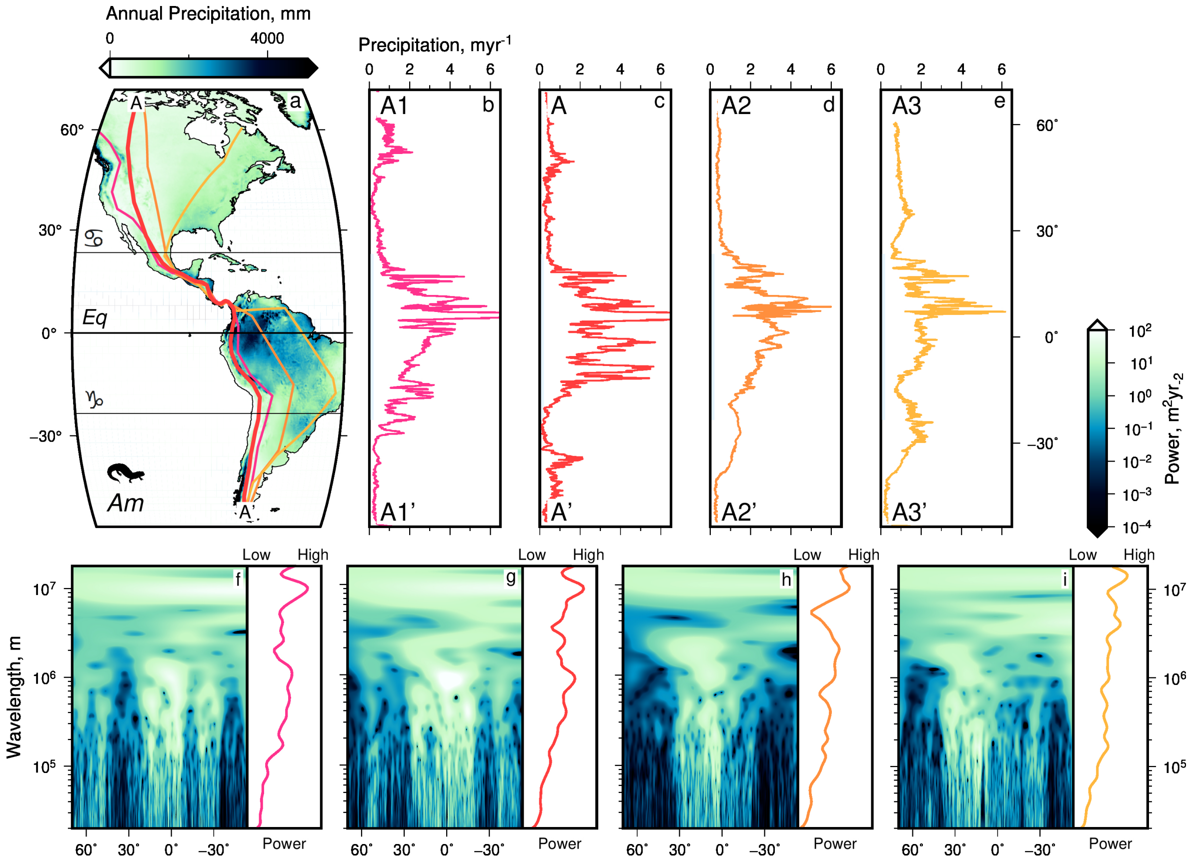


**Figure 38: Analysis of alternative transects through annual precipitation across the Americas**. As Figure 36 but for mean annual precipitation instead of species richness (Karger *et al.*, 2017).

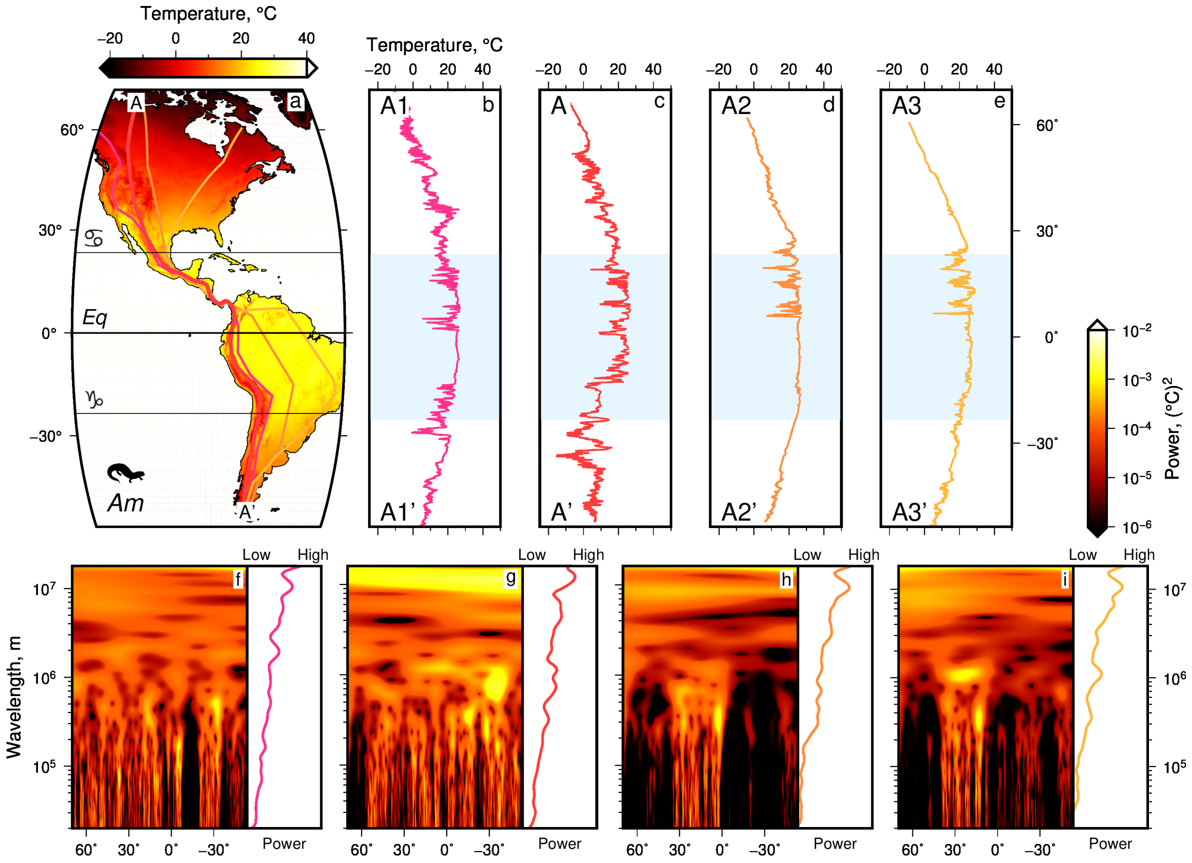


**Figure 39: Analysis of alternative transects through mean temperature across the Americas**. As Figure 36 but for mean temperature instead of species richness (Karger *et al.*, 2017).

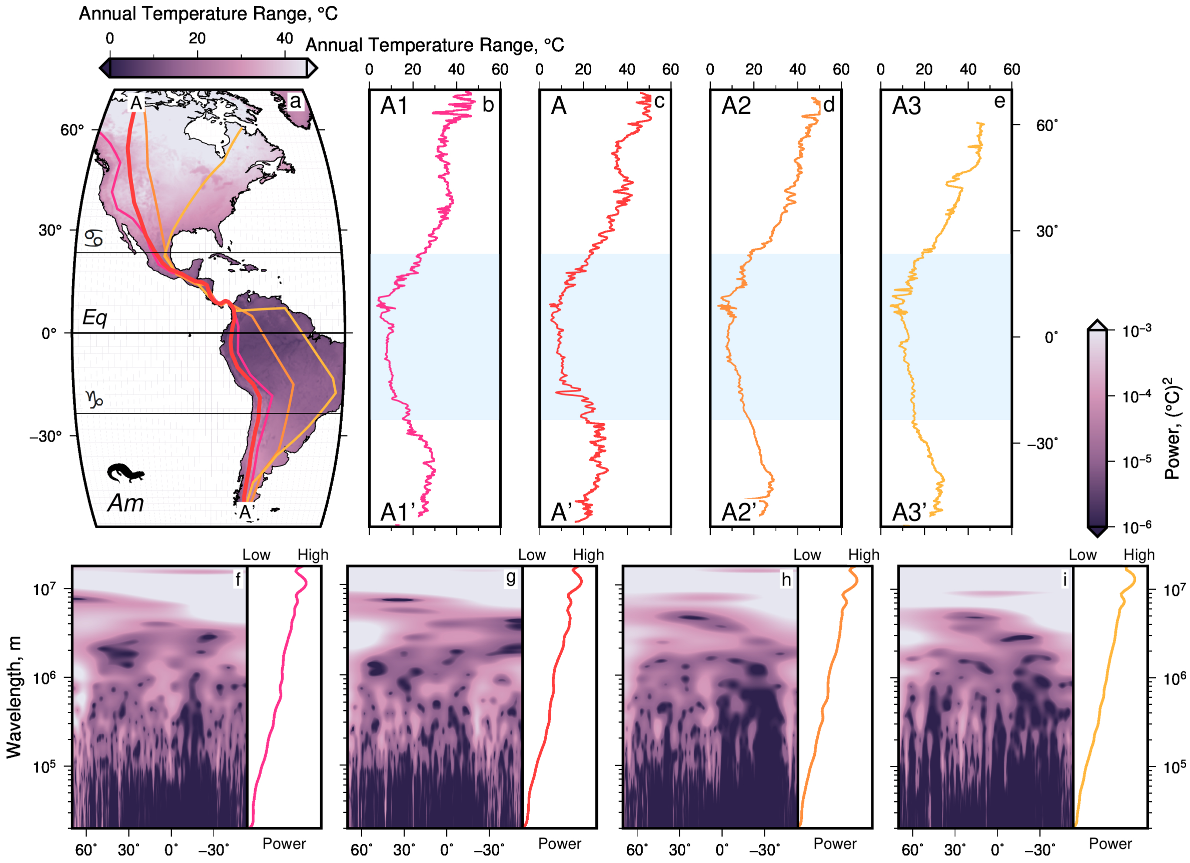


**Figure 40: Analysis of alternative transects through annual temperature range across the Americas**. As Figure 36 but for annual temperature range instead of species richness (Karger *et al.*, 2017).

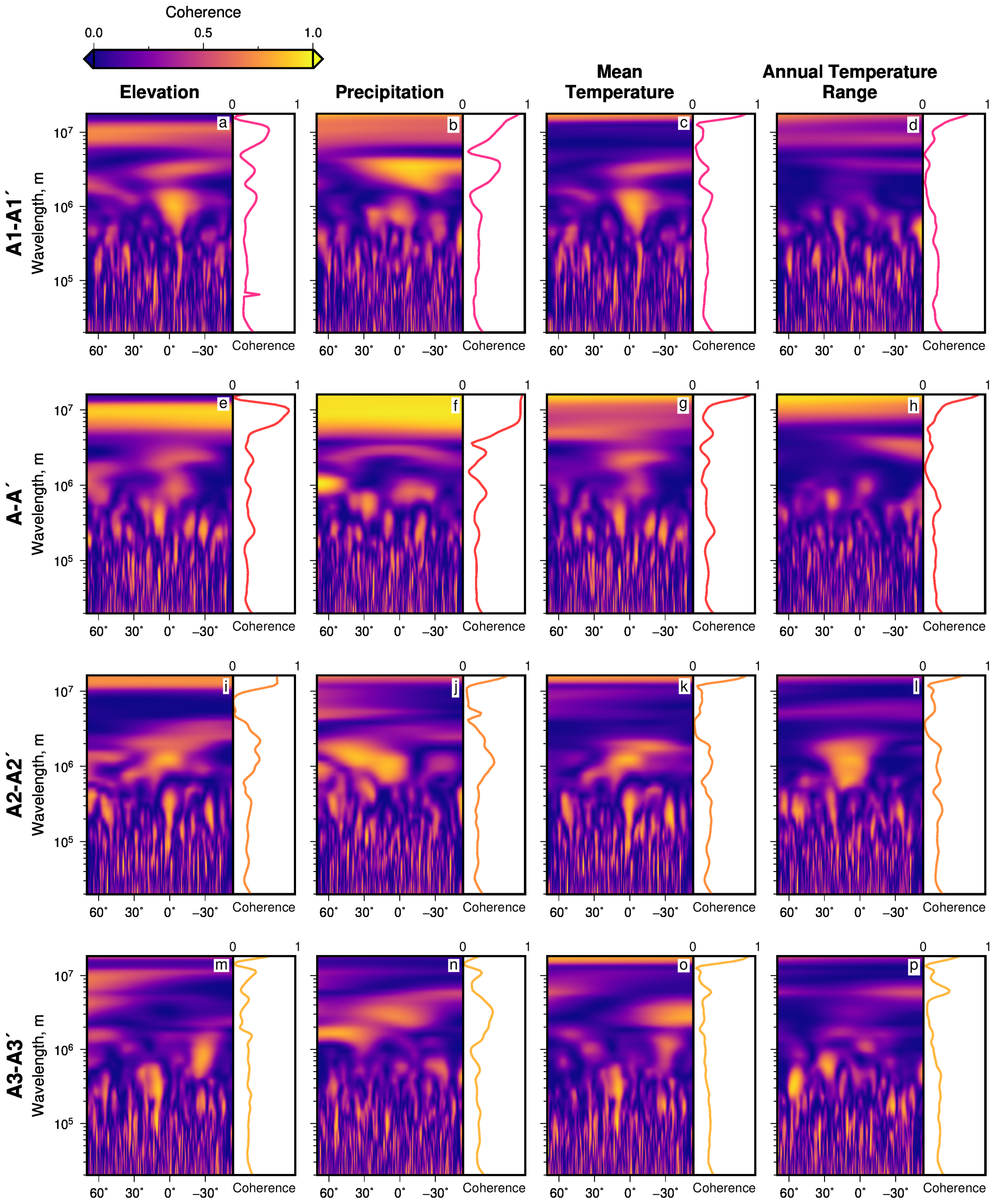


**Figure 41: Coherence between amphibian species richness and environment, along alternative American transects**. (a) Coherence between amphibian species richness and elevation, along westernmost transect in Figure 36, A1—A1′, as a function of latitude and scale. Side panel = mean coherence across all latitudes at given scale. (b)–(d) As (a) but for coherence between amphibian species richness and annual precipitation, mean temperature, and annual temperature range respectively. (e)–(h) As (a)–(d) but for transect A—A′ (see Figure 2 of main text). (i)–(l) As (a)–(d) but for transect A2—A2′ (see Figure 36. (m)–(p) As (a)–(d) but for easternmost transect, A3—A3′.
